# Neuroanatomical Correlates of Negative Symptoms in Schizophrenia

**DOI:** 10.1101/2025.09.22.677864

**Authors:** SO M Vijayakumar Kamalakannan, Alie G. Male, Musa Yilanli, Annalisa Lella, Jaylen Lee, Yann Quidé, Melissa J. Green, Murray J. Cairns, Vaughan J. Carr, Stanley Catts, Frans A. Henskens, Assen Jablensky, Carmel Loughland, Patricia Michie, Bryan Mowry, Christos Pantelis, Paul Rasser, Ulrich Shall, Rodney J. Scott, Thomas W. Weickert, Ayse Belger, Juan Bustillo, Kelvin Lim, Judith M. Ford, Daniel H. Mathalon, Adrian Preda, Bryon Mueller, Steven G. Potkin, Theodore D. Satterhwaite, Ruben C. Gur, Raquel E. Gur, Nerisa Banaj, Daniela Vecchio, Fabrizio Piras, Federica Piras, Stefan Ehrlich, Fabio Bernardoni, Stefan Borgwardt, Derin Cobia, Kate Alpert, Lei Wang, Ingrid Agartz, Erik G. Jönsson, Stefan Kaiser, Edith Pomarol-Clotet, Raymond Salvador, Carlos López-Jaramillo, Ana M. Diaz-Zuluaga, Julian Pineda-Zapata, Tilo Kircher, Frederike Stein, Axel Krug, Udo Dannlowski, Dominik Grotegerd, Jan-Bernard Marsman, André Aleman, Anthony O. Ahmed, Gregory P. Strauss, Paul A. Thompson, Matthias Krischner, Vince D. Calhoun, Jessica A. Turner, Theo G.M. van Erp

## Abstract

**Background:** Schizophrenia is characterized by widespread structural brain abnormalities, but associations between structural abnormalities and negative symptom severity are not well understood. Negative symptoms have been conceptualized in a hierarchical structure of two second-order dimensions—motivation and pleasure (MAP) and expression (EXP)—and five first-order domains: anhedonia, avolition, and asociality (MAP), and blunted affect and alogia (EXP). A better understanding of the neural circuitry underlying negative symptom dimensions and domains is important given their reported association with poor functional outcome and lack of available treatments.

**Study Design:** The meta-analysis included 1,591 individuals with schizophrenia across 16 samples with structural imaging and Scale for Assessment of Negative Symptoms data. The study generated correlations of cortical thickness and subcortical volumes with the negative symptom dimensions and domains.

**Study results:** Negative symptoms showed mainly negative associations with cortical thickness and subcortical volumes. The effect sizes were small but there was a pattern of associations in predominantly frontal lobe cortical thickness and limbic subcortical volumes. The regional correlation patterns of cortical thickness and subcortical volumes with symptom domains support the conceptualized hierarchical structure of negative symptoms: correlations of MAP domains were stronger with the MAP than EXP dimension, and vice versa. Exploratory analyses with receptor densities further supported the hierarchy.

**Conclusion:** Our findings reveal small but consistent associations between negative symptom dimensions and predominantly prefrontal region cortical thickness, and limbic region volumes.

These findings advance our understanding of the network of anatomical regions that may contribute to the severity of negative symptoms in schizophrenia.

## 1. Introduction

Schizophrenia is associated with replicable structural brain abnormalities^1–6^ but, likely in part due to insufficiently powered studies and other methodological issues^7,8^, their relationships with symptoms remain not fully determined. A better understanding of neural circuitry underlying negative symptoms is relevant given reported associations with poor functional outcome^9–11^ and lack of adequate treatments^12^.

The NIMH-MATRICS Consensus Statement on Negative Symptoms recognized five negative symptom domains, i.e., avolition, anhedonia, asociality, alogia, and blunted affect, and suggested they may map onto separate neurobiological substrates that could represent separate therapeutic targets^13^. Confirmatory factor analyses^14–21^ and a network analysis^22^ have confirmed a hierarchical structure with two second-order dimensions [motivation and pleasure (MAP) and expression (EXP)] and five first-order domains (anhedonia, avolition, and asociality part of the MAP dimension, and blunted affect and alogia part of the EXP dimension) corroborating the consensus statement^14,22,23^. Strauss and colleagues (2018)^14^ reported an initial mapping of the negative symptom domains to the Research Domain Criteria (RDoC) matrix^23^, though this mapping remains to be validated.

Relationships between brain structural abnormalities and negative symptoms have been mapped in several ways. First, studies have compared individuals with (deficit schizophrenia; DSZ^24,25^) or without primary and enduring negative symptom schizophrenia (non-deficit schizophrenia; NDSZ) and healthy controls^26–31^. Our prior meta-analysis found that, compared to controls, DSZ has a unique pattern of thinner left prefrontal and temporal cortex compared to NDSZ^27^. Second, studies have examined relationships between brain structure metrics and total negative symptom severity^3,32–34^. Our prior meta-analysis replicated earlier findings^33^ that left medial orbital frontal cortex (mOFC) thickness was significantly associated with total negative symptom severity^32^. Our subsequent study found significant relationships between medial and lateral frontal (including the mOFC), and also lateral temporal and parietal cortical thickness and total negative symptoms^3^. No significant associations between overall negative symptom severity and cortical surface area were found, suggesting that relationships between brain structure and negative symptoms may be restricted to gray matter. Finally, to our knowledge, no studies have examined relationships between structural brain abnormalities and the EXP and MAP dimensions or anhedonia, avolition, asociality, blunted affect and alogia subdomain factor scores, which may arguably map better to neurocircuitry than summary or global symptom ratings^35^; for review, see^8^).

This study examines relationships between negative symptoms based on the Scale for the Assessment of Negative Symptoms (SANS) and deep brain structure (subcortical) volumes and cortical thickness in individuals with schizophrenia. This enables refinement of the mapping between negative symptom dimensions/domains and brain structure based on the RDoC framework. This study implemented and tested a COINSTAC (Collaborative Informatics and Neuroimaging Suite Toolkit for Anonymous Computation^36^) analysis pipeline, generating a published and validated federated analysis for future replication studies.

Based on the proposed mapping of negative symptoms dimensions (MAP, EXP) and negative symptom domains (anhedonia, avolition, and asociality, blunted affect, alogia) to the RDoC by Strauss and colleagues^23^, and our prior findings of relationships between cortical thickness and total negative symptoms, we hypothesized that (i) anhedonia is negatively associated with anterior insula, dorsal anterior cingulate cortex (dACC), medial orbitofrontal (mOFC), and ventromedial prefrontal cortex (vmPFC) thickness, and pallidum (P) and nucleus accumbens (NAcc) volumes; (ii) asociality is negatively associated with fusiform face area and orbitofrontal cortex (OFC) thickness and amygdala, nucleus accumbens (NAcc), and pallidum (P) volumes; (iii) avolition is negatively associated with orbitofrontal (OFC) and medial prefrontal cortex (medial PFC) thickness, and amygdala, nucleus accumbens (NAcc), and ventral pallidum (VP) volumes; and (iv) blunted affect and (v) alogia are negatively associated with superior temporal gyrus (STG) thickness. An exploratory analysis examined associations between published regional receptor densities^37^ and this meta-analysis’ regional correlation maps of cortical thickness with negative symptom severity.

## 2. Methods

### 2.1. Study Samples

Sixteen cross-sectional study samples, totaling 1,591 individuals with schizophrenia (SZ) assessed with high-resolution structural brain scans and the SANS^38^ (Scale for the Assessment of Negative Symptoms), contributed to the analysis via the ENIGMA Schizophrenia Working Group (**Supplementary Tables 1-2**). Sample-size weighted mean (range) age across samples was 36.7 (25.7-42.9) years and samples were on average 69% male (based on self-reported sex).

Weighted mean age at onset and duration of illness across the samples were 22.8 (19.1-26.7) and 13.7 (1.1-18.8) years. Weighted mean PANSS^39^ (Positive and Negative Syndrome Scale) total, negative, and positive scores across the samples were 66.7 (48.6-86.3), 17.4 (14.5-23.5), and 16.2 (10.7-20.9); weighted mean SANS and SAPS^40^ (Scale for the Assessment of Positive Symptoms) scores were 22.6 (6.9-40.1) and 19.7 (6.4-31.5). For samples that recorded current antipsychotic type and/or dose, numbers (percentages) of patients on second-generation (atypical), first-generation (typical), both, or none, were 975 (68%), 152 (11%), 106 (7%), and 207 (14%), respectively. Sample-size weighted mean chlorpromazine dose equivalent (CPZ), based on Woods^41^, was 405 (209-655). Each study sample was collected with participants’ written informed consent approved by local Institutional Review Boards.

### 2.2. Image Acquisition and Processing

Each study collected high-resolution T1-weighted structural magnetic resonance imaging (MRI) brain scans (see **Supplementary Table 1** for details on scanners, vendors, field strengths, sequences, and acquisition parameters). ENIGMA’s quality assurance protocol was performed at each site prior to analysis and included visual checks of the cortical segmentations and region-by-region removal of values for segmentations found to be incorrect (http://enigma.usc.edu/protocols/imaging-protocols).

Histograms of all regions’ values for each site were also computed for visual inspection.

All sites processed T1-weighted structural brain scans using FreeSurfer^42^ (http://surfer.nmr.mgh.harvard.edu; see versions used in **Supplementary Table 2**) and extracted deep brain structure volumes (accumbens, amygdala, caudate, hippocampus, pallidum, putamen, thalamus, and lateral ventricle), and cortical thickness for 70 Desikan-Killiany^43^ (DK) atlas regions (34 regions per hemisphere + left and right hemisphere mean thickness).

### 2.3. Negative Symptom Severity Measures

In addition to total negative symptom severity (SANS Total), negative symptom domain factor scores for 2 factor (MAP and EXP) and 5 factor (anhedonia, asociality, avolition, blunted affect, alogia) models^23,44^ were calculated based on Scale for the Assessment of Negative Symptoms (SANS^45^) ratings. We used equations based on confirmatory factor analyses (Strauss & Ahmed, personal communication, January 6, 2020; see **Supplemental Information** for details on the factor score computations). Males had higher negative symptom severity than females for each of the measures [range *d* =-0.23 to-0.13]. (see **Supplementary Table 3**).

### 2.4. Statistical Meta-analyses

Within each sample, associations between each symptom measure and mean cortical thickness in DK atlas regions of interest, and mean subcortical volumes were examined using univariate linear regression (R’s linear model function lm). Negative symptom measures were regressed onto structural brain measures co-varying for sex (self-reported) and age, for the cortical thickness analyses, and sex, age, and intracranial volume for the deep brain structure volumes.

Exploratory analysis then examined associations between each negative symptom measure, and left or right hemisphere regions. Analysis of multi-scanner studies (ASRB, FBIRN, MCIC, Osaka, UPENN) included binary dummy covariates for n-1 scanners. Additional analyses statistically controlled for possible confounding variables by adding medication type or medication dose as covariates to the model.

All 16 sites participated in the main meta-analysis, though the subcortical results from SCORE and the cortical results from FOR2107-MS were corrupted and not included in the main meta-analyses resulting in meta-analyses of data from 15 sites for both the cortical and subcortical analyses. Six sites also successfully participated in the analysis based on the statistical analysis pipeline implemented in the Collaborative Informatics and Neuroimaging Suite Toolkit for Anonymous Computation (COINSTAC)^36^.

Effect sizes (Pearson’s r-values) were transformed to Fisher *z*-correlations before random-effects meta-analyses of partial correlation effect sizes for each of the measures were performed using the R (version 3.2.2) metafor package (version 1.9-7)^46^. Variances were estimated using ML (maximum likelihood) and statistical significance tested using *t*-tests. False Discovery Rate (*p*_FDR_<0.05)^47^ was used to adjust for multiple comparisons.

Between-symptom Spearman’s correlations of relationships between effect size (z-correlation) patterns for cortical thickness and subcortical volumes across symptom dimensions and domains were examined. Finally, to identify possible involvement of different neurobiological targets (receptors) between negative symptom dimensions and domains, spin tests^48,49^ examined relationships between this meta-analysis’ regional effect size (correlation) patterns for cortical thickness and published receptor distributions^37^.

## 3. Results

### 3.1. Associations Between Structural Abnormalities and Negative Symptom Severity

Overall, the associations between brain structure and symptoms were mostly negative and small [range *r*_cortical thickness_ =-0.11 to 0.06; range *r*_subcortical volumes_ =-0.10 to 0.09]. In addition to the few associations that survived FDR-correction, there were multiple nominally significant associations (*p* < 0.05 before FDR correction). For brevity, we only describe those relevant to our hypotheses in the text accompanied by complete results in tables and figures. In the models with medication type or medication dose included as covariates, effect sizes were similar, but fewer associations remained FDR-level significant (see **Supplementary Table 4**).

#### 3.1.1. Associations with Cortical Thickness

##### 3.1.1.1. SANS Total

SANS Total was significantly associated with cortical thickness of mean superior frontal gyrus (*r*_1325_=-0.09, 95%CI [-0.136--0.051], *p_FDR_*=0.012). Exploratory analysis by hemisphere (left, right) showed significant associations with the left (*r*_1350_=-0.08, 95%CI [-0.127--0.041], *p*_FDR_=0.031) and right (*r*_1341_=-0.10, 95%CI[-0.141--0.051], *p*_FDR_=0.030) superior frontal gyri. There was also a significant association with the left pars orbitalis gyrus (*r*_1327_=-0.08, 95% CI [-0.12--0.035], *p_FDR_*=0.035). Apart from the associations that remained significant after FDR correction, there were 16 nominally significant associations with SANS Total (15 negative and 1 positive; 7 for mean (see **Figure 1** and **2**), and 7 for left and 2 for right hemisphere; see **Supplementary Table 5**).

**Figure 1.**
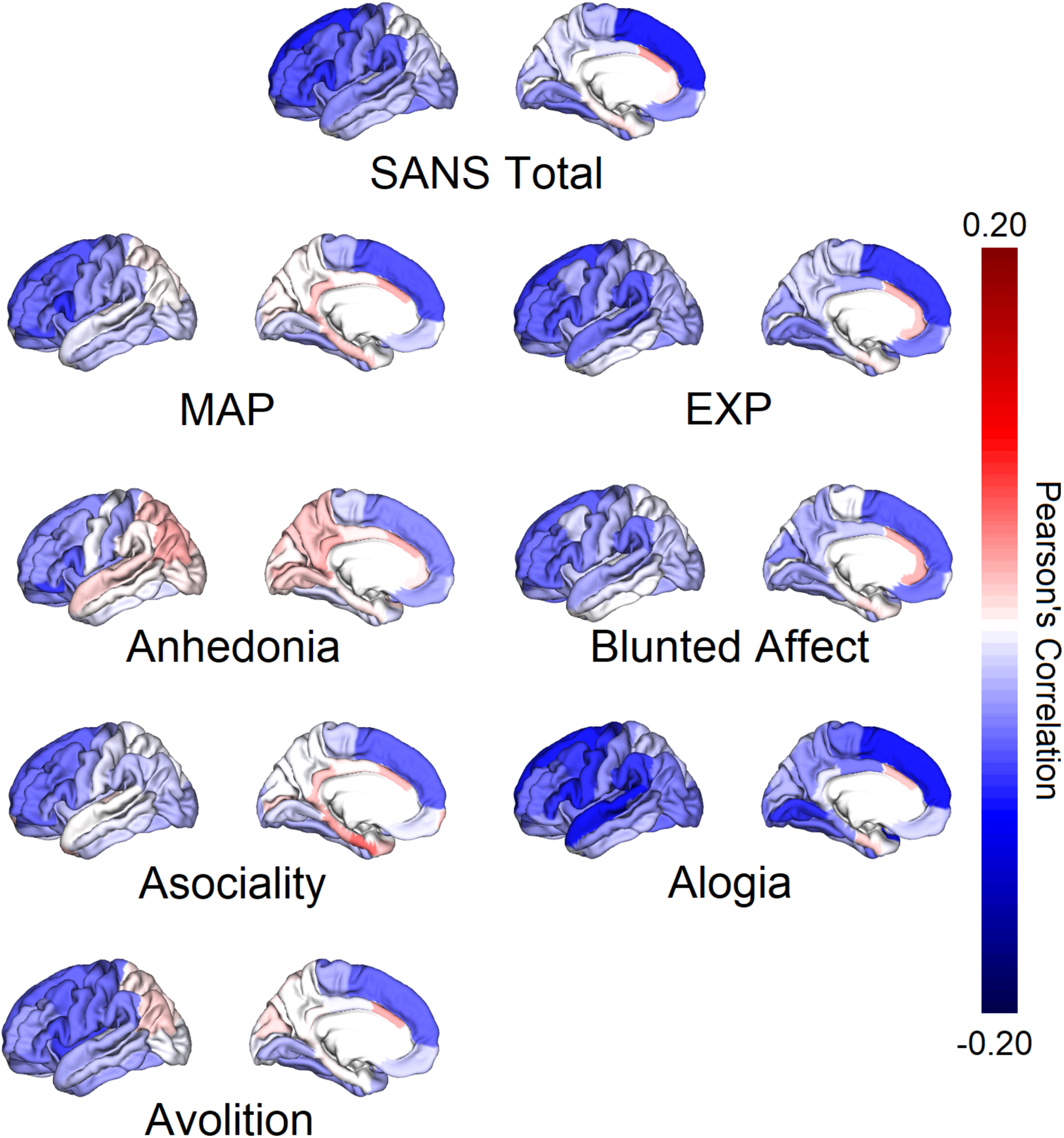
Correlations between Mean Cortical Thickness and Negative Symptom Factors. Note: FDR-corrected* and nominally significant correlations between cortical thickness were found between: ***SANS Total*** and mean Superior Frontal*, Caudal Middle Frontal, Pars Opercularis, Pars Orbitalis, Pars Triangularis, Precentral, Rostral Middle Frontal and Supramarginal; ***MAP*** and mean Caudal Middle Frontal, Pars Orbitalis, Rostral Middle Frontal, Superior Frontal and Insula; *Anhedonia* and Mean Pars Orbitalis; *Asociality* and Mean Pars Orbitalis, Rostral Middle Frontal and Superior Frontal; *Avolition* and Mean Caudal Middle Frontal, Pars Orbitalis, Pars Triangularis, Precentral, Mean Superior Frontal and Mean Insula; ***EXP*** and mean Pars Opercularis, Pars Orbitalis, Pars Triangularis, Precentral, Rostral Middle Frontal, Superior Frontal and Supramarginal; *Blunted Affect* and Mean Pars Opercularis, Pars Orbitalis, Pars Triangularis, Rostral Middle Frontal and Superior Frontal; *Alogia* and Mean Caudal Middle Frontal*, Precentral*, Superior Frontal*, Superior Temporal*, Lateral Occipital, Lingual, Paracentral, Pars Opercularis, Pars Triangularis, Posterior Cingulate, Rostral Middle Frontal and Supramarginal gyri.

**Figure 2.**
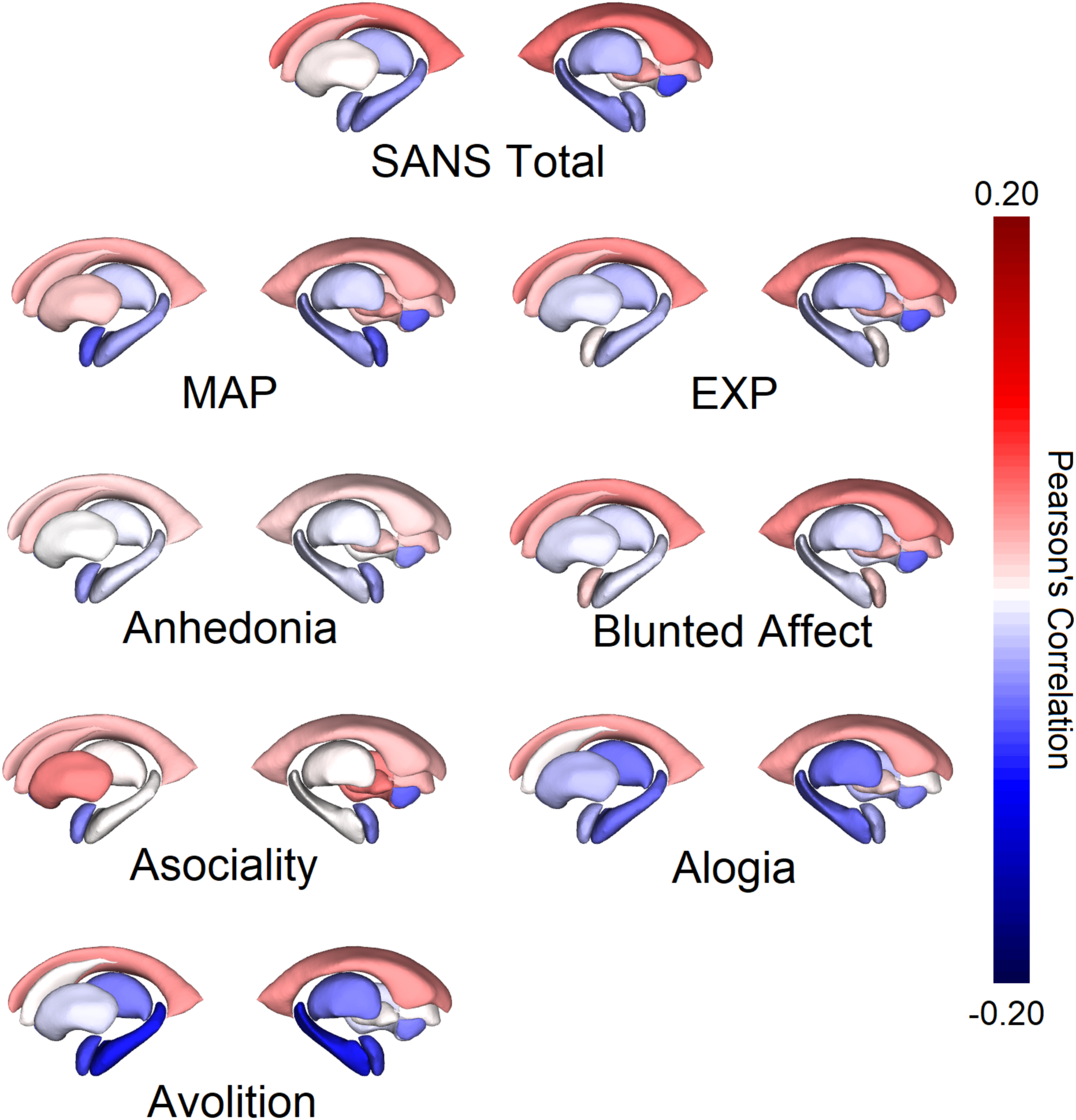
Correlations between Mean Deep Brain Structure Volumes and Negative Symptom Factors. Note: FDR-corrected* and nominally significant correlations between cortical thickness were found between: ***SANS Total*** and mean Lateral Ventricle and Accumbens. ***MAP*** and mean Accumbens; *Asociality* and Accumbens; *Avolition* and mean Thalamus, Hippocampus* and Amygdala. ***EXP*** and mean Accumbens.

##### 3.1.1.2. MAP Dimension and Domains

There were no associations between MAP and cortical thickness that remained significant after FDR correction. There were, however, 15 nominally significant negative associations with MAP (5 for mean (see **Figure 1**), and 5 for left and right hemisphere each; see **Supplementary Table 6)**.

With the exception of *avolition*, none of the associations between the other MAP subdomains (i.e., anhedonia and asociality) and cortical thickness remained significant after FDR correction (**Figure 1** and **Supplementary Tables 7-9**).

*Avolition* was not significantly associated with mean regional cortical thickness after FDR correction, but was significantly associated with left hemisphere pars triangularis (*r*_1304_=-0.09, 95% CI [-0.127--0.045], *p*_FDR_=0.027) and right hemisphere caudal middle frontal (*r*_1337_=-0.08, 95% CI [-0.116--0.039], *p*_FDR_=0.027) gyri. Furthermore, in support of our hypotheses, *avolition* was also nominally associated with mean (*r*_1301_=-0.06, 95% CI [-0.115--0.012], *p*= 0.006) and left hemisphere pars orbitalis (*r*_1327_=-0.05, 95% CI [-0.088--0.003], *p*=0.038), mean (*r*_1325_=-0.07, 95% CI,*p*=0.018) and right hemisphere superior frontal (*r*_1341_=-0.07, 95% CI [-0.118--0.019], *p*=0.01), and left lateral orbitofrontal (*r*_1364_=-0.04, 95% CI [-0.087-0.00], *p*=0.048) gyri.

There were 6 nominally significant associations with *anhedonia* (1 for mean, 4 for left and 1 for right hemisphere respectively), 6 with *asociality* (3 for mean, and 3 for right hemisphere), and 12 nominally significant negative associations – including those hypothesized reported above – with avolition (6 for mean, and 5 for left and 1 for right hemisphere). Of note, in support of our hypotheses, *asociality* was nominally associated with mean pars orbitalis thickness (*r*_1301_=-0.06, 95% CI [-0.111--0.017], *p*=0.012).

##### 3.1.1.3. EXP Dimension and Domains

There were no associations between EXP and cortical thickness that remained significant after FDR correction. However, there were 17 nominally significant negative associations with EXP (7 for mean (see **Figure 1**), and 7 for left and 3 for right hemisphere; see **Supplementary Table S10**).

Of the EXP subdomains (see **Figure 1**), none of the associations between *blunted affect* and cortical thickness were significant after FDR correction. However, there were 11 nominally significant (9 negative and 2 positive; 5 for mean, 4 for left and 2 for right hemisphere) associations for *blunted affect* (see **Supplementary Table 11**).

In support of our hypothesis, *alogia* was significantly associated with mean superior temporal gyrus (*r*_1132_=-0.11, 95% CI [-0.164--0.046], *p_FDR_*= 0.034). Furthermore, there were significant associations with mean caudal middle frontal (*r*_1323_=-0.08, 95% CI [-0.135--0.029], *p_FDR_*=0.043), precentral (*r*_1318_=-0.09, 95% CI [-0.144--0.031], *p_FDR_*=0.043), superior frontal (*r*_1325_=-0.10, 95% CI [-0.139--0.056], *p_FDR_*=0.007), and superior temporal (*r*_1132_=-0.11, 95% CI [-0.164--0.046], *p_FDR_*=0.034) gyri. Exploratory analyses by hemisphere showed significant associations between *alogia* and right hemisphere caudal middle frontal (*r*_1337_=-0.10, 95% CI [-0.156--0.041], *p_FDR_*=0.044), and left (*r*_1350_=-0.10, 95% CI [-0.144--0.058], *p_FDR_*=0.014) and right (*r*_1341_=-0.09, 95% CI [-0.134--0.047], *p_FDR_*=0.019) hemisphere superior frontal gyri. There was also a significant association with left rostral middle frontal gyrus (*r*_1312_=-0.08, 95% CI [-0.125--0.035], *p_FDR_*=0.044). Furthermore, there were 25 nominally significant negative associations with *alogia* (8 for mean, and 10 for left and 7 for right hemisphere; see **Supplementary Table 12**).

#### 3.1.2. Associations with Subcortical Volumes

##### 3.1.2.1. SANS Total

While there were no significant associations between SANS Total and subcortical volumes after FDR correction, there were 4 nominally significant associations (1 negative and 3 positive; 2 for mean (see **Figure 2**), 1 for left and 1 for right hemisphere respectively; see **Supplementary Table 13**).

##### 3.1.2.2. MAP Dimension and Domains

While there were no associations between MAP and subcortical volumes that remained significant after FDR correction, there were 2 nominally significant negative associations (1 for mean (see **Figure 2**), and 1 for right hemisphere; see **Supplementary Table 14**). As with cortical thickness, there were no associations that remained significant after FDR correction among the MAP domains with the exception of avolition (**Figure 2** and **Supplementary Tables 14-16**). While there were none for *anhedonia*, there were 2 nominally significant associations between *asociality* and subcortical volumes. Specifically, in support of our hypothesis, *asociality* was nominally significantly associated with mean (*r*_1318_=-0.07, 95% CI [-0.126--0.005], *p*=0.037) and right hemisphere (*r*_1335_=-0.05, 95% CI [-0.09--0.005], *p*=0.032) nucleus accumbens volumes.

In contrast, *avolition* was significantly associated with mean hippocampus volumes (*r*_1335_=-0.10, 95% CI [-0.141--0.051], *p_FDR_*=0.004) and exploratory analysis by hemisphere revealed that *avolition* was significantly associated with both left (*r*_1339_=-0.09, 95% CI [-0.138--0.045], *p_FDR_*=0.007) and right (*r*_1349_=-0.10, 95% CI [-0.142--0.048], *p_FDR_*=0.007) hippocampus volumes. There were also 5 nominally significant associations for *avolition* (2 for mean, 1 for left and 2 for right hemispheres respectively). Of note, in line with our hypothesis, there were nominally significant associations between *avolition*, and mean (*r*_1341_=-0.08, 95% CI [-0.139--0.018], *p*=0.015) and left (*r*_1345_=-0.06, 95% CI [-0.114--0.015], *p*=0.015) amygdala volumes).

##### 3.1.2.3. EXP Dimension and Domains

There were no associations that remained significant after FDR correction between the EXP dimension (see **Figures 2** and **Supplementary Table 18**), nor the constituent domains of *blunted affect* or *alogia* (see **Supplementary Tables 19-20**), and subcortical volumes. While there were also no nominally significant associations between the EXP dimension or domain of blunted affect and any of the subcortical volumes, there was 1 nominally significant association between alogia and right hippocampus.

### 3.2. COINSTAC Analysis

Six samples successfully participated in the COINSTAC analysis (FBIRN, FIDMAG, MCIC, UCISZ, UMCG, and ZURICH). Direct comparison of the COINSTAC analysis with the traditional R meta-analysis from these six sites found identical results, supporting the use of the decentralized analysis platform (**Supplementary Figure 3**).

### 3.3. Relationships between Symptom Association Effect Size Maps

The relationships of MAP dimension with MAP domain (anhedonia, asociality, and avolition) factor scores showed higher correlations for cortical thickness (*r*=0.85, 0.91, and 0.91; **Figure 3a**) and subcortical volumes (*r*=0.90, 0.88, and 0.84; **Figure 3b**) than with EXP domain (blunted affect, and alogia) factor scores (*r*=0.62 and 0.59; *r*=0.65 and 0.76, respectively). Likewise, the relationships of EXP dimension with EXP domain (blunted affect, and alogia) factor scores showed higher correlations for cortical thickness (*r*=0.96 and 0.74; see **Figure 3a**) and subcortical volumes (*r*=0.97 and 0.88; see **Figure 3b**) than with MAP domain (anhedonia, asociality, and avolition) factor scores (*r*=0.49, 0.71, and 0.59; *r*=0.74, 0.62, and 0.71, respectively). Steiger’s Z-tests (R’s cocor package^50,51^) showed that the correlation for blunted affect with EXP was significantly higher than for alogia with EXP (*Z* = 4.55, *p* <0.001); the correlations for the other domains did not differ from each other.

**Figure 3a.**
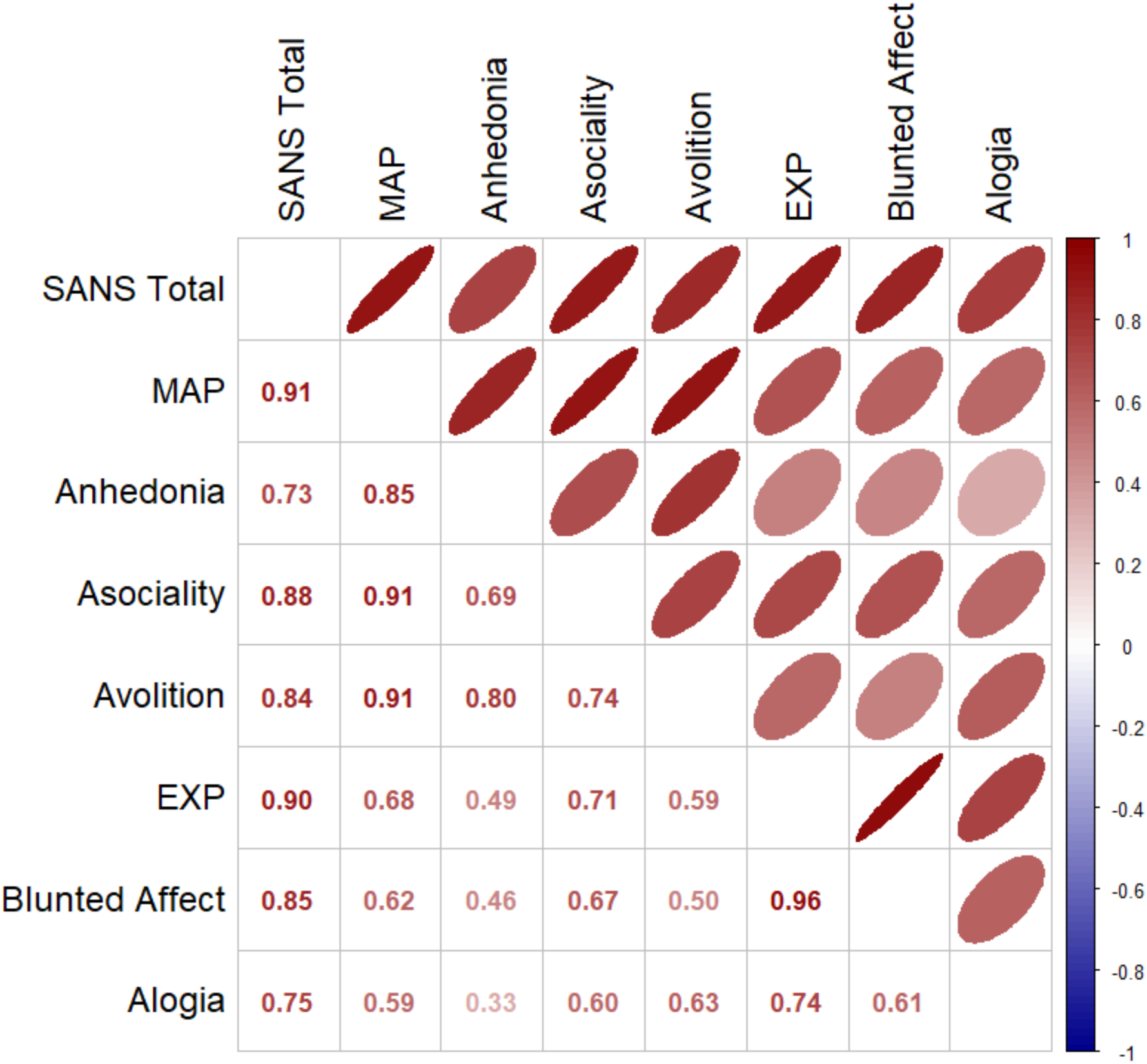
Between Symptom Spearman Rank Correlations of Effect Size Patterns across Mean Cortical Thickness Regions

**Figure 3b.**
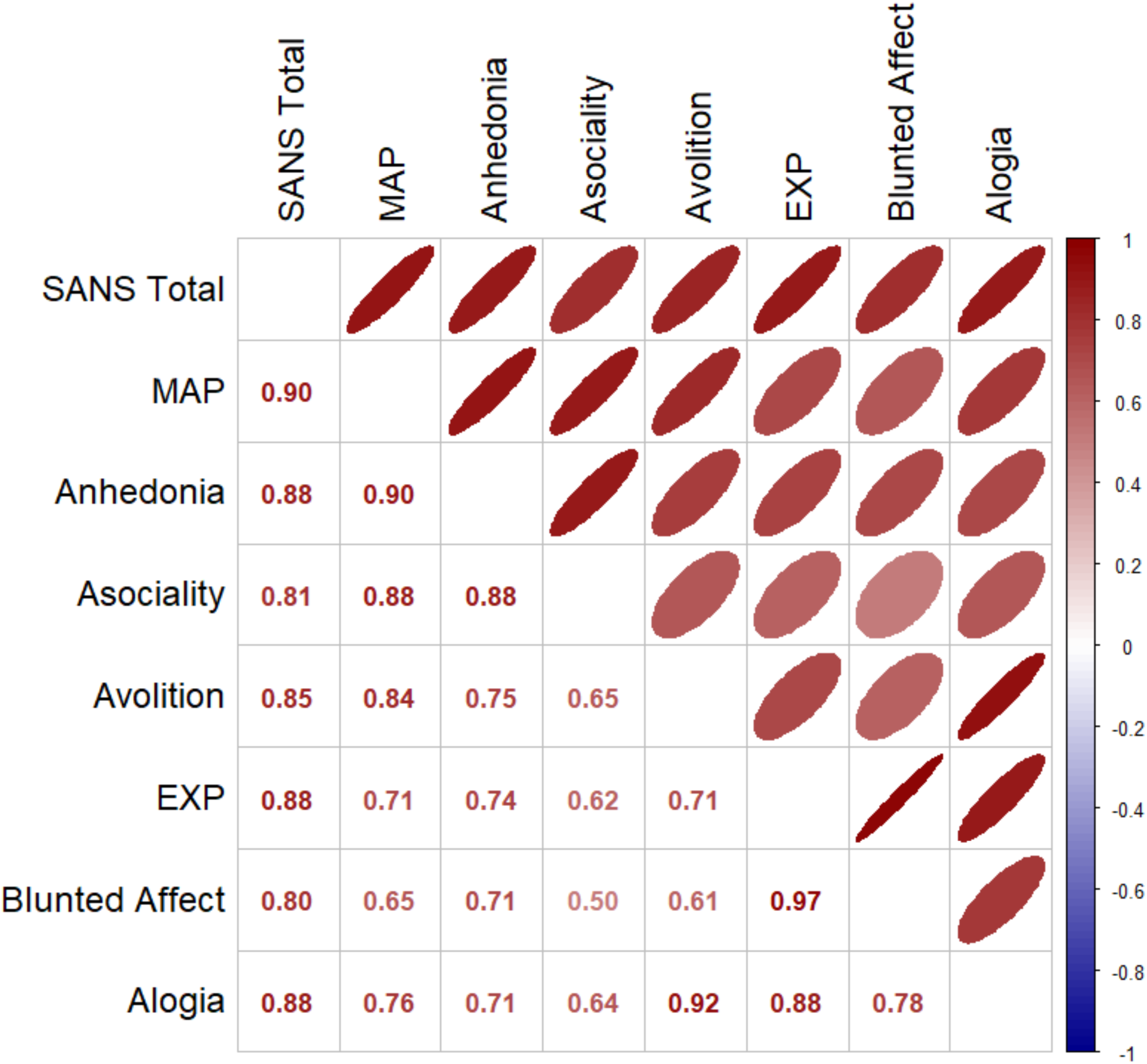
Between Symptom Spearman Rank Correlations of Effect Size Patterns across Mean Deep Brain Structure Regions

**Figure 4.**
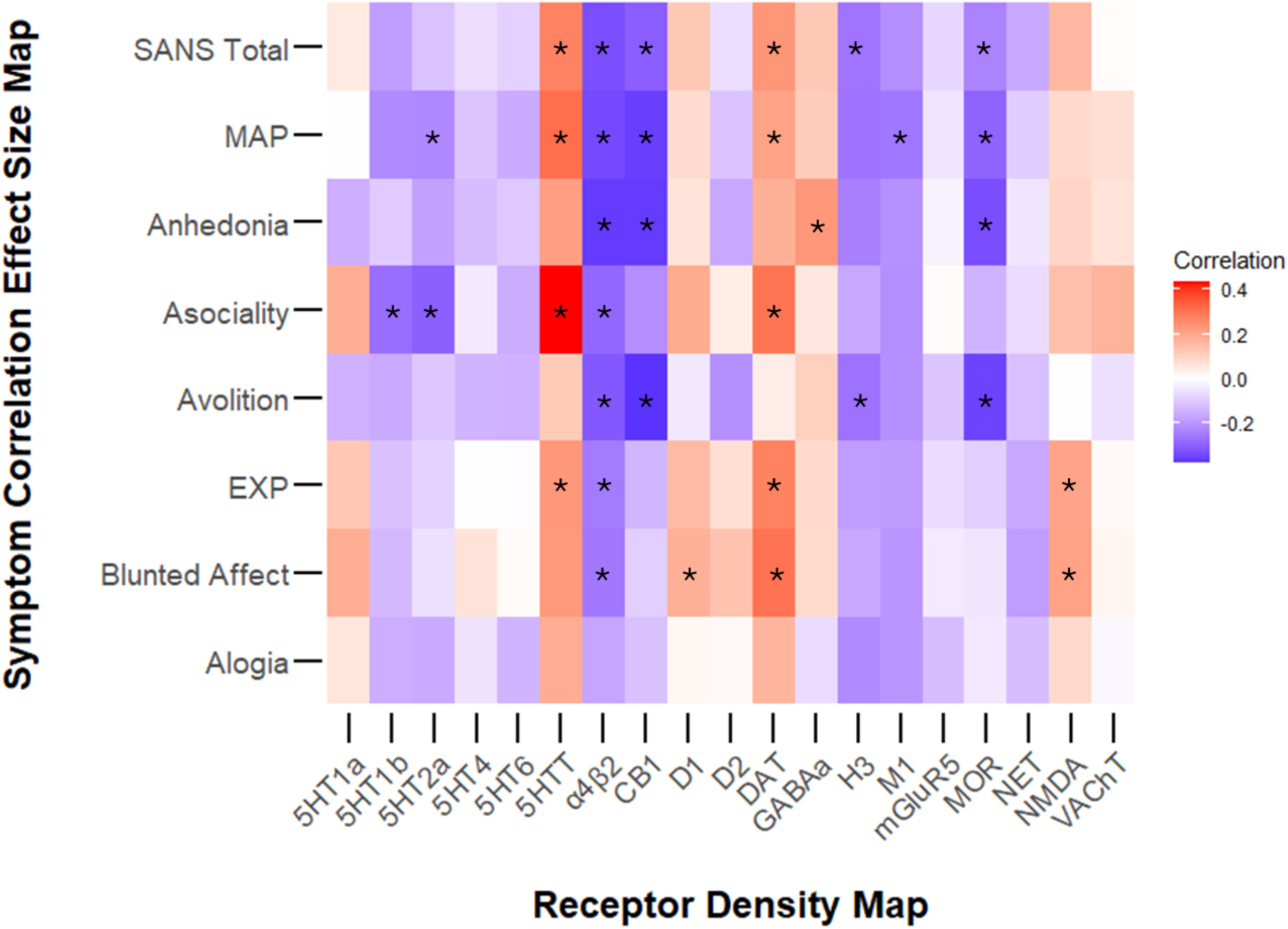
Heatmap of Associations Between Cortical Correlation Effect Size Maps and Receptor Distribution Maps. Note: * denotes significant associations

### 3.4 Correlations between Symptom Association Effect Size Maps and Receptor Distribution Maps

For SANS Total, higher regional receptor densities of A4B2 (*r*=-0.33, *p*=0.009), CB1 (*r*-0.30, *p*= 0.008), H3 (*r*=-0.26, *p*=0.033) and MOR (*r*-0.23, *p*=0.031) were correlated with stronger regional negative associations between SANS Total and cortical thickness, while lower densities of 5HTT (*r*= 0.27, *p*=0.018) and DAT (*r*=0.23, *p*=0.034) were correlated with stronger negative associations (see **Supplementary Table S21** for details).

For MAP, higher regional receptor densities of 5HT2a (*r*=-0.22, *p*=0.028), A4B2 (*r*=-0.34, *p*=0.009), CB1 (*r*=-0.36, *p*=0.004), M1 (*r*=-0.26, *p*=0.031) and MOR (*r*=-0.29, *p*=0.018) were correlated with stronger regional negative associations between MAP and cortical thickness, while lower densities of 5HTT (*r*= 0.31, *p*=0.002) and DAT (*r*=0.21, *p*=0.044) were correlated with stronger negative associations.

Among the MAP domains, for anhedonia, higher regional receptor densities of A4B2 (*r*=-0.36, *p*=0.006), CB1 (*r*=-0.36, *p*=0.001), and MOR (*r*=-0.33, *p*=0.005) were correlated with stronger regional negative associations between anhedonia and cortical thickness, while lower receptor densities of GABAa (*r*=0.23, *p*=0.041) were correlated with stronger negative associations. For asociality, higher regional receptor densities of 5HT1b (*r*=-0.27, *p*=0.021), 5HT2a (*r*=-0.30, *p*=0.005), A4B2 (*r*=-0.29, *p*=0.019), and CB1 (*r*=-0.21, *p*=0.0495) were correlated with stronger regional negative associations between asociality and cortical thickness, while lower densities of 5HTT (*r*=0.43, *p*<0.001), and DAT (*r*=0.30, *p*=0.007) were correlated with stronger negative associations. For avolition, higher regional receptor densities of A4B2 (*r*=-0.31, *p*=0.012), CB1 (*r*=-0.38, *p*=0.001), H3 (*r*=-0.26, *p*=0.041) and MOR (*r*=-0.35, *p*=0.004) were correlated with stronger regional negative associations between avolition and cortical thickness, while no lower receptor densities were correlated with stronger negative associations.

For the EXP dimension, higher regional receptor densities of A4B2 (*r*=-0.25, *p*=0.0415) were correlated with stronger regional negative associations between EXP and cortical thickness, while lower densities of 5HTT (*r*=0.23, *p*=0.044), DAT (*r*=0.28, *p*=0.013) and NMDA receptors (*r*=0.20, *p*=0.036) were correlated with stronger negative associations.

Among the EXP domains, for blunted affect, higher regional receptor densities of A4B2 (*r*=-0.25, *p*=0.042) were correlated with stronger regional negative associations between blunted affect and cortical thickness, while lower densities of 5HTT (*r*= 0.23, *p*=0.04), D1 (*r*=0.15, *p*=0.062), DAT (*r*=0.30, *p*=0.013), and NMDA receptors (*r*=0.20, *p*=0.036) were correlated with stronger negative associations. None of the receptor densities were significantly correlated with regional associations between alogia and cortical thickness.

## 4. Discussion

The principal findings of this study are that: 1) cortical thickness and subcortical volumes showed predominantly negative associations with negative symptoms; 2) correlation effect sizes were small, and few associations remained significant after FDR correction; 3) mainly frontal and limbic regions showed significant associations; 4) brain-symptom associations of MAP domains were stronger with the MAP than EXP dimension, and vice versa; 5) meta-analyses could be performed on decentralized data sets using COINSTAC; 6) patterns of cortical-symptom associations were related to distinct receptor density distributions.

In line with our hypotheses, we found several small predominantly negative (FDR or nominally) significant associations for mean superior frontal (SFG), pars orbitalis (IFGOrb), and rostral middle frontal (rMFG) cortical thickness, and mean NAcc volumes across SANS Total, MAP and EXP dimensions. However, after FDR-correction, only mean SFG remained significant for SANS Total. While of similar magnitude, there were fewer significant associations when medication type or dose were controlled for, likely reflective of reduced power in the smaller samples with both imaging and medication data.

This study replicated several of our previously reported associations between negative symptoms and thinner cortex in frontal regions^3,32^. However, the association between MOFC cortical thickness and total negative symptoms ^32^ did not replicate, possibly due to this study’s smaller sample size. Compared to our prior findings, there were also fewer associations of negative symptoms with temporal and parietal regions^3,27^. The negative association with NAcc volume is consistent with involvement of reward-related circuitry in negative symptoms.

Notably, many associations with SANS Total overlap with those with MAP and EXP (and their domains), suggesting at least some shared anatomical substrate across negative symptom dimensions.

For MAP, although no associations remained significant after FDR correction, several negative and nominal associations mirrored the patterns for SANS Total. Notably, mean SFG, IFGOrb, caudal (c) and rostral (r) MFG showed overlap, while the association with the insula and cMFG were unique to MAP; though exploratory analyses by hemisphere found that both these regions were also correlated with alogia (an EXP subdomain). The insula plays an important role in interoception and integrating sensory, affective, and cognitive processes^52^ which are integral to experiencing pleasure^53,54^. The insula is also involved in reward anticipation and (risky) decision making, where it is thought to integrate internal bodily states with expected reward value to inform goal-directed behavior^55–57^. The cMFG region is a part of the dorsolateral prefrontal cortex implicated in cognitive control and goal-directed behavior^58,59^. Together, roles for the insula and cMFG in MAP are consistent with the idea that individuals with psychosis have difficulty using reward information to guide motivated and goal-directed behavior^60^.

At the level of MAP domains, our findings support several RDoC model hypotheses^14^.

Asociality was nominally associated with mean IFGOrb thickness and NAcc volume. Avolition was significantly associated with hippocampus volume and nominally associated with mean SFG and IFGOrb thickness, and mean amygdala volume. Exploratory analysis by hemisphere found nominal associations of avolition with the left lateral orbitofrontal (lOFC) gyrus. The associations with lOFC thickness and amygdala volume were unique to MAP domains.

Exploratory analyses by hemisphere showed that left lOFC gyrus was associated with anhedonia too. While thinner lOFC has been associated with greater anhedonia scores in depression^61^, it is also involved in reward valuation and in discrimination of rewarding and non-rewarding stimuli^62^. The amygdala is also part of the OFC reward-valuation circuitry^63,64^. Hence, thinner lOFC and lower amygdala and hippocampal volumes suggest impaired differentiation and encoding of rewarding and non-rewarding stimuli, respectively. Such disruptions in reward reinforcement and goal-directed behavior may contribute to lack of motivation to seek rewards.

Although EXP was not significantly associated with any regions after FDR correction, the pattern of associations largely mirrored SANS Total. Mean SFG, pars opercularis (IFGOp), IFGOrb and triangularis (IFGTri), rMFG and precentral gyri in the frontal lobe, and mean supramarginal (SMG) thickness in the parietal region overlapped, while mean IFGOp and IFGTri, precentral, and SMG were unique to EXP and not MAP. Among these, only the association with SMG did not overlap with any of the MAP domains. Of the associations unique to EXP, IFGOp and IFGTri are involved in verbal and non-verbal expressive communication.

These left hemisphere regions comprise Broca’s area and are involved in written and spoken language; their right hemisphere homologs are involved in gesticulation, facial expression, and modulation of timing and intonation of speech^65,66^. A meta-analysis found that medial rostral middle frontal cortex activation was associated with mentalizing mental and emotional states of self and others^67^. While the precentral gyrus is associated with motor initiation and expressive gestures^68^, the SMG has been implicated in phonological processing^69^. Taken together, these regions are consistent with neurocircuitry that underlie expressive deficits.

At the level of EXP domains, our findings support one hypothesis from Strauss’ RDoC model^14^. Specifically, alogia was associated with mean superior temporal gyrus (STG). This association was not only unique to EXP domains but was unique to alogia alone. The STG is essential for processing speech, language, and auditory information^70,71^.

Mean NAcc was associated with EXP and MAP. Additionally, in total there were 10/72 (13.9% FDR and nominally) significant subcortical associations with MAP domains, but only 1/48 (2.1% nominally) with EXP domains, suggesting larger subcortical involvement in MAP compared to EXP domains. The most robust significant associations for cortical thickness were found with alogia, suggesting that cortical thickness in expressive language regions are particularly relevant to alogia. Recent work has addressed relationships between cognitive and MAP symptoms^72^, and the field would benefit from similar work examining relationships between cognitive and expressive negative symptoms.

In our exploratory analysis regarding receptor densities, 5HTT and DAT transporter densities were positively correlated with associations between cortical thickness and nearly all dimensions and domains (except alogia). This finding suggests that 5HTT and DAT transporters are likely relevant to negative symptoms and loss of these transporters may underlie the exacerbation of negative symptoms. 5HTT and DAT findings in schizophrenia are mixed^73–77^ and not well-studied within the context of negative symptoms.

In contrast, A4B2 was negatively correlated with the associations between negative symptom severity and cortical thickness, suggesting possible compensatory or protective mechanisms through upregulation of A4B2 receptors. A4b2 is a nicotinic acetylcholine receptor that has been implicated in cognitive functioning. While this correlation might imply that cognitive disruption is highly intertwined with negative symptom severity, it might also be related to the high rates of tobacco use in individuals with Schizophrenia, which are shown to result in upregulation of A4B2 receptors^78^.

Distinct receptors associated with MAP include CB1 and MOR, and with EXP include NMDA. These finding suggests that MAP and EXP related functions might involve distinct neural substrates with endocannabinoid^79,80^ and opioid receptors^81,82^ (relevant to pleasure and addiction) implicated in MAP domains, and glutamatergic NMDA^83,84^ (implicated in mood disorders) more closely associated with blunted affect (an EXP domain). The distinct receptor correlations, along with more negative correlations with receptor densities for MAP, and more positive correlations with receptor densities for EXP, further support the hierarchical factor structure of negative symptoms^14,15^. This is further corroborated by the stronger correlations between MAP-related domains and MAP versus EXP, and vice versa for both cortical thickness and subcortical volumes.

Our study has several strengths. To the best of our knowledge, this is the first study to examine relationships between brain structure and MAP and EXP dimensions, and their respective domains in schizophrenia. It includes data from 16 samples, addressing non-replication of findings due to small sample size, and reducing site-specific biases, increasing the generalizability. It also suggests that the observed associations are independent from medication treatment. Finally, this study replicated findings based on R-code meta-analysis of results provided by each site with the federated COINSTAC pipeline meta-analysis, supporting the utility of the COINSTAC platform^36^ for consortium meta-analyses.

Several weaknesses must also be noted. While this study examined relationships between brain morphology and negative symptom severity, functional imaging or white matter connectivity studies might be more sensitive in identifying brain-symptom relationships.

Moreover, our analysis focused solely on negative symptoms and did not investigate possible overlap or interactions with positive symptoms or depression. Finally, this study is correlational and does not determine causality. Future research could provide more direct evidence for causal links through assessments of longitudinal changes in brain morphology and functional connectivity and symptoms over time, or combination of imaging with treatment approaches.

Taken together, we found small effect sizes between predominantly frontal lobe and limbic regions and negative symptoms. The shared and unique patterns of associations between brain structure and negative symptom dimensions and domains, and their distinct associations with receptor densities suggest that the use of MAP/EXP dimensions might have utility for exploring brain-symptom relationships. However, as MAP-domains revealed associations with reward-related structures (e.g., amygdala and hippocampus with avolition) and EXP-domains revealed associations with language and expression related structures (e.g., alogia with STG), using the domains may yield more refined mapping of the neural circuitry underlying negative symptoms.

## Acknowledgements

We would like to acknowledge Meelah Hamilton (now deceased), Jesseca E. Rowland, Nicholas Vella, Inika Gillis and Nicole O′Reilly for assistance with data collection and entry. We would also like to thank the volunteers who participated in this study. We acknowledge recruitment assistance from the Australian Schizophrenia Research Bank (ASRB), which was supported by the National Health and Medical Research Council of Australia (NHMRC) Enabling Grant (No. 386500), the Pratt Foundation, Ramsay Health Care, the Viertel Charitable Foundation and the Schizophrenia Research Institute.

Australian Schizophrenia Research Bank, supported by the NHMRC (Enabling Grant, 386500), the Pratt Foundation, Ramsay Health Care, the Viertel Charitable Foundation, and the Schizophrenia Research Institute. Chief Investigators for ASRB were Stanley V. Catts, Patricia T. Michie., Bryan J. Mowry, Ulrich Schall., Rodney J. Scott, Vaughan J. Carr, Frans A. Henskens, Christos Pantelis, Assen Jablensky. We thank Carmel M. Loughland, the ASRB Manager, and acknowledge the help of Jason Bridge for ASRB database queries.

