## Supplement 1 for "Neuroanatomical Correlates of Negative Symptoms in Schizophrenia"

Supplementary Figure 1a. Pattern of Mean Correlations Between Cortical Thickness and SANS Total, MAP and EXP

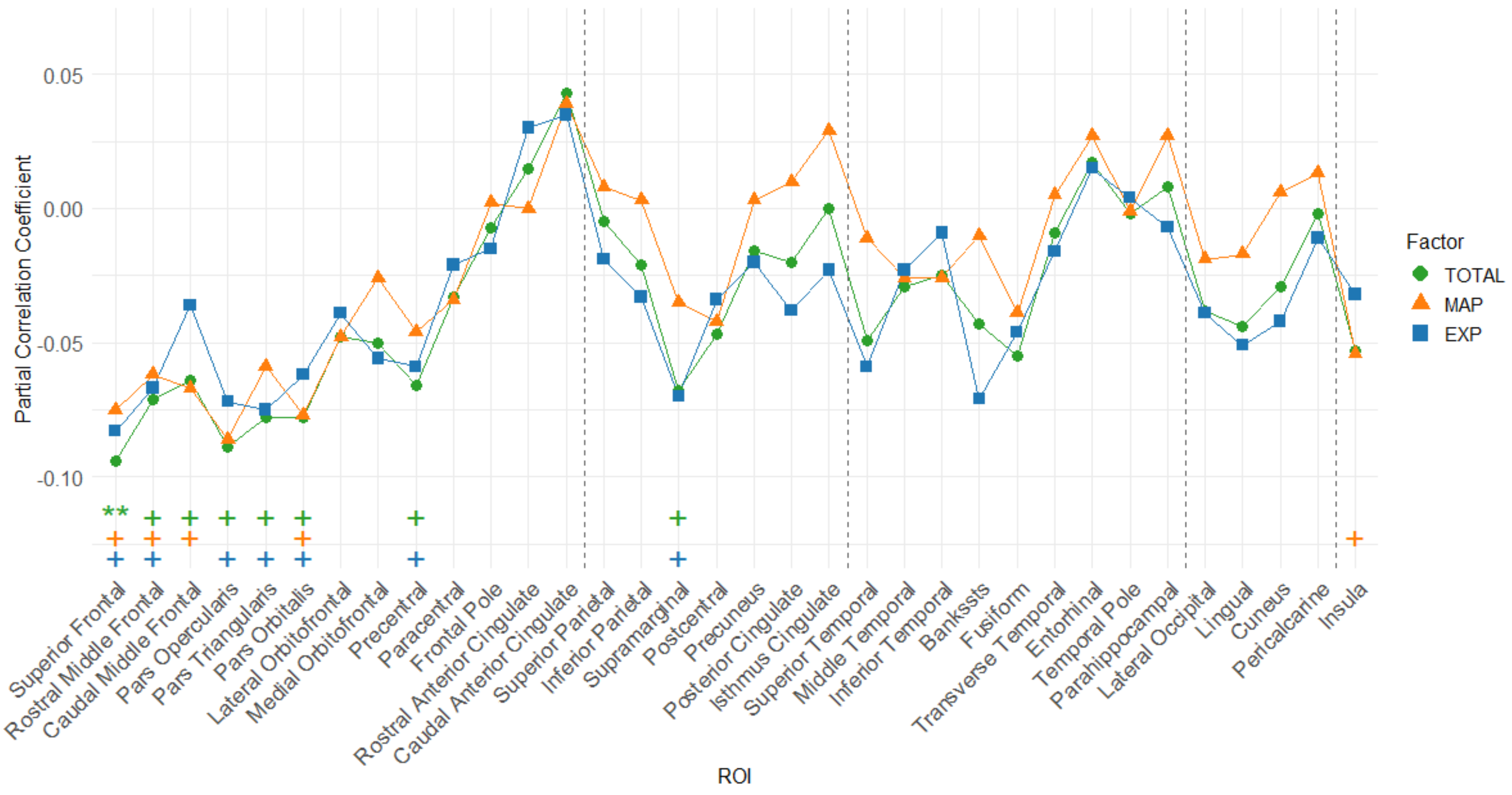


Note: FDR significant denoted with ** and nominal significance denoted with +

Supplementary Figure 1b. Pattern of Mean Correlations Between Cortical Thickness, and MAP dimensions and domains

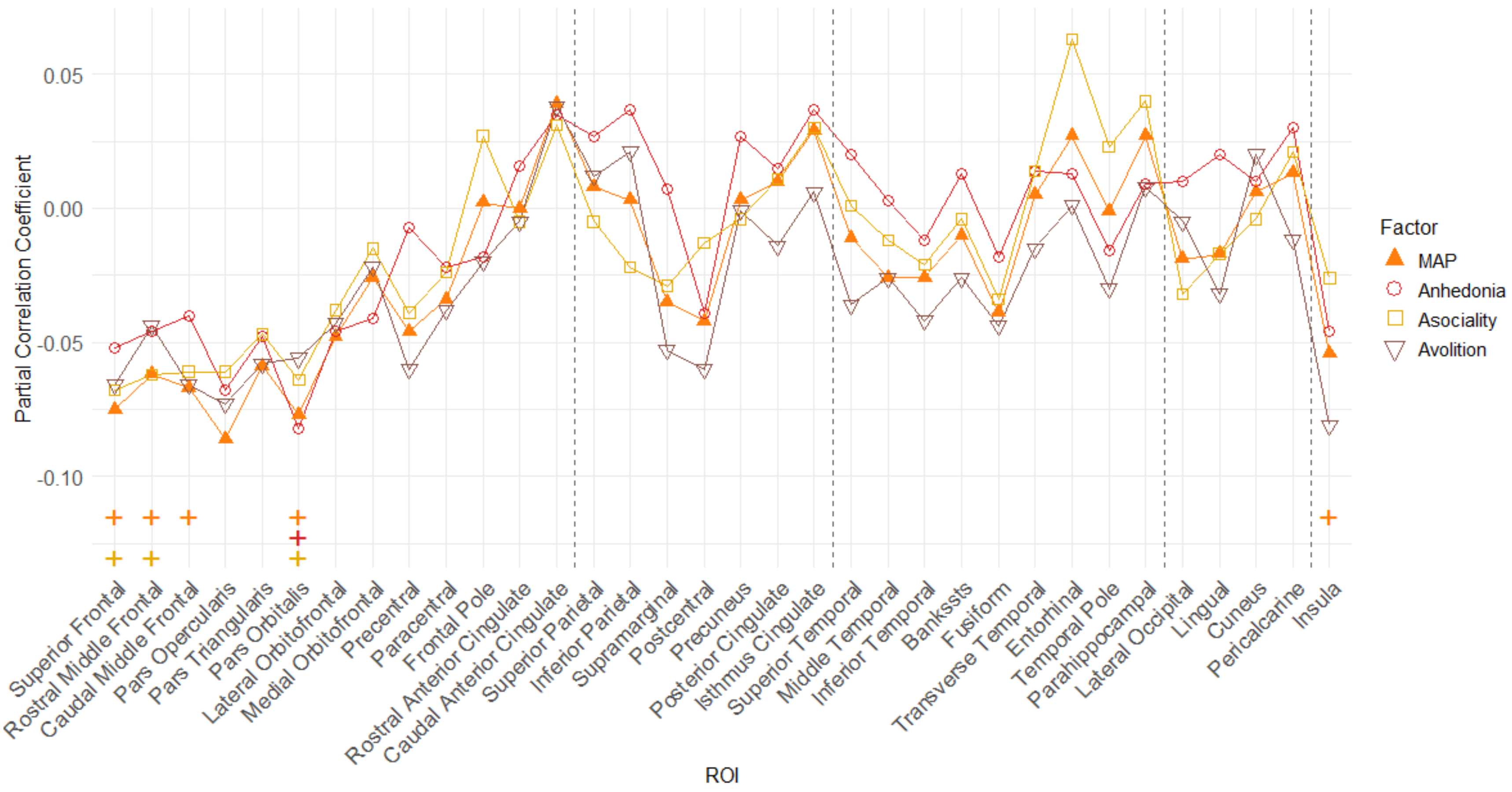


Note: FDR significant denoted with ** and nominal significance denoted with +

Supplementary Figure 1c. Pattern of Mean Correlations Between Cortical Thickness, and EXP dimensions and domains

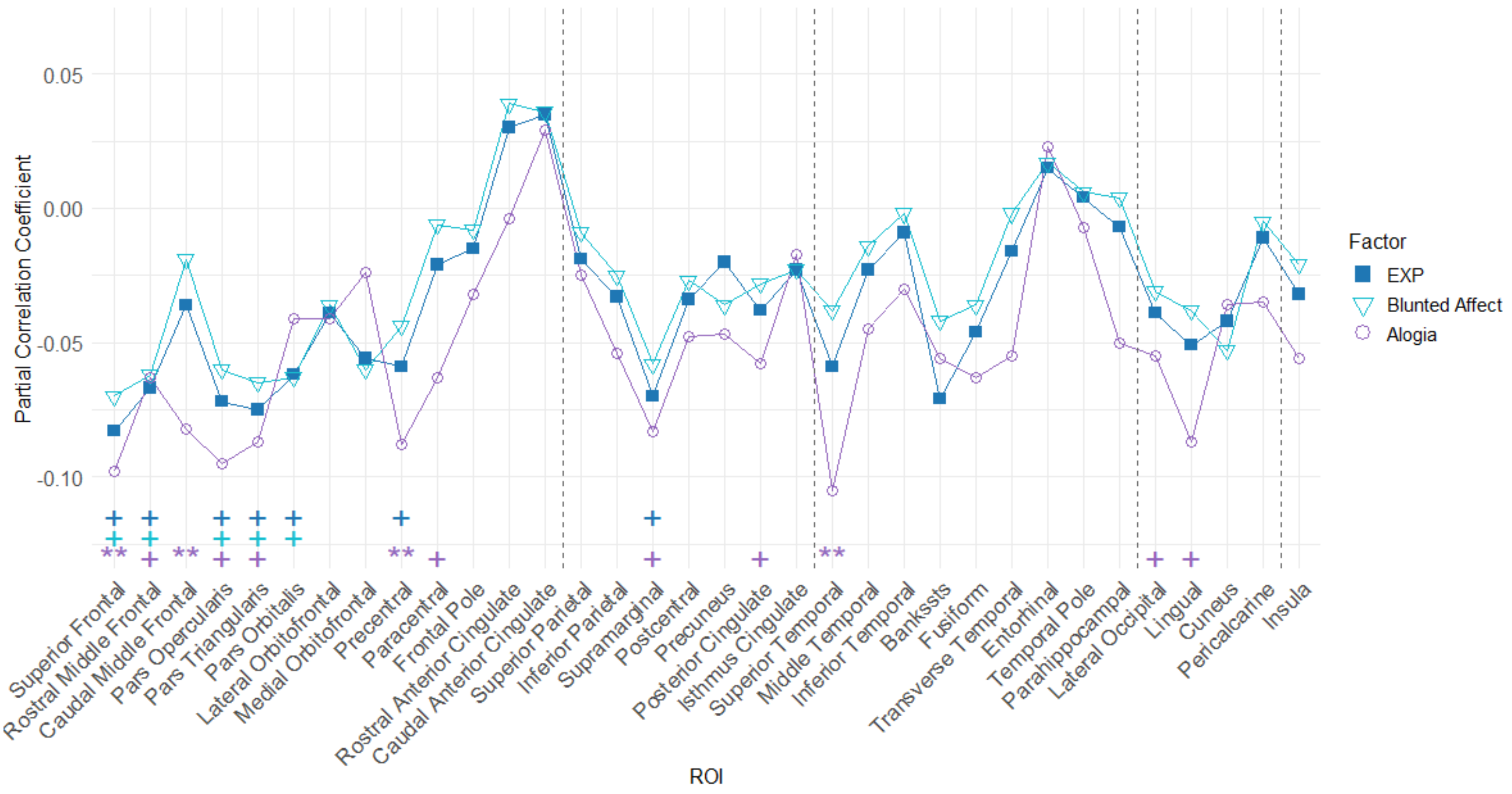


Note: FDR significant denoted with ** and nominal significance denoted with +

Supplementary Figure 2a. Pattern of Mean Correlations Between Subcortical Volumes and SANS Total, MAP and EXP

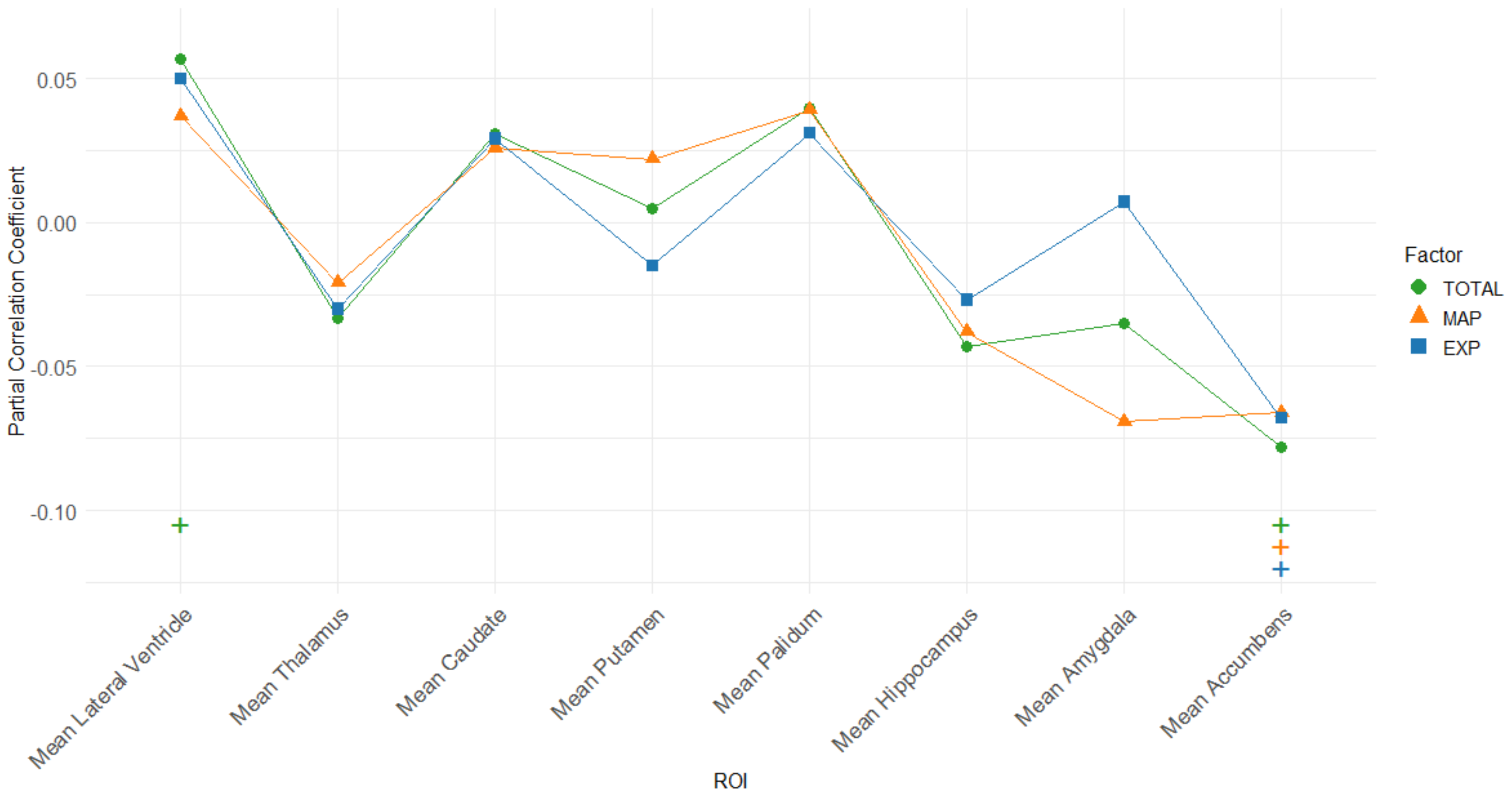


Note: Nominal significance denoted with +

Supplementary Figure 2b. Pattern of Mean Correlations Between Subcortical Volumes, and MAP dimensions and domains

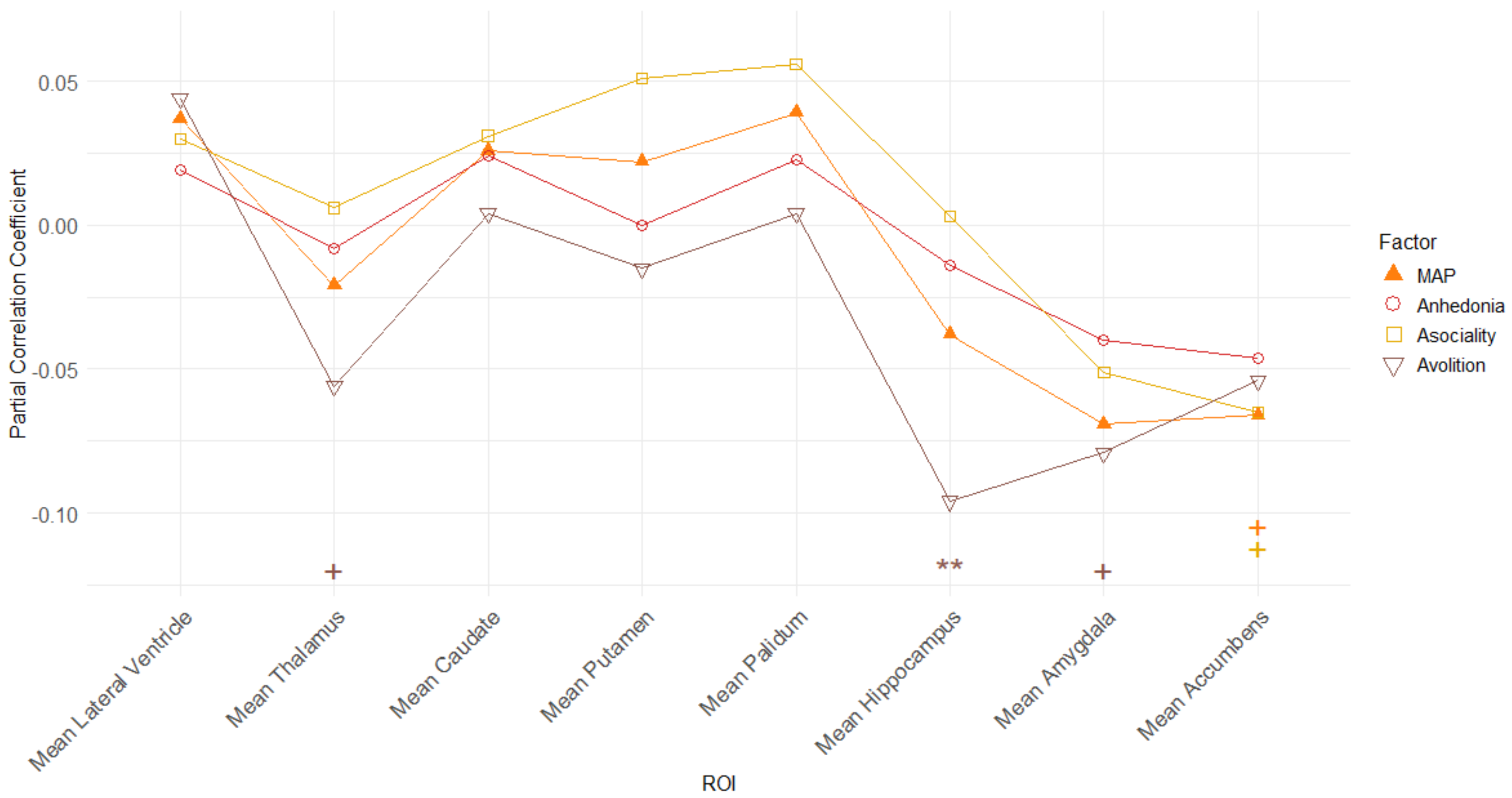


Note: FDR significant denoted with ** and nominal significance denoted with +

Supplementary Figure 2c. Pattern of Mean Correlations Between Subcortical Volumes, and EXP dimensions and domains

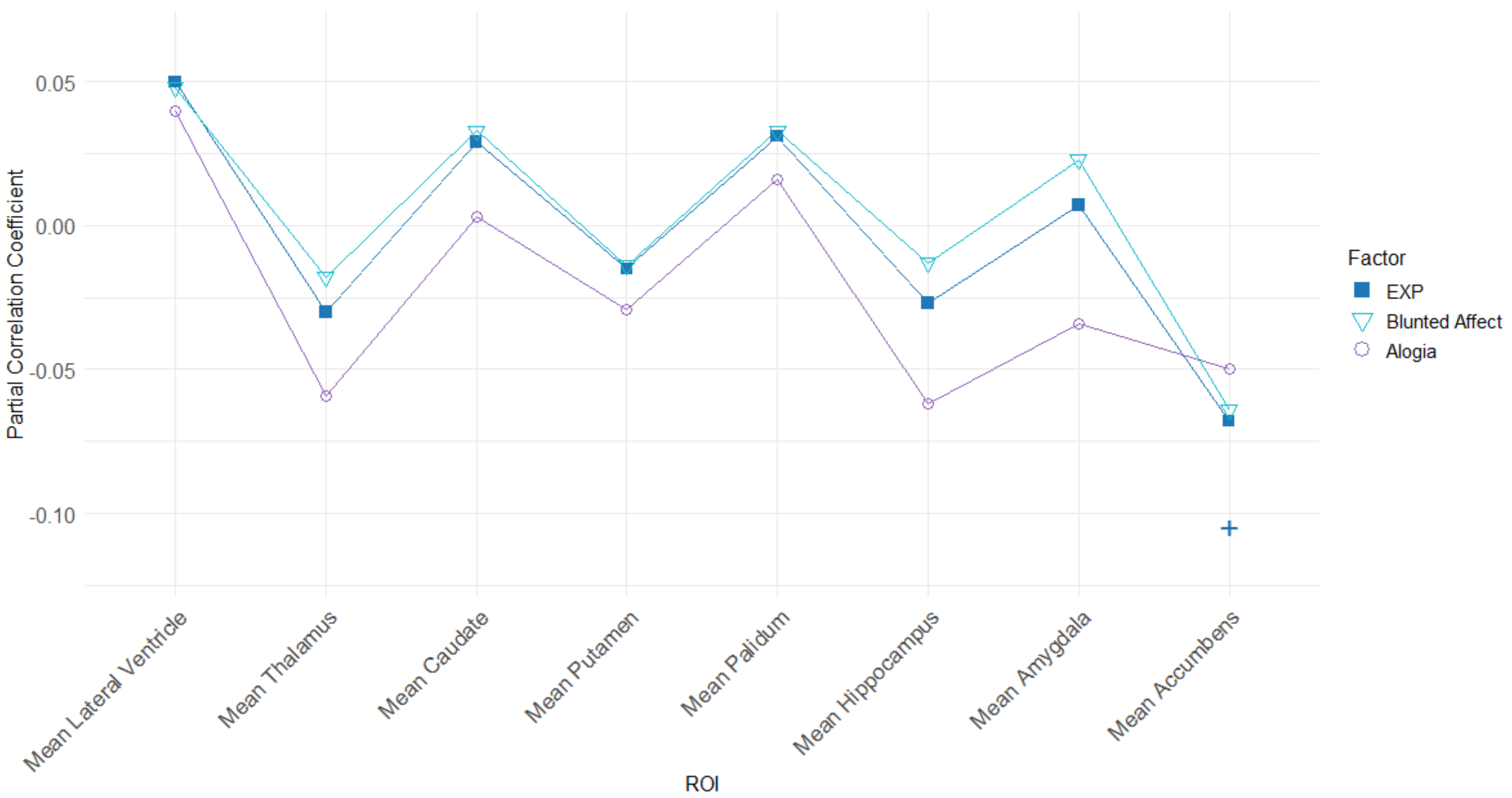


Note: Nominal significance denoted with +

Supplementary Figure 3. Traditional R and COINSTAC meta-analyses yield identical results

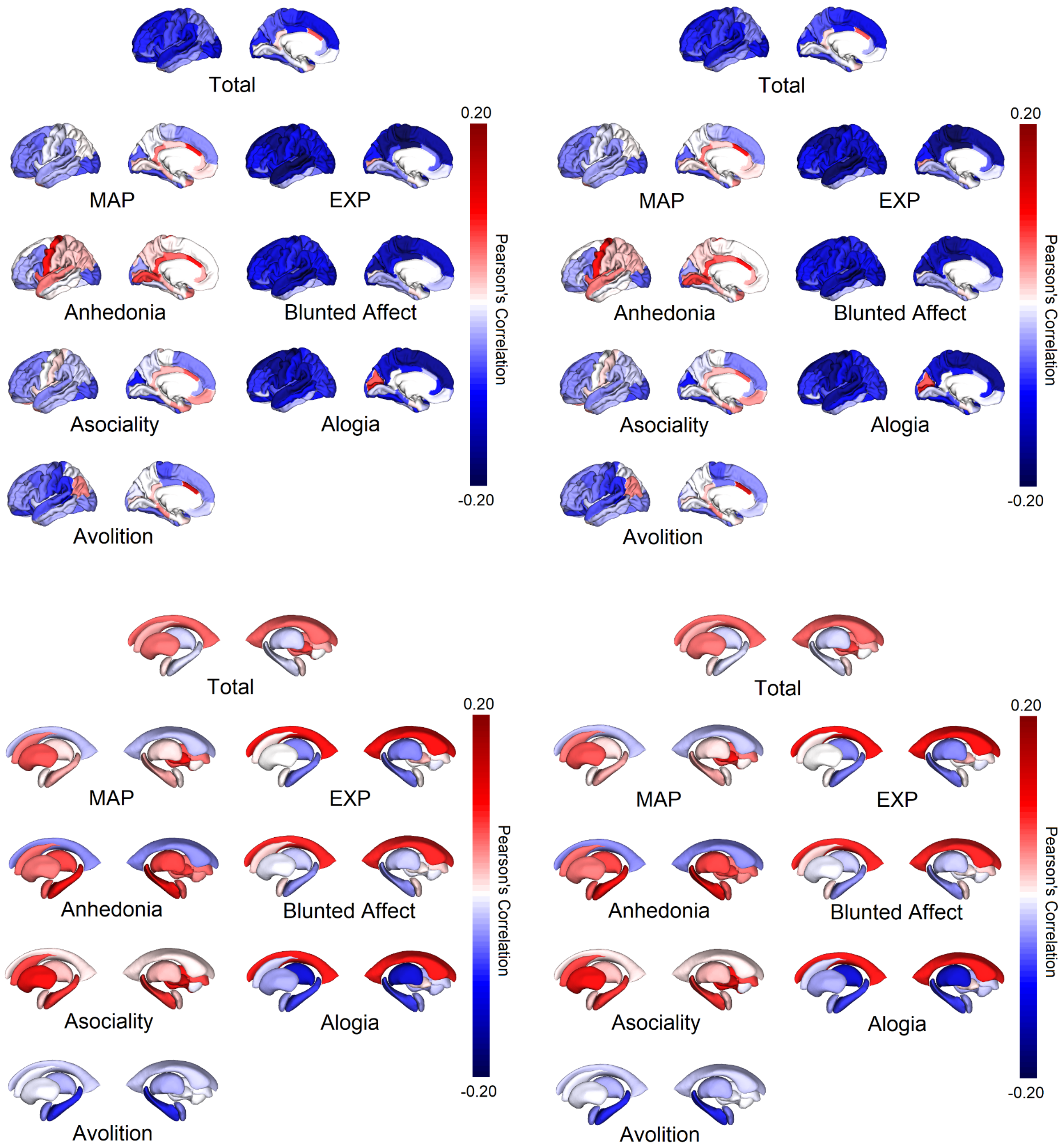


Traditional R meta-analysis (left) and COINSTAC analysis (right); analyses based on data from six sites.

Supplementary Table S1. Sample Image Acquisition and Image Processing Details

| Sample | Number of Scanners | Scanner Vendor & Type | Imaging Protocols | Slice Orientation | FreeSurfer Version | Operating System | Number of subjects removed from analysis due to QC failure |
|---|---|---|---|---|---|---|---|
| ASRB | 5 | Siemens Avanto 1.5T | High-resolution T1-weighted structural magnetic resonance imaging (sMRI) brain scans (MPRAGE) were acquired using an optimized magnetization-prepared rapid acquisition gradient echo on 1.5 T Siemens Avanto scanners (Siemens, Erlangen, Germany) across five Australian research sites (Loughland and al., 2010). Image parameters were set to 176 slices of 1mm thickness, no gap with field-of-view 250 x 250 mm2, repetition time 1980 ms, echo time 4.3 ms, data acquisition matrix 256 x 256, with a flip matrix of 15°, resulting in a voxel size of 0.98×0.98×1.0 mm3 | Sagittal | v5.1.0 | Mac OSX | 0 |

| | | | | | | | |
|---|---|---|---|---|---|---|---|
| FBIRN (Phase3) | 7 | 3T Siemens Tim Trio; 3T GE | High-resolution structural imaging scans were acquired on six 3T Siemens Tim® Trio System and one 3T General Electric Discovery MR750 scanner. MP-RAGE scan parameters for the Siemens scanner were: scan plane=sagittal, TR/TE/TI=2300/2.94/1100 ms, GRAPPA acceleration factor=2, flip angle=9°, resolution=256×256x160, FOV=220mm2, voxel size=0.86x0.86x1.2mm, and NEX=1. IR-SPGR scan parameters for the General Electric scanner were: scan plane=sagittal, TR/TE/TI=5.95/1.99/450ms , ASSET acceleration factor=2, a flip angle=12°, resolution=256×256x166, FOV=220mm2, voxel size=0.86x0.86x1.2mm, and NEX=1. All scans covered the entire brain. | Sagittal | v5.3.0 | Centos 64bit | 0 |
| FIDMAG | 1 | 1.5T GE Signa | 180 axial slices; 1mm slice thickness, no gap, matrix size 512x512; 0.5x0.5x1mm3 voxel resolution; TE 4ms, TR 2000ms, flip angle 15º | Axial | v5.3.0 | Linux Ubuntu | |
| FSLRome | 1 | 3T Siemens Allegra | T1-weighted, 3D MDEFT, 1x1x1xmm, TE/TR =2.4/7.92 ms, flip angle=15 | Sagittal | 6.0dev | linux | |
| FOR2107-MR | 1 | 3T Siemens Magnetom TiroTim syngo | Site Marburg: 3D T1-weighted magnetization prepared rapid acquisition gradient echo (MPRAGE); TR=1900ms, TE=2.26ms, TI=900ms, FA=9°, voxel size=1.0x1.0x1.0mm³, Acquisition Direction Sagittal, 176 slices, slice gap 0.5mm. | Sagittal | v5.3.0 | Red Hat Enterprise Linux Server release 5.11 (Tikanga) | 0 |

| | | | | | | | |
|---|---|---|---|---|---|---|---|
| FOR2017-MS | 1 | 3T Siemens PRISMA | T1-weighted magnetization prepared rapid acquisition gradient echo (MPRAGE); TR=2130ms, TE=2.28ms, TI=900ms, FA=8°, voxel size=1.0x1.0x1.0mm³, Acquisition Direction Sagittal, 192 slices, slice gap 0.5mm. | Sagittal | v5.3.0 | Red Hat Enterprise Linux Server release 5.11 (Tikanga) | 0 |
| GIPSI | 1 | 3T Philips Ingenia | 3D T1-weighted TFE sequence: TR=4.7ms, TE=2ms, flip angle=8°, voxel size=1.0x0.6x0.6mm³ (no gap) and 160 axial slices | Axial | v5.3.0 | Mac OSX | 0 |
| HUBIN | 1 | 1.5T Signa HDxt | T1-weighted SPGR pulse sequence: TR=24ms, TE=6ms, FOV=240mm, acquisition matrix=256x192, 124 slices, flip angle=35 degrees, slice thickness=1.5mm | Coronal | v5.3.0 | Linux | NA |
| IGP | 1 | 3T Philips Achieva TX | 3D MPRAGE; TR=8.9ms, TE=4.1ms, FOV=240mm, matrix 268 x 268, 200 slices, slice thickness 0.9mm (no gap) | Sagittal | v5.3.0 | Mac OSX | 0 |
| MCIC | 3 | 1.5, 3T Siemens and GE | T1 scans: TR = 2530 ms for 3 T, TR = 12 ms for 1.5 T; TE = 3.79 ms for 3 T, TE = 4.76 ms for 1.5 T; FA = 7 for 3 T, FA = 20 for 1.5 T; TI = 1100 for 3 T; Bandwidth = 181 for 3 T, Bandwidth = 110 for 1.5 T; 0.625×0.625 mm voxel size; slice thickness 1.5 mm; FOV 256×256×128 cm matrix; FOV = 16 cm (could be increased to 18 cm when needed for full brain coverage). | Coronal | v4.0.1 | Linux of various flavors | 5 subjects failed automated segmentation procedure due to excessive motion artifacts; 2 participants' MRI data failed the manual inspection |

| | | | | | | | |
|---|---|---|---|---|---|---|---|
| NU | 1 | 1.5T Vision | 1) 3D turbo-FLASH: TR=20 ms, TE=5.4 ms, flip=30˚, ACQ=1, 256x256 matrix, 1xl mm in-plane resolution, 180 slices, slice thickness 1 mm, 13:30 min scan time and 2) 3D MPRAGE (2-4 repeats): TR=9.7 ms, TE=4 ms, flip=10˚, ACQ=1, 256x256 matrix, 1xl mm in-plane resolution, 128 slices, slice thickness 1.25 mm, 5:36 min scan time each | Axial | v5.3.0 | centos6 x86_64 | |
| SCORE | 1 | | MPRAGE: aquisition matrix: 256×256×176, isotropic spatial resolution: 1x1x1mm3, TI=1000ms, TR=2s, TE=3.4 ms, flip angle: 8° and bandwidth of 200 Hz/pixel | Sagittal | 6.0dev | | |
| UCISZ | 1 | Philips 3T Achieva | T1TFE:200 sagittal slices, 320x274 matrix size, .75mm isotropic, TR = 11ms, TE =4.562ms, flip angle = 18° | Sagittal | v6.0dev | Centos 3.10.72-1.el6.elrepo.x86_64 | 0 |
| UMCG | 1 | Philips 3T Intera | WIP T1 3D SENSE, 170 slices, 1mm slice thickness, inplane matrix 256x256, FOV: 230x170x256mm, flip angle 9 degrees) | Axial | v6.0.0 | Ubuntu 16.04 LTS | 0 |
| UPENN | 2 | Siemens 3T | MPRAGE, TR=1810 ms, TE= 3.51 ms, TI=1100 ms, flip angle 9, FOV= 240 x 180 mm, matrix= 256 × 192, resolution = 0.9 x 0.9 mm, slices = 160, slice/skip thickness = 1 mm/0 mm | Axial | v5.3.0 | Linux Red Hat Enterprise 5 | 0 |
| ZURICH | 1 | Philips 3T | 3D T1-weighted images were acquired with an ultrafast gradient echo T1-weighted sequence (TR=8.4ms, TE=3.8ms, flip angle=8°) in 160 sagittal plan slices (1mm slice thickness, no slice gap) of 240×240mm2 resulting in 1x1x1mm3voxels. | Sagittal | v6.0.0 | Linux | 0 |

Supplementary Table S2. Sample Breakdown

| Sample | N | Gender M/F | Mean Age (Years) | Mean Age of Onset (Years) | Mean Duration of Illness | PANSS Total | PANSS Neg | PANSS Pos | SANS Total | SAPS Total | Mean CPZ | N No Antipsychotic Medication | N First Generation AP | N Second Generation AP | N Both First- and Second Generation AP |
|---|---|---|---|---|---|---|---|---|---|---|---|---|---|---|---|
| ASRB | 263 | 177/86 | 38.6 | 23.6 | 15.0 | ... | ... | ... | 18.5 | ... | ... | 44 | 198 | 12 | 9 |
| FBIRN | 187 | 141/46 | 38.9 | 21.8 | 17.2 | 58.7 | 14.6 | 15.5 | 19.6 | 19.0 | 372.6 | 0 | 139 | 20 | 10 |
| FIDMAG | 48 | 35/13 | 39.7 | 23.8 | 15.8 | 74.4 | 23.5 | 15.4 | 37.2 | ... | 570.8 | 0 | 36 | 4 | 8 |
| FOR2107-MR | 37 | 23/14 | 37.2 | 21.2 | 15.9 | ... | ... | ... | 18.8 | 13.2 | 403.7 | 6 | 25 | 3 | 3 |
| FOR2107-MS | 8 | 4/4 | 33.4 | 22.2 | 11.1 | ... | ... | ... | 8.1 | 6.4 | 306.7 | 1 | 7 | 0 | 0 |
| FSLRome | 164 | 110/54 | 39.4 | 24.5 | 14.9 | 86.3 | 21.0 | 20.9 | 28.8 | 31.5 | 302.2 | 11 | 82 | 26 | 42 |
| GIPSI | 40 | 32/8 | 33.0 | 19.1 | 13.9 | ... | ... | ... | 33.3 | 9.6 | 434.0 | 3 | 27 | 2 | 8 |
| HUBIN | 94 | 70/24 | 41.7 | 24.5 | 17.1 | ... | ... | ... | 22.3 | 9.0 | 272.7 | 6 | 38 | 40 | 10 |
| IGP | 68 | 40/28 | 41.7 | 22.9 | 18.8 | 55.5 | 14.5 | 13.8 | 6.9 | 13.0 | 655.2 | 10 | 53 | 3 | 2 |
| MCIC | 148 | 113/35 | 32.9 | 22.8 | 10.2 | ... | ... | ... | 23.3 | 22.8 | 533.5 | 8 | 117 | 10 | 7 |
| NU | 108 | 74/34 | 34.2 | 20.9 | 13.2 | ... | ... | ... | 33.0 | 23.2 | ... | 9 | 79 | 17 | 0 |
| SCORE | 141 | 98/43 | 25.7 | 24.5 | 1.1 | ... | ... | ... | 15.1 | ... | 207.9 | 104 | 35 | 2 | 0 |
| UCISZ | 27 | 22/5 | 42.9 | 25.0 | 17.5 | 60.0 | 16.0 | 15.6 | 22.8 | 13.4 | ... | ... | ... | ... | ... |
| UMCG | 21 | 16/5 | 37.9 | 26.7 | 11.2 | 65.2 | 19.3 | 13.0 | 40.1 | ... | ... | 2 | 19 | 0 | 0 |
| UPENN | 177 | 105/72 | 38.9 | 20.7 | 17.3 | ... | ... | ... | 23.7 | 18.3 | 481.6 | 0 | 65 | 13 | 5 |
| ZURICH | 60 | 45/15 | 30.5 | 22.2 | 8.4 | 48.6 | 14.5 | 10.7 | 24.9 | ... | 494.8 | 3 | 55 | 0 | 2 |
| Total | 1591 | 1105/ 486 | 36.7 | 22.8 | 13.7 | 66.7 | 17.4 | 16.2 | 22.6 | 19.7 | 405.1 | 207 | 975 | 152 | 106 |

Supplementary Table S3. Higher Negative Symptom Severity in Men than Women with Schizophrenia

| | **Cohen's *d*** | **SE** | **95% CI** | ***t*-score** | ***p*-value** | ***N* males** | ***N* females** | **Total *N*** |
|---|---|---|---|---|---|---|---|---|
| SANS Total | -0.233 | 0.064 | [-0.371 - -0.096] | -3.647 | 0.003 | 1207 | 774 | 1981 |
| EXP | -0.213 | 0.063 | [-0.348 - -0.077] | -3.366 | 0.005 | 1269 | 801 | 2070 |
| MAP | -0.182 | 0.059 | [-0.309 - -0.055] | -3.082 | 0.008 | 1213 | 777 | 1990 |
| Anhedonia | -0.131 | 0.045 | [-0.229 - -0.034] | -2.899 | 0.012 | 1275 | 803 | 2078 |
| Avolition | -0.139 | 0.052 | [-0.252 - -0.027] | -2.655 | 0.019 | 1216 | 780 | 1996 |
| Asociality | -0.217 | 0.063 | [-0.352 - -0.082] | -3.445 | 0.004 | 1272 | 804 | 2076 |
| Blunted Affect | -0.212 | 0.062 | [-0.345 - -0.080] | -3.436 | 0.004 | 1273 | 805 | 2078 |
| Alogia | -0.144 | 0.053 | [-0.257 - -0.031] | -2.740 | 0.016 | 1274 | 801 | 2075 |

Note: These comparisons are based on cortical data of all individuals with schizophrenia from each site with SANS data, including ones without imaging data.

Supplementary Table S4. Sensitivity Analysis Comparing Effect Sizes of Main Analysis Against Models with the Inclusion of Medication Type and Medication Dose

| | Only sex and age | | | | WAP (medication type) | | | | WCPZ (medication dose) | | | | |
|---|---|---|---|---|---|---|---|---|---|---|---|---|---|
| ROI | Partial *r* | *N* | *p* | FDR | Partial *r* | *N* | *p* | FDR | Partial *r* | *N* | *p* | FDR | Factor |
| Left Pars Orbitalis | -0.078 | 1327 | 0.002 | 0.035 | -0.074 | 1193 | 0.005 | 0.114 | -0.044 | 747 | 0.306 | 0.960 | CT SANS Total |
| Left Superior Frontal | -0.084 | 1350 | 0.001 | 0.031 | -0.071 | 1215 | 0.004 | 0.114 | -0.090 | 746 | 0.010 | 0.326 | CT SANS Total |
| Right Superior Frontal | -0.096 | 1341 | <0.001 | 0.03 | -0.078 | 1206 | 0.002 | 0.114 | -0.094 | 750 | 0.003 | 0.222 | CT SANS Total |
| Mean Superior Frontal | -0.094 | 1325 | <0.001 | 0.012 | -0.077 | 1190 | 0.002 | 0.059 | -0.097 | 744 | 0.004 | 0.144 | CT SANS Total |
| Left Pars Triangularis | -0.086 | 1304 | 0.001 | 0.027 | -0.081 | 1171 | 0.002 | 0.088 | -0.076 | 746 | 0.024 | 0.329 | CT Avolition |
| Right Caudal Middle Frontal | -0.078 | 1337 | 0.001 | 0.027 | -0.077 | 1202 | 0.003 | 0.088 | -0.076 | 749 | 0.019 | 0.329 | CT Avolition |
| Left Rostral Middle Frontal | -0.08 | 1312 | 0.002 | 0.044 | -0.088 | 1180 | 0.006 | 0.083 | -0.113 | 745 | 0.001 | 0.023 | CT Alogia |
| Left Superior Frontal | -0.101 | 1350 | <0.001 | 0.014 | -0.093 | 1215 | 0.001 | 0.082 | -0.13 | 746 | 0.001 | 0.023 | CT Alogia |
| Right Caudal Middle Frontal | -0.099 | 1337 | 0.003 | 0.044 | -0.102 | 1202 | 0.004 | 0.082 | -0.107 | 749 | 0.018 | 0.172 | CT Alogia |
| Right Superior Frontal | -0.09 | 1341 | 0.001 | 0.019 | -0.082 | 1206 | 0.004 | 0.082 | -0.108 | 750 | 0.001 | 0.023 | CT Alogia |
| Mean Caudal Middle Frontal | -0.082 | 1323 | 0.005 | 0.043 | -0.092 | 1189 | 0.004 | 0.04 | -0.094 | 748 | 0.020 | 0.112 | CT Alogia |
| Mean Lingual | -0.087 | 1328 | 0.008 | 0.057 | -0.106 | 1193 | 0.005 | 0.04 | -0.094 | 729 | 0.058 | 0.198 | CT Alogia |
| Mean Precentral | -0.088 | 1318 | 0.005 | 0.043 | -0.09 | 1185 | 0.006 | 0.043 | -0.102 | 743 | 0.006 | 0.072 | CT Alogia |
| Mean Rostral Middle Frontal | -0.063 | 1263 | 0.013 | 0.065 | -0.074 | 1131 | 0.014 | 0.081 | -0.089 | 737 | 0.002 | 0.028 | CT Alogia |
| Mean Superior Frontal | -0.098 | 1325 | <0.001 | 0.007 | -0.088 | 1190 | 0.002 | 0.04 | -0.124 | 744 | <0.001 | 0.01 | CT Alogia |
| Mean Superior Temporal | -0.105 | 1132 | 0.002 | 0.034 | -0.103 | 1001 | 0.003 | 0.04 | -0.102 | 680 | 0.010 | 0.085 | CT Alogia |
| Left Lateral Ventricle | 0.050 | 1263 | 0.018 | 0.143 | 0.05 | 1211 | 0.001 | 0.017 | 0.064 | 787 | 0.001 | 0.012 | SV SANS Total |
| Mean Lateral Ventricle | 0.057 | 1263 | 0.013 | 0.093 | 0.055 | 1211 | 0.002 | 0.014 | 0.074 | 787 | 0.002 | 0.016 | SV SANS Total |
| Left Hippocampus | -0.092 | 1339 | 0.001 | 0.007 | -0.087 | 1196 | 0.005 | 0.038 | -0.065 | 776 | 0.093 | 0.497 | SV Avolition |
| Right Hippocampus | -0.096 | 1349 | 0.001 | 0.007 | -0.093 | 1206 | 0.002 | 0.038 | -0.084 | 785 | 0.012 | 0.199 | SV Avolition |
| Mean Hippocampus | -0.096 | 1335 | <0.001 | 0.004 | -0.092 | 1193 | 0.003 | 0.021 | -0.075 | 774 | 0.033 | 0.264 | SV Avolition |

Note: CT = Cortical Thickness and SV = Subcortical Volumes

Supplementary Table S5 Associations between Cortical Thickness and SANS Total

| ROI | Partial *r* | SE | *t*-Score | 95% CI | I2 | *N* | *p-value* | FDR |
|---|---|---|---|---|---|---|---|---|
| Left Bankssts | -0.041 | 0.014 | -1.462 | [-0.100 - 0.019] | 0.000 | 1266 | 0.166 | 0.476 |
| *Left Caudal Anterior Cingulate* | *0.054* | *0.011* | *2.382* | *[ 0.005 - 0.103]* | *0.000* | *1362* | *0.032* | *0.235* |
| Left Caudal Middle Frontal | -0.035 | 0.014 | -1.260 | [-0.096 - 0.025] | 0.000 | 1351 | 0.228 | 0.476 |
| Left Cuneus | -0.010 | 0.018 | -0.278 | [-0.087 - 0.067] | 0.000 | 1275 | 0.785 | 0.905 |
| Left Entorhinal | 0.000 | 0.011 | -0.011 | [-0.046 - 0.046] | 0.000 | 1278 | 0.992 | 0.992 |
| *Left Fusiform* | *-0.067* | *0.015* | *-2.175* | *[-0.133 - -0.001]* | *0.000* | *1308* | *0.047* | *0.268* |
| Left Inferior Parietal | 0.004 | 0.015 | 0.126 | [-0.059 - 0.066] | 0.000 | 1234 | 0.901 | 0.973 |
| Left Inferior Temporal | -0.022 | 0.014 | -0.778 | [-0.083 - 0.039] | 0.000 | 1275 | 0.449 | 0.650 |
| Left Isthmus Cingulate | -0.009 | 0.014 | -0.329 | [-0.067 - 0.049] | 0.000 | 1361 | 0.747 | 0.891 |
| Left Lateral Occipital | -0.038 | 0.016 | -1.214 | [-0.105 - 0.029] | 0.000 | 1296 | 0.245 | 0.476 |
| Left Lateral Orbitofrontal | -0.040 | 0.013 | -1.520 | [-0.096 - 0.016] | 0.000 | 1364 | 0.151 | 0.469 |
| Left Lingual | -0.039 | 0.015 | -1.309 | [-0.103 - 0.025] | 0.000 | 1339 | 0.212 | 0.476 |
| Left Medial Orbitofrontal | -0.039 | 0.016 | -1.221 | [-0.108 - 0.030] | 0.000 | 1347 | 0.242 | 0.476 |
| Left Middle Temporal | -0.029 | 0.014 | -1.077 | [-0.087 - 0.029] | 0.000 | 1218 | 0.300 | 0.550 |
| Left Parahippocampal | -0.009 | 0.015 | -0.295 | [-0.075 - 0.057] | 0.000 | 1342 | 0.772 | 0.905 |
| Left Paracentral | -0.035 | 0.012 | -1.420 | [-0.088 - 0.018] | 0.000 | 1360 | 0.177 | 0.476 |
| *Left Pars Opercularis* | *-0.094* | *0.016* | *-2.940* | *[-0.162 - -0.026]* | *0.000* | *1323* | *0.011* | *0.110* |
| ***Left Pars Orbitalis*** | ***-0.078*** | ***0.010*** | ***-3.915*** | ***[-0.120 - -0.035]*** | ***0.000*** | ***1327*** | ***0.002*** | ***0.035*** |
| *Left Pars Triangularis* | *-0.069* | *0.012* | *-2.889* | *[-0.121 - -0.018]* | *0.000* | *1304* | *0.012* | *0.110* |
| Left Pericalcarine | 0.013 | 0.017 | 0.396 | [-0.058 - 0.084] | 0.000 | 1350 | 0.698 | 0.879 |
| Left Postcentral | -0.048 | 0.015 | -1.620 | [-0.110 - 0.015] | 0.000 | 1336 | 0.127 | 0.433 |
| Left Posterior Cingulate | -0.028 | 0.015 | -0.957 | [-0.090 - 0.035] | 0.000 | 1363 | 0.355 | 0.587 |
| *Left Precentral* | *-0.063* | *0.014* | *-2.183* | *[-0.125 - -0.001]* | *0.000* | *1344* | *0.047* | *0.268* |
| Left Precuneus | -0.034 | 0.012 | -1.415 | [-0.087 - 0.018] | 0.000 | 1337 | 0.179 | 0.476 |
| Left Rostral Anterior Cingulate | 0.029 | 0.011 | 1.262 | [-0.020 - 0.078] | 0.000 | 1347 | 0.228 | 0.476 |
| *Left Rostral Middle Frontal* | *-0.061* | *0.011* | *-2.847* | *[-0.107 - -0.015]* | *0.000* | *1312* | *0.013* | *0.110* |
| ***Left Superior Frontal*** | ***-0.084*** | ***0.010*** | ***-4.180*** | ***[-0.127 - -0.041]*** | ***0.000*** | ***1350*** | ***0.001*** | ***0.031*** |
| Left Superior Parietal | -0.002 | 0.019 | -0.044 | [-0.081 - 0.078] | 23.374 | 1283 | 0.966 | 0.986 |
| Left Superior Temporal | -0.055 | 0.014 | -1.955 | [-0.114 - 0.005] | 0.000 | 1197 | 0.071 | 0.283 |
| *Left Supramarginal* | *-0.056* | *0.012* | *-2.340* | *[-0.106 - -0.005]* | *0.000* | *1216* | *0.035* | *0.235* |
| Left Frontal Pole | -0.018 | 0.011 | -0.800 | [-0.065 - 0.030] | 0.000 | 1368 | 0.437 | 0.646 |
| Left Temporal Pole | -0.009 | 0.011 | -0.437 | [-0.054 - 0.036] | 0.000 | 1329 | 0.669 | 0.858 |
| Left Transverse Temporal | 0.007 | 0.014 | 0.255 | [-0.052 - 0.065] | 0.000 | 1362 | 0.802 | 0.909 |
| Left Insula | -0.040 | 0.015 | -1.355 | [-0.103 - 0.023] | 0.000 | 1365 | 0.197 | 0.476 |
| Right Bankssts | -0.040 | 0.023 | -0.857 | [-0.139 - 0.060] | 57.461 | 1317 | 0.406 | 0.613 |
| Right Caudal Anterior Cingulate | 0.003 | 0.016 | 0.088 | [-0.067 - 0.073] | 0.000 | 1361 | 0.931 | 0.986 |
| *Right Caudal Middle Frontal* | *-0.085* | *0.013* | *-3.331* | *[-0.139 - -0.030]* | *0.000* | *1337* | *0.005* | *0.084* |
| Right Cuneus | -0.021 | 0.015 | -0.724 | [-0.084 - 0.042] | 0.000 | 1316 | 0.481 | 0.674 |
| Right Entorhinal | 0.032 | 0.012 | 1.277 | [-0.021 - 0.084] | 0.000 | 1164 | 0.222 | 0.476 |
| Right Fusiform | -0.034 | 0.014 | -1.188 | [-0.095 - 0.027] | 0.000 | 1281 | 0.255 | 0.481 |
| Right Inferior Parietal | -0.028 | 0.019 | -0.716 | [-0.110 - 0.055] | 0.000 | 1226 | 0.486 | 0.674 |
| Right Inferior Temporal | -0.033 | 0.018 | -0.923 | [-0.111 - 0.044] | 13.384 | 1283 | 0.371 | 0.587 |
| Right Isthmus Cingulate | 0.008 | 0.012 | 0.341 | [-0.044 - 0.060] | 0.000 | 1351 | 0.739 | 0.891 |
| Right Lateral Occipital | -0.033 | 0.017 | -0.953 | [-0.106 - 0.041] | 0.000 | 1287 | 0.357 | 0.587 |
| Right Lateral Orbitofrontal | -0.057 | 0.016 | -1.816 | [-0.124 - 0.010] | 0.000 | 1333 | 0.091 | 0.343 |
| Right Lingual | -0.029 | 0.014 | -1.060 | [-0.089 - 0.030] | 0.000 | 1350 | 0.307 | 0.550 |
| Right Medial Orbitofrontal | -0.047 | 0.018 | -1.347 | [-0.122 - 0.028] | 7.462 | 1316 | 0.199 | 0.476 |
| Right Middle Temporal | -0.028 | 0.020 | -0.698 | [-0.112 - 0.057] | 23.117 | 1255 | 0.497 | 0.676 |

| ROI | Partial *r* | SE | *t*-Score | 95% CI | *I2* | *N* | *p-value* | FDR |
|---|---|---|---|---|---|---|---|---|
| Right Parahippocampal | 0.022 | 0.012 | 0.935 | [-0.028 - 0.072] | 0.000 | 1322 | 0.365 | 0.587 |
| Right Paracentral | -0.029 | 0.010 | -1.516 | [-0.070 - 0.012] | 0.000 | 1361 | 0.152 | 0.469 |
| Right Pars Opercularis | -0.063 | 0.016 | -1.980 | [-0.131 - 0.005] | 0.000 | 1305 | 0.068 | 0.283 |
| Right Pars Orbitalis | -0.035 | 0.013 | -1.366 | [-0.090 - 0.020] | 0.000 | 1336 | 0.193 | 0.476 |
| Right Pars Triangularis | -0.055 | 0.013 | -2.066 | [-0.113 - 0.002] | 0.000 | 1305 | 0.058 | 0.283 |
| Right Pericalcarine | -0.016 | 0.014 | -0.579 | [-0.077 - 0.045] | 0.000 | 1350 | 0.572 | 0.762 |
| Right Postcentral | -0.035 | 0.014 | -1.241 | [-0.096 - 0.026] | 0.000 | 1342 | 0.235 | 0.476 |
| Right Posterior Cingulate | -0.012 | 0.013 | -0.481 | [-0.067 - 0.043] | 0.000 | 1364 | 0.638 | 0.834 |
| Right Precentral | -0.040 | 0.010 | -2.025 | [-0.083 - 0.002] | 0.000 | 1333 | 0.062 | 0.283 |
| Right Precuneus | 0.001 | 0.012 | 0.051 | [-0.052 - 0.054] | 0.000 | 1333 | 0.960 | 0.986 |
| Right Rostral Anterior Cingulate | -0.013 | 0.020 | -0.335 | [-0.098 - 0.072] | 46.962 | 1327 | 0.742 | 0.891 |
| *Right Rostral Middle Frontal* | *-0.065* | *0.011* | *-3.029* | *[-0.111 - -0.019]* | *0.000* | *1302* | *0.009* | *0.110* |
| ***Right Superior Frontal*** | ***-0.096*** | ***0.010*** | ***-4.570*** | ***[-0.141 - -0.051]*** | ***0.000*** | ***1341*** | ***<0.001*** | ***0.030*** |
| Right Superior Parietal | -0.004 | 0.016 | -0.136 | [-0.073 - 0.064] | 10.016 | 1314 | 0.893 | 0.973 |
| Right Superior Temporal | -0.034 | 0.017 | -0.996 | [-0.108 - 0.039] | 0.000 | 1234 | 0.336 | 0.586 |
| Right Supramarginal | -0.069 | 0.020 | -1.692 | [-0.156 - 0.019] | 44.337 | 1223 | 0.113 | 0.403 |
| Right Frontal Pole | -0.004 | 0.013 | -0.153 | [-0.059 - 0.051] | 0.000 | 1365 | 0.881 | 0.973 |
| Right Temporal Pole | -0.001 | 0.008 | -0.037 | [-0.034 - 0.033] | 0.000 | 1225 | 0.971 | 0.986 |
| Right Transverse Temporal | -0.022 | 0.012 | -0.891 | [-0.074 - 0.031] | 0.000 | 1363 | 0.388 | 0.600 |
| Right Insula | -0.060 | 0.015 | -2.022 | [-0.123 - 0.004] | 1.166 | 1363 | 0.063 | 0.283 |
| *Left Thickness* | *-0.055* | *0.013* | *-2.181* | *[-0.108 - -0.001]* | *0.000* | *1370* | *0.047* | *1.000* |
| *Right Thickness* | *-0.056* | *0.012* | *-2.251* | *[-0.109 - -0.003]* | *0.000* | *1370* | *0.041* | *1.000* |
| Left Surface Area | -0.049 | 0.017 | -1.422 | [-0.122 - 0.025] | 5.082 | 1370 | 0.177 | 1.000 |
| Right Surface Area | -0.041 | 0.017 | -1.246 | [-0.112 - 0.030] | 0.000 | 1370 | 0.233 | 1.000 |
| Mean Bankssts | -0.043 | 0.018 | -1.197 | [-0.119 - 0.034] | 0.000 | 1230 | 0.251 | 0.474 |
| Mean Caudal Anterior Cingulate | 0.043 | 0.013 | 1.589 | [-0.015 - 0.100] | 0.000 | 1354 | 0.134 | 0.326 |
| *Mean Caudal Middle Frontal* | *-0.064* | *0.014* | *-2.334* | *[-0.122 - -0.005]* | *0.000* | *1323* | *0.035* | *0.170* |
| Mean Cuneus | -0.029 | 0.021 | -0.697 | [-0.117 - 0.060] | 47.276 | 1243 | 0.497 | 0.705 |
| Mean Entorhinal | 0.017 | 0.012 | 0.733 | [-0.034 - 0.069] | 0.000 | 1137 | 0.476 | 0.705 |
| Mean Fusiform | -0.055 | 0.015 | -1.843 | [-0.119 - 0.009] | 0.000 | 1240 | 0.087 | 0.326 |
| Mean Inferior Parietal | -0.021 | 0.017 | -0.597 | [-0.094 - 0.053] | 0.000 | 1160 | 0.560 | 0.733 |
| Mean Inferior Temporal | -0.025 | 0.017 | -0.745 | [-0.098 - 0.047] | 0.000 | 1227 | 0.468 | 0.705 |
| Mean Isthmus Cingulate | 0.000 | 0.013 | 0.009 | [-0.055 - 0.056] | 0.000 | 1345 | 0.993 | 0.993 |
| Mean Lateral Occipital | -0.038 | 0.018 | -1.080 | [-0.114 - 0.038] | 11.541 | 1237 | 0.299 | 0.534 |
| Mean Lateral Orbitofrontal | -0.048 | 0.015 | -1.637 | [-0.110 - 0.015] | 0.000 | 1327 | 0.124 | 0.326 |
| Mean Lingual | -0.044 | 0.015 | -1.482 | [-0.107 - 0.020] | 0.000 | 1328 | 0.160 | 0.349 |
| Mean Medial Orbitofrontal | -0.050 | 0.017 | -1.430 | [-0.124 - 0.025] | 0.000 | 1298 | 0.175 | 0.349 |
| Mean Middle Temporal | -0.029 | 0.019 | -0.782 | [-0.108 - 0.051] | 8.438 | 1162 | 0.447 | 0.705 |
| Mean Parahippocampal | 0.008 | 0.014 | 0.269 | [-0.053 - 0.068] | 0.000 | 1305 | 0.792 | 0.899 |
| Mean Paracentral | -0.033 | 0.011 | -1.467 | [-0.080 - 0.015] | 0.000 | 1352 | 0.164 | 0.349 |
| *Mean Pars Opercularis* | *-0.089* | *0.017* | *-2.578* | *[-0.162 - -0.015]* | *0.000* | *1273* | *0.022* | *0.124* |
| *Mean Pars Orbitalis* | *-0.078* | *0.011* | *-3.554* | *[-0.125 - -0.031]* | *0.000* | *1301* | *0.003* | *0.052* |
| *Mean Pars Triangularis* | *-0.078* | *0.014* | *-2.864* | *[-0.137 - -0.020]* | *0.000* | *1256* | *0.013* | *0.085* |
| Mean Pericalcarine | -0.002 | 0.016 | -0.062 | [-0.069 - 0.065] | 0.000 | 1340 | 0.951 | 0.980 |
| Mean Postcentral | -0.047 | 0.014 | -1.659 | [-0.108 - 0.014] | 0.000 | 1316 | 0.119 | 0.326 |
| Mean Posterior Cingulate | -0.020 | 0.015 | -0.696 | [-0.083 - 0.042] | 0.000 | 1358 | 0.498 | 0.705 |
| *Mean Precentral* | *-0.066* | *0.011* | *-2.964* | *[-0.114 - -0.018]* | *0.000* | *1318* | *0.010* | *0.085* |
| Mean Precuneus | -0.016 | 0.013 | -0.633 | [-0.072 - 0.039] | 0.000 | 1311 | 0.537 | 0.730 |
| Mean Rostral Anterior Cingulate | 0.015 | 0.016 | 0.491 | [-0.052 - 0.083] | 7.899 | 1309 | 0.631 | 0.795 |
| *Mean Rostral Middle Frontal* | *-0.071* | *0.010* | *-3.365* | *[-0.115 - -0.026]* | *0.000* | *1263* | *0.005* | *0.052* |
| ***Mean Superior Frontal*** | ***-0.094*** | ***0.010*** | ***-4.686*** | ***[-0.136 - -0.051]*** | ***0.000*** | ***1325*** | ***<0.001*** | ***0.012*** |

| ROI | Partial *r* | SE | *t*-Score | 95% CI | *I2* | *N* | *p-value* | FDR |
|---|---|---|---|---|---|---|---|---|
| Mean Superior Parietal | -0.005 | 0.018 | -0.145 | [-0.084 - 0.073] | 20.122 | 1248 | 0.887 | 0.950 |
| Mean Superior Temporal | -0.049 | 0.015 | -1.632 | [-0.112 - 0.015] | 0.000 | 1132 | 0.125 | 0.326 |
| *Mean Supramarginal* | *-0.068* | *0.015* | *-2.209* | *[-0.133 - -0.002]* | *0.000* | *1147* | *0.044* | *0.188* |
| Mean Frontal Pole | -0.007 | 0.013 | -0.267 | [-0.060 - 0.047] | 0.000 | 1363 | 0.793 | 0.899 |
| Mean Temporal Pole | -0.002 | 0.008 | -0.136 | [-0.035 - 0.031] | 0.000 | 1206 | 0.894 | 0.950 |
| Mean Transverse Temporal | -0.009 | 0.012 | -0.379 | [-0.063 - 0.044] | 0.000 | 1356 | 0.710 | 0.862 |
| Mean Insula | -0.053 | 0.015 | -1.780 | [-0.117 - 0.011] | 0.000 | 1358 | 0.097 | 0.326 |

Note: nominally significant results are italicized and FDR significant results are italicized and bolded

Supplementary Table S6 Associations between Cortical Thickness and SANS MAP

| ROI | Partial *r* | SE | *t*-Score | 95% CI | *I2* | *N* | *p-value* | FDR |
|---|---|---|---|---|---|---|---|---|
| Left Bankssts | -0.010 | 0.014 | -0.338 | [-0.070 - 0.051] | 0.000 | 1266 | 0.740 | 0.912 |
| Left Caudal Anterior Cingulate | 0.049 | 0.015 | 1.602 | [-0.016 - 0.113] | 0.000 | 1362 | 0.131 | 0.526 |
| Left Caudal Middle Frontal | -0.049 | 0.015 | -1.671 | [-0.111 - 0.014] | 0.000 | 1351 | 0.117 | 0.497 |
| Left Cuneus | -0.009 | 0.015 | -0.305 | [-0.072 - 0.054] | 0.000 | 1275 | 0.765 | 0.912 |
| Left Entorhinal | -0.001 | 0.011 | -0.056 | [-0.047 - 0.044] | 0.000 | 1278 | 0.956 | 0.970 |
| Left Fusiform | -0.049 | 0.013 | -1.835 | [-0.105 - 0.008] | 0.000 | 1308 | 0.088 | 0.446 |
| Left Inferior Parietal | 0.025 | 0.014 | 0.918 | [-0.033 - 0.083] | 0.000 | 1234 | 0.374 | 0.714 |
| Left Inferior Temporal | -0.023 | 0.013 | -0.886 | [-0.080 - 0.033] | 0.000 | 1275 | 0.391 | 0.714 |
| Left Isthmus Cingulate | 0.017 | 0.016 | 0.531 | [-0.053 - 0.087] | 0.000 | 1361 | 0.604 | 0.838 |
| Left Lateral Occipital | -0.011 | 0.017 | -0.334 | [-0.084 - 0.062] | 0.000 | 1296 | 0.744 | 0.912 |
| Left Lateral Orbitofrontal | -0.042 | 0.010 | -2.009 | [-0.087 - 0.003] | 0.000 | 1364 | 0.064 | 0.397 |
| Left Lingual | -0.017 | 0.014 | -0.602 | [-0.078 - 0.044] | 0.000 | 1339 | 0.557 | 0.838 |
| Left Medial Orbitofrontal | -0.014 | 0.015 | -0.459 | [-0.080 - 0.052] | 0.000 | 1347 | 0.653 | 0.888 |
| Left Middle Temporal | -0.014 | 0.014 | -0.531 | [-0.073 - 0.044] | 0.000 | 1218 | 0.604 | 0.838 |
| Left Parahippocampal | 0.011 | 0.013 | 0.409 | [-0.046 - 0.067] | 0.000 | 1342 | 0.688 | 0.912 |
| Left Paracentral | -0.043 | 0.015 | -1.489 | [-0.106 - 0.019] | 0.000 | 1360 | 0.159 | 0.583 |
| *Left Pars Opercularis* | *-0.092* | *0.019* | *-2.451* | *[-0.171 - -0.012]* | *8.474* | *1323* | *0.028* | *0.230* |
| *Left Pars Orbitalis* | *-0.063* | *0.010* | *-3.020* | *[-0.108 - -0.018]* | *0.000* | *1327* | *0.009* | *0.167* |
| *Left Pars Triangularis* | *-0.062* | *0.012* | *-2.660* | *[-0.112 - -0.012]* | *0.000* | *1304* | *0.019* | *0.230* |
| Left Pericalcarine | -0.008 | 0.025 | -0.164 | [-0.115 - 0.099] | 63.999 | 1350 | 0.872 | 0.926 |
| Left Postcentral | -0.037 | 0.017 | -1.079 | [-0.109 - 0.036] | 0.000 | 1336 | 0.299 | 0.714 |
| Left Posterior Cingulate | 0.015 | 0.013 | 0.575 | [-0.040 - 0.070] | 0.000 | 1363 | 0.574 | 0.838 |
| Left Precentral | -0.052 | 0.019 | -1.351 | [-0.133 - 0.030] | 34.545 | 1344 | 0.198 | 0.674 |
| Left Precuneus | -0.019 | 0.010 | -0.948 | [-0.063 - 0.025] | 0.000 | 1337 | 0.359 | 0.714 |
| Left Rostral Anterior Cingulate | 0.004 | 0.012 | 0.165 | [-0.049 - 0.058] | 0.000 | 1347 | 0.871 | 0.926 |
| *Left Rostral Middle Frontal* | *-0.051* | *0.010* | *-2.408* | *[-0.095 - -0.006]* | *0.000* | *1312* | *0.030* | *0.230* |
| *Left Superior Frontal* | *-0.059* | *0.012* | *-2.348* | *[-0.112 - -0.005]* | *0.000* | *1350* | *0.034* | *0.232* |
| Left Superior Parietal | 0.010 | 0.016 | 0.326 | [-0.058 - 0.079] | 0.000 | 1283 | 0.749 | 0.912 |
| Left Superior Temporal | -0.022 | 0.016 | -0.703 | [-0.090 - 0.046] | 0.000 | 1197 | 0.494 | 0.781 |
| Left Supramarginal | -0.022 | 0.013 | -0.868 | [-0.076 - 0.032] | 0.000 | 1216 | 0.400 | 0.714 |
| Left Frontal Pole | -0.017 | 0.012 | -0.721 | [-0.066 - 0.033] | 0.000 | 1368 | 0.483 | 0.781 |
| Left Temporal Pole | -0.024 | 0.013 | -0.932 | [-0.078 - 0.031] | 0.000 | 1329 | 0.367 | 0.714 |
| Left Transverse Temporal | 0.024 | 0.013 | 0.932 | [-0.032 - 0.081] | 0.000 | 1362 | 0.367 | 0.714 |
| Left Insula | -0.039 | 0.011 | -1.809 | [-0.085 - 0.007] | 0.000 | 1365 | 0.092 | 0.446 |
| Right Bankssts | -0.003 | 0.015 | -0.119 | [-0.066 - 0.059] | 0.000 | 1317 | 0.907 | 0.935 |
| Right Caudal Anterior Cingulate | 0.003 | 0.012 | 0.140 | [-0.046 - 0.053] | 0.000 | 1361 | 0.891 | 0.932 |

| ROI | Partial *r* | SE | *t*-Score | 95% CI | *I2* | *N* | *p-value* | FDR |
|---|---|---|---|---|---|---|---|---|
| *Right Caudal Middle Frontal* | *-0.075* | *0.012* | *-2.986* | *[-0.128 - -0.021]* | *0.000* | *1337* | *0.010* | *0.167* |
| Right Cuneus | 0.020 | 0.012 | 0.826 | [-0.032 - 0.073] | 0.000 | 1316 | 0.423 | 0.719 |
| Right Entorhinal | 0.033 | 0.010 | 1.703 | [-0.009 - 0.075] | 0.000 | 1164 | 0.111 | 0.497 |
| Right Fusiform | -0.034 | 0.014 | -1.246 | [-0.093 - 0.025] | 0.000 | 1281 | 0.233 | 0.692 |
| Right Inferior Parietal | -0.009 | 0.019 | -0.242 | [-0.090 - 0.072] | 0.000 | 1226 | 0.812 | 0.921 |
| Right Inferior Temporal | -0.029 | 0.018 | -0.775 | [-0.107 - 0.050] | 23.828 | 1283 | 0.451 | 0.748 |
| Right Isthmus Cingulate | 0.032 | 0.012 | 1.275 | [-0.022 - 0.085] | 0.000 | 1351 | 0.223 | 0.692 |
| Right Lateral Occipital | -0.029 | 0.017 | -0.865 | [-0.101 - 0.043] | 0.000 | 1287 | 0.402 | 0.714 |
| Right Lateral Orbitofrontal | -0.055 | 0.014 | -1.922 | [-0.115 - 0.006] | 0.000 | 1333 | 0.075 | 0.426 |
| Right Lingual | -0.008 | 0.014 | -0.284 | [-0.068 - 0.052] | 0.000 | 1350 | 0.781 | 0.915 |
| Right Medial Orbitofrontal | -0.032 | 0.018 | -0.910 | [-0.107 - 0.043] | 0.000 | 1316 | 0.378 | 0.714 |
| Right Middle Temporal | -0.039 | 0.023 | -0.851 | [-0.135 - 0.059] | 46.262 | 1255 | 0.409 | 0.714 |
| Right Parahippocampal | 0.033 | 0.014 | 1.216 | [-0.025 - 0.091] | 0.000 | 1322 | 0.244 | 0.692 |
| Right Paracentral | -0.023 | 0.013 | -0.882 | [-0.077 - 0.032] | 0.000 | 1361 | 0.393 | 0.714 |
| Right Pars Opercularis | -0.061 | 0.021 | -1.472 | [-0.148 - 0.028] | 45.063 | 1305 | 0.163 | 0.583 |
| *Right Pars Orbitalis* | *-0.050* | *0.010* | *-2.436* | *[-0.093 - -0.006]* | *0.000* | *1336* | *0.029* | *0.230* |
| Right Pars Triangularis | -0.036 | 0.015 | -1.219 | [-0.100 - 0.027] | 0.000 | 1305 | 0.243 | 0.692 |
| Right Pericalcarine | 0.005 | 0.014 | 0.186 | [-0.054 - 0.064] | 0.000 | 1350 | 0.855 | 0.926 |
| Right Postcentral | -0.020 | 0.019 | -0.541 | [-0.101 - 0.061] | 32.594 | 1342 | 0.597 | 0.838 |
| Right Posterior Cingulate | 0.000 | 0.011 | 0.003 | [-0.046 - 0.047] | 0.000 | 1364 | 0.998 | 0.998 |
| Right Precentral | -0.025 | 0.013 | -0.980 | [-0.080 - 0.030] | 0.000 | 1333 | 0.344 | 0.714 |
| Right Precuneus | 0.023 | 0.010 | 1.156 | [-0.019 - 0.065] | 0.000 | 1333 | 0.267 | 0.714 |
| Right Rostral Anterior Cingulate | -0.011 | 0.016 | -0.360 | [-0.078 - 0.056] | 1.721 | 1327 | 0.724 | 0.912 |
| *Right Rostral Middle Frontal* | *-0.062* | *0.010* | *-3.137* | *[-0.105 - -0.020]* | *0.000* | *1302* | *0.007* | *0.167* |
| *Right Superior Frontal* | *-0.083* | *0.013* | *-3.301* | *[-0.136 - -0.029]* | *0.000* | *1341* | *0.005* | *0.167* |
| Right Superior Parietal | 0.008 | 0.016 | 0.254 | [-0.060 - 0.076] | 0.000 | 1314 | 0.803 | 0.921 |
| Right Superior Temporal | -0.005 | 0.014 | -0.167 | [-0.065 - 0.055] | 0.000 | 1234 | 0.870 | 0.926 |
| Right Supramarginal | -0.047 | 0.025 | -0.930 | [-0.154 - 0.061] | 66.345 | 1223 | 0.368 | 0.714 |
| Right Frontal Pole | 0.009 | 0.015 | 0.312 | [-0.054 - 0.072] | 0.000 | 1365 | 0.760 | 0.912 |
| Right Temporal Pole | 0.021 | 0.011 | 0.971 | [-0.025 - 0.067] | 0.000 | 1225 | 0.348 | 0.714 |
| Right Transverse Temporal | -0.013 | 0.011 | -0.588 | [-0.062 - 0.035] | 0.000 | 1363 | 0.566 | 0.838 |
| *Right Insula* | *-0.063* | *0.012* | *-2.552* | *[-0.116 - -0.010]* | *0.000* | *1363* | *0.023* | *0.230* |
| Left Thickness | -0.037 | 0.013 | -1.390 | [-0.094 - 0.020] | 0.000 | 1370 | 0.186 | 1.000 |
| Right Thickness | -0.037 | 0.014 | -1.322 | [-0.096 - 0.023] | 0.000 | 1370 | 0.207 | 1.000 |
| Left Surface Area | -0.056 | 0.017 | -1.606 | [-0.130 - 0.019] | 0.000 | 1370 | 0.130 | 1.000 |
| Right Surface Area | -0.054 | 0.016 | -1.660 | [-0.123 - 0.016] | 0.000 | 1370 | 0.119 | 1.000 |
| Mean Bankssts | -0.010 | 0.014 | -0.382 | [-0.069 - 0.048] | 0.000 | 1230 | 0.708 | 0.956 |
| Mean Caudal Anterior Cingulate | 0.039 | 0.013 | 1.443 | [-0.019 - 0.097] | 0.000 | 1354 | 0.171 | 0.576 |
| *Mean Caudal Middle Frontal* | *-0.067* | *0.014* | *-2.388* | *[-0.126 - -0.007]* | *0.000* | *1323* | *0.032* | *0.269* |
| Mean Cuneus | 0.006 | 0.015 | 0.213 | [-0.057 - 0.069] | 0.000 | 1243 | 0.835 | 0.987 |
| Mean Entorhinal | 0.027 | 0.013 | 1.040 | [-0.028 - 0.082] | 0.000 | 1137 | 0.316 | 0.722 |
| Mean Fusiform | -0.039 | 0.014 | -1.390 | [-0.100 - 0.021] | 0.000 | 1240 | 0.186 | 0.576 |
| Mean Inferior Parietal | 0.003 | 0.017 | 0.092 | [-0.068 - 0.074] | 0.000 | 1160 | 0.928 | 0.987 |
| Mean Inferior Temporal | -0.026 | 0.017 | -0.770 | [-0.098 - 0.046] | 0.000 | 1227 | 0.454 | 0.812 |
| Mean Isthmus Cingulate | 0.029 | 0.015 | 0.988 | [-0.034 - 0.092] | 0.000 | 1345 | 0.340 | 0.722 |
| Mean Lateral Occipital | -0.019 | 0.018 | -0.518 | [-0.097 - 0.060] | 0.000 | 1237 | 0.613 | 0.947 |
| Mean Lateral Orbitofrontal | -0.048 | 0.012 | -2.032 | [-0.099 - 0.003] | 0.000 | 1327 | 0.062 | 0.269 |
| Mean Lingual | -0.017 | 0.015 | -0.586 | [-0.080 - 0.046] | 0.000 | 1328 | 0.567 | 0.918 |
| Mean Medial Orbitofrontal | -0.026 | 0.017 | -0.781 | [-0.099 - 0.046] | 0.000 | 1298 | 0.448 | 0.812 |
| Mean Middle Temporal | -0.026 | 0.021 | -0.622 | [-0.114 - 0.063] | 29.114 | 1162 | 0.544 | 0.918 |
| Mean Parahippocampal | 0.027 | 0.014 | 0.941 | [-0.034 - 0.087] | 0.000 | 1305 | 0.363 | 0.725 |

| ROI | Partial *r* | SE | *t*-Score | 95% CI | *I2* | *N* | *p-value* | FDR |
|---|---|---|---|---|---|---|---|---|
| Mean Paracentral | -0.034 | 0.014 | -1.215 | [-0.092 - 0.026] | 0.000 | 1352 | 0.244 | 0.660 |
| Mean Pars Opercularis | -0.086 | 0.021 | -2.017 | [-0.177 - 0.005] | 39.194 | 1273 | 0.063 | 0.269 |
| *Mean Pars Orbitalis* | *-0.077* | *0.010* | *-3.894* | *[-0.118 - -0.034]* | *0.000* | *1301* | *0.002* | *0.055* |
| Mean Pars Triangularis | -0.059 | 0.014 | -2.144 | [-0.118 - 0.000] | 0.000 | 1256 | 0.050 | 0.269 |
| Mean Pericalcarine | 0.013 | 0.018 | 0.374 | [-0.064 - 0.090] | 0.000 | 1340 | 0.714 | 0.956 |
| Mean Postcentral | -0.042 | 0.017 | -1.194 | [-0.116 - 0.033] | 0.000 | 1316 | 0.252 | 0.660 |
| Mean Posterior Cingulate | 0.010 | 0.012 | 0.417 | [-0.042 - 0.062] | 0.000 | 1358 | 0.683 | 0.956 |
| Mean Precentral | -0.046 | 0.015 | -1.569 | [-0.109 - 0.017] | 0.000 | 1318 | 0.139 | 0.525 |
| Mean Precuneus | 0.003 | 0.010 | 0.142 | [-0.039 - 0.044] | 0.000 | 1311 | 0.889 | 0.987 |
| Mean Rostral Anterior Cingulate | 0.000 | 0.015 | -0.016 | [-0.064 - 0.063] | 0.000 | 1309 | 0.987 | 0.987 |
| *Mean Rostral Middle Frontal* | *-0.062* | *0.010* | *-3.111* | *[-0.104 - -0.019]* | *0.000* | *1263* | *0.008* | *0.087* |
| *Mean Superior Frontal* | *-0.075* | *0.012* | *-3.133* | *[-0.126 - -0.024]* | *0.000* | *1325* | *0.007* | *0.087* |
| Mean Superior Parietal | 0.008 | 0.016 | 0.247 | [-0.062 - 0.078] | 0.000 | 1248 | 0.808 | 0.987 |
| Mean Superior Temporal | -0.011 | 0.015 | -0.351 | [-0.075 - 0.054] | 0.000 | 1132 | 0.731 | 0.956 |
| Mean Supramarginal | -0.035 | 0.017 | -0.991 | [-0.109 - 0.040] | 0.000 | 1147 | 0.338 | 0.722 |
| Mean Frontal Pole | 0.002 | 0.014 | 0.077 | [-0.060 - 0.064] | 0.000 | 1363 | 0.940 | 0.987 |
| Mean Temporal Pole | -0.001 | 0.010 | -0.047 | [-0.042 - 0.040] | 0.000 | 1206 | 0.963 | 0.987 |
| Mean Transverse Temporal | 0.005 | 0.012 | 0.196 | [-0.047 - 0.057] | 0.000 | 1356 | 0.848 | 0.987 |
| *Mean Insula* | *-0.054* | *0.012* | *-2.245* | *[-0.106 - -0.002]* | *0.000* | *1358* | *0.041* | *0.269* |

Note: nominally significant results are italicized and FDR significant results are italicized and bolded

Supplementary Table S7 Associations between Cortical Thickness and SANS Anhedonia (Factor 1)

| ROI | Partial *r* | SE | *t*-Score | 95% CI | *I2* | *N* | *p-value* | FDR |
|---|---|---|---|---|---|---|---|---|
| Left Bankssts | 0.014 | 0.014 | 0.485 | [-0.048 - 0.076] | 0.000 | 1266 | 0.635 | 0.948 |
| Left Caudal Anterior Cingulate | 0.056 | 0.014 | 1.968 | [-0.005 - 0.116] | 0.000 | 1362 | 0.069 | 0.507 |
| Left Caudal Middle Frontal | -0.024 | 0.013 | -0.896 | [-0.081 - 0.033] | 0.000 | 1351 | 0.385 | 0.784 |
| Left Cuneus | -0.003 | 0.011 | -0.120 | [-0.051 - 0.045] | 0.000 | 1275 | 0.906 | 0.948 |
| Left Entorhinal | -0.013 | 0.007 | -0.969 | [-0.042 - 0.016] | 0.000 | 1278 | 0.349 | 0.784 |
| Left Fusiform | -0.031 | 0.007 | -2.101 | [-0.062 - 0.001] | 0.000 | 1308 | 0.054 | 0.507 |
| Left Inferior Parietal | 0.045 | 0.014 | 1.570 | [-0.017 - 0.107] | 0.000 | 1234 | 0.139 | 0.551 |
| Left Inferior Temporal | -0.025 | 0.013 | -0.964 | [-0.080 - 0.030] | 0.000 | 1275 | 0.351 | 0.784 |
| Left Isthmus Cingulate | 0.016 | 0.017 | 0.462 | [-0.058 - 0.089] | 0.000 | 1361 | 0.651 | 0.948 |
| Left Lateral Occipital | 0.010 | 0.016 | 0.302 | [-0.059 - 0.079] | 0.000 | 1296 | 0.767 | 0.948 |
| *Left Lateral Orbitofrontal* | *-0.046* | *0.010* | *-2.356* | *[-0.087 - -0.004]* | *0.000* | *1364* | *0.034* | *0.507* |
| Left Lingual | 0.013 | 0.012 | 0.544 | [-0.039 - 0.065] | 0.000 | 1339 | 0.595 | 0.948 |
| Left Medial Orbitofrontal | -0.032 | 0.015 | -1.071 | [-0.095 - 0.032] | 0.000 | 1347 | 0.302 | 0.784 |
| Left Middle Temporal | -0.005 | 0.012 | -0.195 | [-0.057 - 0.047] | 0.000 | 1218 | 0.848 | 0.948 |
| Left Parahippocampal | -0.006 | 0.013 | -0.253 | [-0.061 - 0.049] | 0.000 | 1342 | 0.804 | 0.948 |
| Left Paracentral | -0.006 | 0.022 | -0.127 | [-0.098 - 0.087] | 47.282 | 1360 | 0.901 | 0.948 |
| *Left Pars Opercularis* | *-0.088* | *0.015* | *-3.015* | *[-0.150 - -0.026]* | *0.000* | *1323* | *0.009* | *0.290* |
| *Left Pars Orbitalis* | *-0.070* | *0.009* | *-3.802* | *[-0.109 - -0.031]* | *0.000* | *1327* | *0.002* | *0.132* |
| *Left Pars Triangularis* | *-0.057* | *0.010* | *-2.853* | *[-0.100 - -0.014]* | *0.000* | *1304* | *0.013* | *0.290* |
| Left Pericalcarine | 0.009 | 0.020 | 0.224 | [-0.078 - 0.096] | 42.040 | 1350 | 0.826 | 0.948 |
| Left Postcentral | -0.035 | 0.019 | -0.933 | [-0.114 - 0.045] | 0.000 | 1336 | 0.367 | 0.784 |
| Left Posterior Cingulate | 0.012 | 0.016 | 0.370 | [-0.056 - 0.079] | 0.000 | 1363 | 0.717 | 0.948 |
| Left Precentral | -0.006 | 0.016 | -0.186 | [-0.075 - 0.063] | 0.000 | 1344 | 0.855 | 0.948 |
| Left Precuneus | 0.001 | 0.012 | 0.051 | [-0.051 - 0.053] | 0.000 | 1337 | 0.960 | 0.974 |
| Left Rostral Anterior Cingulate | 0.015 | 0.015 | 0.475 | [-0.052 - 0.081] | 7.451 | 1347 | 0.642 | 0.948 |
| Left Rostral Middle Frontal | -0.044 | 0.011 | -2.063 | [-0.090 - 0.002] | 0.000 | 1312 | 0.058 | 0.507 |
| Left Superior Frontal | -0.041 | 0.013 | -1.589 | [-0.095 - 0.014] | 0.000 | 1350 | 0.134 | 0.551 |

| ROI | Partial *r* | SE | *t*-Score | 95% CI | *I2* | *N* | *p-value* | FDR |
|---|---|---|---|---|---|---|---|---|
| Left Superior Parietal | 0.015 | 0.016 | 0.480 | [-0.053 - 0.083] | 0.000 | 1283 | 0.639 | 0.948 |
| Left Superior Temporal | 0.009 | 0.013 | 0.351 | [-0.047 - 0.065] | 0.000 | 1197 | 0.731 | 0.948 |
| Left Supramarginal | 0.010 | 0.015 | 0.332 | [-0.055 - 0.075] | 0.026 | 1216 | 0.745 | 0.948 |
| Left Frontal Pole | -0.008 | 0.013 | -0.300 | [-0.063 - 0.048] | 0.000 | 1368 | 0.768 | 0.948 |
| Left Temporal Pole | -0.041 | 0.014 | -1.507 | [-0.099 - 0.017] | 0.000 | 1329 | 0.154 | 0.551 |
| Left Transverse Temporal | 0.020 | 0.007 | 1.351 | [-0.011 - 0.051] | 0.000 | 1362 | 0.198 | 0.673 |
| Left Insula | -0.043 | 0.014 | -1.517 | [-0.104 - 0.018] | 0.000 | 1365 | 0.152 | 0.551 |
| Right Bankssts | 0.014 | 0.012 | 0.555 | [-0.039 - 0.066] | 0.000 | 1317 | 0.588 | 0.948 |
| Right Caudal Anterior Cingulate | -0.011 | 0.011 | -0.519 | [-0.058 - 0.035] | 0.000 | 1361 | 0.612 | 0.948 |
| Right Caudal Middle Frontal | -0.052 | 0.014 | -1.863 | [-0.112 - 0.008] | 0.000 | 1337 | 0.084 | 0.507 |
| Right Cuneus | 0.023 | 0.013 | 0.915 | [-0.031 - 0.078] | 0.000 | 1316 | 0.375 | 0.784 |
| Right Entorhinal | 0.017 | 0.010 | 0.840 | [-0.027 - 0.062] | 0.000 | 1164 | 0.415 | 0.784 |
| Right Fusiform | -0.018 | 0.008 | -1.138 | [-0.053 - 0.016] | 0.000 | 1281 | 0.274 | 0.784 |
| Right Inferior Parietal | 0.023 | 0.017 | 0.709 | [-0.047 - 0.094] | 0.000 | 1226 | 0.490 | 0.877 |
| Right Inferior Temporal | 0.007 | 0.014 | 0.237 | [-0.054 - 0.067] | 0.000 | 1283 | 0.816 | 0.948 |
| Right Isthmus Cingulate | 0.048 | 0.014 | 1.737 | [-0.011 - 0.108] | 0.000 | 1351 | 0.104 | 0.507 |
| Right Lateral Occipital | 0.002 | 0.016 | 0.064 | [-0.068 - 0.072] | 11.604 | 1287 | 0.950 | 0.974 |
| Right Lateral Orbitofrontal | -0.051 | 0.014 | -1.860 | [-0.109 - 0.008] | 0.537 | 1333 | 0.084 | 0.507 |
| Right Lingual | 0.028 | 0.015 | 0.922 | [-0.037 - 0.093] | 0.000 | 1350 | 0.372 | 0.784 |
| Right Medial Orbitofrontal | -0.045 | 0.017 | -1.314 | [-0.117 - 0.028] | 28.664 | 1316 | 0.210 | 0.680 |
| Right Middle Temporal | 0.004 | 0.017 | 0.120 | [-0.069 - 0.078] | 0.000 | 1255 | 0.906 | 0.948 |
| Right Parahippocampal | 0.021 | 0.014 | 0.768 | [-0.037 - 0.079] | 0.000 | 1322 | 0.455 | 0.837 |
| Right Paracentral | -0.011 | 0.016 | -0.326 | [-0.080 - 0.059] | 0.000 | 1361 | 0.749 | 0.948 |
| Right Pars Opercularis | -0.046 | 0.023 | -1.000 | [-0.145 - 0.053] | 60.735 | 1305 | 0.334 | 0.784 |
| *Right Pars Orbitalis* | *-0.055* | *0.012* | *-2.243* | *[-0.107 - -0.002]* | *0.000* | *1336* | *0.042* | *0.507* |
| Right Pars Triangularis | -0.023 | 0.018 | -0.651 | [-0.100 - 0.054] | 0.000 | 1305 | 0.526 | 0.917 |
| Right Pericalcarine | 0.028 | 0.013 | 1.065 | [-0.029 - 0.085] | 0.000 | 1350 | 0.305 | 0.784 |
| Right Postcentral | -0.029 | 0.017 | -0.857 | [-0.100 - 0.043] | 9.654 | 1342 | 0.406 | 0.784 |
| Right Posterior Cingulate | 0.008 | 0.015 | 0.269 | [-0.057 - 0.074] | 0.000 | 1364 | 0.792 | 0.948 |
| Right Precentral | 0.004 | 0.016 | 0.142 | [-0.062 - 0.071] | 0.000 | 1333 | 0.889 | 0.948 |
| Right Precuneus | 0.043 | 0.012 | 1.758 | [-0.009 - 0.094] | 0.000 | 1333 | 0.101 | 0.507 |
| Right Rostral Anterior Cingulate | 0.005 | 0.016 | 0.144 | [-0.063 - 0.072] | 0.000 | 1327 | 0.887 | 0.948 |
| Right Rostral Middle Frontal | -0.043 | 0.013 | -1.646 | [-0.099 - 0.013] | 0.000 | 1302 | 0.122 | 0.551 |
| Right Superior Frontal | -0.058 | 0.015 | -1.902 | [-0.124 - 0.007] | 0.000 | 1341 | 0.078 | 0.507 |
| Right Superior Parietal | 0.033 | 0.016 | 1.045 | [-0.035 - 0.102] | 0.000 | 1314 | 0.314 | 0.784 |
| Right Superior Temporal | 0.018 | 0.010 | 0.863 | [-0.027 - 0.062] | 0.000 | 1234 | 0.403 | 0.784 |
| Right Supramarginal | 0.000 | 0.016 | 0.003 | [-0.069 - 0.070] | 0.000 | 1223 | 0.997 | 0.997 |
| Right Frontal Pole | -0.035 | 0.014 | -1.233 | [-0.097 - 0.026] | 0.000 | 1365 | 0.238 | 0.735 |
| Right Temporal Pole | 0.007 | 0.011 | 0.326 | [-0.039 - 0.053] | 0.000 | 1225 | 0.749 | 0.948 |
| Right Transverse Temporal | 0.005 | 0.012 | 0.219 | [-0.047 - 0.058] | 0.000 | 1363 | 0.830 | 0.948 |
| Right Insula | -0.046 | 0.013 | -1.802 | [-0.100 - 0.009] | 0.000 | 1363 | 0.093 | 0.507 |
| Left Thickness | -0.018 | 0.013 | -0.675 | [-0.073 - 0.038] | 0.000 | 1370 | 0.511 | 1.000 |
| Right Thickness | -0.012 | 0.014 | -0.410 | [-0.072 - 0.049] | 0.000 | 1370 | 0.688 | 1.000 |
| Left Surface Area | -0.046 | 0.015 | -1.576 | [-0.108 - 0.017] | 0.000 | 1370 | 0.137 | 1.000 |
| Right Surface Area | -0.041 | 0.013 | -1.524 | [-0.099 - 0.017] | 0.000 | 1370 | 0.150 | 1.000 |
| Mean Bankssts | 0.013 | 0.013 | 0.496 | [-0.042 - 0.067] | 0.000 | 1230 | 0.628 | 0.788 |
| Mean Caudal Anterior Cingulate | 0.035 | 0.011 | 1.578 | [-0.013 - 0.083] | 0.000 | 1354 | 0.137 | 0.583 |
| Mean Caudal Middle Frontal | -0.040 | 0.014 | -1.456 | [-0.099 - 0.019] | 0.000 | 1323 | 0.167 | 0.624 |
| Mean Cuneus | 0.010 | 0.012 | 0.432 | [-0.041 - 0.061] | 0.000 | 1243 | 0.672 | 0.788 |
| Mean Entorhinal | 0.013 | 0.010 | 0.613 | [-0.031 - 0.057] | 0.000 | 1137 | 0.550 | 0.779 |
| Mean Fusiform | -0.018 | 0.007 | -1.387 | [-0.047 - 0.010] | 0.000 | 1240 | 0.187 | 0.624 |

| ROI | Partial *r* | SE | *t*-Score | 95% CI | *I2* | *N* | *p-value* | FDR |
|---|---|---|---|---|---|---|---|---|
| Mean Inferior Parietal | 0.037 | 0.015 | 1.222 | [-0.028 - 0.103] | 0.000 | 1160 | 0.242 | 0.670 |
| Mean Inferior Temporal | -0.012 | 0.014 | -0.439 | [-0.071 - 0.047] | 0.000 | 1227 | 0.667 | 0.788 |
| Mean Isthmus Cingulate | 0.037 | 0.016 | 1.133 | [-0.033 - 0.105] | 0.000 | 1345 | 0.276 | 0.670 |
| Mean Lateral Occipital | 0.010 | 0.017 | 0.304 | [-0.063 - 0.083] | 0.000 | 1237 | 0.766 | 0.840 |
| Mean Lateral Orbitofrontal | -0.046 | 0.011 | -2.069 | [-0.094 - 0.002] | 0.000 | 1327 | 0.057 | 0.583 |
| Mean Lingual | 0.020 | 0.014 | 0.720 | [-0.040 - 0.081] | 0.000 | 1328 | 0.484 | 0.779 |
| Mean Medial Orbitofrontal | -0.041 | 0.015 | -1.340 | [-0.107 - 0.025] | 0.000 | 1298 | 0.202 | 0.624 |
| Mean Middle Temporal | 0.003 | 0.015 | 0.112 | [-0.063 - 0.070] | 0.000 | 1162 | 0.912 | 0.912 |
| Mean Parahippocampal | 0.009 | 0.012 | 0.370 | [-0.043 - 0.060] | 0.000 | 1305 | 0.717 | 0.813 |
| Mean Paracentral | -0.022 | 0.017 | -0.652 | [-0.093 - 0.050] | 0.000 | 1352 | 0.525 | 0.779 |
| Mean Pars Opercularis | -0.068 | 0.020 | -1.727 | [-0.152 - 0.017] | 26.002 | 1273 | 0.106 | 0.583 |
| *Mean Pars Orbitalis* | *-0.082* | *0.012* | *-3.512* | *[-0.131 - -0.032]* | *0.000* | *1301* | *0.003* | *0.117* |
| Mean Pars Triangularis | -0.048 | 0.015 | -1.657 | [-0.111 - 0.014] | 0.000 | 1256 | 0.120 | 0.583 |
| Mean Pericalcarine | 0.030 | 0.016 | 0.971 | [-0.036 - 0.097] | 0.000 | 1340 | 0.348 | 0.740 |
| Mean Postcentral | -0.039 | 0.018 | -1.066 | [-0.117 - 0.040] | 0.000 | 1316 | 0.305 | 0.690 |
| Mean Posterior Cingulate | 0.015 | 0.016 | 0.461 | [-0.054 - 0.083] | 0.000 | 1358 | 0.652 | 0.788 |
| Mean Precentral | -0.007 | 0.016 | -0.232 | [-0.075 - 0.060] | 0.000 | 1318 | 0.820 | 0.869 |
| Mean Precuneus | 0.027 | 0.012 | 1.165 | [-0.023 - 0.077] | 0.000 | 1311 | 0.263 | 0.670 |
| Mean Rostral Anterior Cingulate | 0.016 | 0.015 | 0.544 | [-0.048 - 0.081] | 0.000 | 1309 | 0.595 | 0.788 |
| Mean Rostral Middle Frontal | -0.046 | 0.012 | -1.832 | [-0.099 - 0.008] | 0.000 | 1263 | 0.088 | 0.583 |
| Mean Superior Frontal | -0.052 | 0.013 | -1.946 | [-0.108 - 0.005] | 0.000 | 1325 | 0.072 | 0.583 |
| Mean Superior Parietal | 0.027 | 0.016 | 0.844 | [-0.042 - 0.096] | 0.000 | 1248 | 0.413 | 0.779 |
| Mean Superior Temporal | 0.020 | 0.012 | 0.793 | [-0.034 - 0.073] | 0.000 | 1132 | 0.441 | 0.779 |
| Mean Supramarginal | 0.007 | 0.017 | 0.201 | [-0.066 - 0.080] | 0.000 | 1147 | 0.843 | 0.869 |
| Mean Frontal Pole | -0.018 | 0.015 | -0.635 | [-0.081 - 0.044] | 0.000 | 1363 | 0.536 | 0.779 |
| Mean Temporal Pole | -0.016 | 0.011 | -0.754 | [-0.062 - 0.030] | 0.000 | 1206 | 0.463 | 0.779 |
| Mean Transverse Temporal | 0.014 | 0.009 | 0.772 | [-0.025 - 0.054] | 0.000 | 1356 | 0.453 | 0.779 |
| Mean Insula | -0.046 | 0.014 | -1.576 | [-0.107 - 0.016] | 0.000 | 1358 | 0.137 | 0.583 |

Note: nominally significant results are italicized and FDR significant results are italicized and bolded

Supplementary Table S8 Associations between Cortical Thickness and SANS Asociality (Factor 2)

| ROI | Partial *r* | SE | *t*-Score | 95% CI | *I2* | *N* | *p-value* | FDR |
|---|---|---|---|---|---|---|---|---|
| Left Bankssts | 0.005 | 0.013 | 0.171 | [-0.052 - 0.061] | 0.000 | 1266 | 0.866 | 0.944 |
| Left Caudal Anterior Cingulate | 0.047 | 0.015 | 1.562 | [-0.017 - 0.110] | 0.000 | 1362 | 0.141 | 0.742 |
| Left Caudal Middle Frontal | -0.047 | 0.015 | -1.556 | [-0.111 - 0.018] | 0.000 | 1351 | 0.142 | 0.742 |
| Left Cuneus | -0.016 | 0.016 | -0.511 | [-0.083 - 0.051] | 0.000 | 1275 | 0.617 | 0.933 |
| Left Entorhinal | 0.005 | 0.014 | 0.177 | [-0.054 - 0.064] | 0.000 | 1278 | 0.862 | 0.944 |
| Left Fusiform | -0.039 | 0.015 | -1.344 | [-0.101 - 0.023] | 0.000 | 1308 | 0.200 | 0.813 |
| Left Inferior Parietal | -0.004 | 0.012 | -0.166 | [-0.055 - 0.047] | 0.000 | 1234 | 0.870 | 0.944 |
| Left Inferior Temporal | -0.013 | 0.010 | -0.606 | [-0.057 - 0.032] | 0.000 | 1275 | 0.554 | 0.925 |
| Left Isthmus Cingulate | 0.022 | 0.017 | 0.634 | [-0.052 - 0.096] | 0.000 | 1361 | 0.536 | 0.925 |
| Left Lateral Occipital | -0.021 | 0.017 | -0.600 | [-0.094 - 0.053] | 0.000 | 1296 | 0.558 | 0.925 |
| Left Lateral Orbitofrontal | -0.027 | 0.012 | -1.118 | [-0.080 - 0.025] | 0.000 | 1364 | 0.282 | 0.813 |
| Left Lingual | -0.018 | 0.016 | -0.574 | [-0.086 - 0.050] | 0.000 | 1339 | 0.575 | 0.927 |
| Left Medial Orbitofrontal | -0.005 | 0.015 | -0.157 | [-0.067 - 0.058] | 0.000 | 1347 | 0.877 | 0.944 |
| Left Middle Temporal | -0.007 | 0.013 | -0.276 | [-0.060 - 0.047] | 0.000 | 1218 | 0.787 | 0.944 |
| Left Parahippocampal | 0.021 | 0.013 | 0.802 | [-0.036 - 0.078] | 0.000 | 1342 | 0.436 | 0.905 |
| Left Paracentral | -0.029 | 0.012 | -1.191 | [-0.080 - 0.023] | 0.000 | 1360 | 0.253 | 0.813 |
| Left Pars Opercularis | -0.075 | 0.018 | -2.024 | [-0.154 - 0.004] | 26.678 | 1323 | 0.062 | 0.609 |
| Left Pars Orbitalis | -0.053 | 0.013 | -2.022 | [-0.108 - 0.003] | 0.000 | 1327 | 0.063 | 0.609 |

| ROI | Partial *r* | SE | *t*-Score | 95% CI | *I2* | *N* | *p-value* | FDR |
|---|---|---|---|---|---|---|---|---|
| Left Pars Triangularis | -0.032 | 0.014 | -1.097 | [-0.094 - 0.030] | 0.000 | 1304 | 0.291 | 0.813 |
| Left Pericalcarine | -0.016 | 0.027 | -0.288 | [-0.130 - 0.100] | 70.860 | 1350 | 0.777 | 0.944 |
| Left Postcentral | -0.013 | 0.016 | -0.388 | [-0.082 - 0.057] | 0.000 | 1336 | 0.704 | 0.944 |
| Left Posterior Cingulate | 0.014 | 0.008 | 0.922 | [-0.019 - 0.048] | 0.000 | 1363 | 0.372 | 0.845 |
| Left Precentral | -0.038 | 0.017 | -1.132 | [-0.109 - 0.034] | 0.000 | 1344 | 0.277 | 0.813 |
| Left Precuneus | -0.021 | 0.012 | -0.920 | [-0.071 - 0.028] | 0.000 | 1337 | 0.373 | 0.845 |
| Left Rostral Anterior Cingulate | 0.000 | 0.014 | 0.002 | [-0.058 - 0.058] | 0.000 | 1347 | 0.998 | 0.998 |
| Left Rostral Middle Frontal | -0.044 | 0.012 | -1.874 | [-0.094 - 0.006] | 0.000 | 1312 | 0.082 | 0.619 |
| Left Superior Frontal | -0.050 | 0.013 | -1.928 | [-0.106 - 0.006] | 0.000 | 1350 | 0.074 | 0.619 |
| Left Superior Parietal | 0.007 | 0.015 | 0.237 | [-0.056 - 0.070] | 0.000 | 1283 | 0.816 | 0.944 |
| Left Superior Temporal | -0.012 | 0.016 | -0.381 | [-0.080 - 0.056] | 0.000 | 1197 | 0.709 | 0.944 |
| Left Supramarginal | -0.022 | 0.011 | -0.988 | [-0.069 - 0.026] | 0.000 | 1216 | 0.340 | 0.845 |
| Left Frontal Pole | 0.001 | 0.013 | 0.041 | [-0.054 - 0.056] | 0.000 | 1368 | 0.968 | 0.982 |
| Left Temporal Pole | 0.019 | 0.015 | 0.655 | [-0.043 - 0.081] | 0.000 | 1329 | 0.523 | 0.925 |
| Left Transverse Temporal | 0.032 | 0.015 | 1.093 | [-0.031 - 0.094] | 0.000 | 1362 | 0.293 | 0.813 |
| Left Insula | -0.004 | 0.010 | -0.210 | [-0.049 - 0.040] | 0.000 | 1365 | 0.837 | 0.944 |
| Right Bankssts | -0.009 | 0.014 | -0.302 | [-0.071 - 0.053] | 0.000 | 1317 | 0.767 | 0.944 |
| Right Caudal Anterior Cingulate | -0.007 | 0.012 | -0.302 | [-0.058 - 0.044] | 0.000 | 1361 | 0.767 | 0.944 |
| Right Caudal Middle Frontal | -0.061 | 0.015 | -2.063 | [-0.124 - 0.002] | 0.000 | 1337 | 0.058 | 0.609 |
| Right Cuneus | 0.012 | 0.015 | 0.378 | [-0.054 - 0.078] | 0.000 | 1316 | 0.711 | 0.944 |
| Right Entorhinal | 0.055 | 0.016 | 1.733 | [-0.013 - 0.122] | 0.000 | 1164 | 0.105 | 0.649 |
| Right Fusiform | -0.028 | 0.015 | -0.953 | [-0.091 - 0.035] | 0.000 | 1281 | 0.357 | 0.845 |
| Right Inferior Parietal | -0.020 | 0.018 | -0.557 | [-0.096 - 0.056] | 0.000 | 1226 | 0.586 | 0.927 |
| Right Inferior Temporal | -0.023 | 0.017 | -0.680 | [-0.095 - 0.049] | 0.000 | 1283 | 0.507 | 0.925 |
| Right Isthmus Cingulate | 0.026 | 0.012 | 1.079 | [-0.025 - 0.077] | 0.000 | 1351 | 0.299 | 0.813 |
| Right Lateral Occipital | -0.043 | 0.017 | -1.248 | [-0.118 - 0.031] | 0.000 | 1287 | 0.232 | 0.813 |
| Right Lateral Orbitofrontal | -0.050 | 0.014 | -1.776 | [-0.110 - 0.010] | 0.000 | 1333 | 0.097 | 0.649 |
| Right Lingual | -0.011 | 0.013 | -0.400 | [-0.068 - 0.046] | 0.000 | 1350 | 0.695 | 0.944 |
| Right Medial Orbitofrontal | -0.015 | 0.015 | -0.490 | [-0.081 - 0.051] | 0.000 | 1316 | 0.631 | 0.933 |
| Right Middle Temporal | -0.034 | 0.022 | -0.772 | [-0.127 - 0.060] | 48.321 | 1255 | 0.453 | 0.906 |
| Right Parahippocampal | 0.047 | 0.016 | 1.462 | [-0.022 - 0.114] | 0.410 | 1322 | 0.166 | 0.806 |
| Right Paracentral | -0.018 | 0.011 | -0.797 | [-0.065 - 0.030] | 0.000 | 1361 | 0.439 | 0.905 |
| Right Pars Opercularis | -0.040 | 0.017 | -1.150 | [-0.115 - 0.035] | 0.000 | 1305 | 0.270 | 0.813 |
| Right Pars Orbitalis | -0.042 | 0.010 | -2.064 | [-0.086 - 0.002] | 0.000 | 1336 | 0.058 | 0.609 |
| Right Pars Triangularis | -0.041 | 0.015 | -1.406 | [-0.103 - 0.021] | 0.000 | 1305 | 0.182 | 0.813 |
| Right Pericalcarine | 0.020 | 0.015 | 0.650 | [-0.045 - 0.085] | 0.000 | 1350 | 0.526 | 0.925 |
| Right Postcentral | -0.004 | 0.016 | -0.129 | [-0.074 - 0.066] | 0.000 | 1342 | 0.899 | 0.944 |
| Right Posterior Cingulate | 0.004 | 0.009 | 0.220 | [-0.033 - 0.041] | 0.000 | 1364 | 0.829 | 0.944 |
| Right Precentral | -0.023 | 0.014 | -0.816 | [-0.083 - 0.037] | 0.000 | 1333 | 0.428 | 0.905 |
| Right Precuneus | 0.014 | 0.010 | 0.686 | [-0.030 - 0.057] | 0.000 | 1333 | 0.504 | 0.925 |
| Right Rostral Anterior Cingulate | -0.008 | 0.015 | -0.255 | [-0.073 - 0.058] | 0.000 | 1327 | 0.802 | 0.944 |
| *Right Rostral Middle Frontal* | *-0.066* | *0.011* | *-3.068* | *[-0.112 - -0.020]* | *0.000* | *1302* | *0.008* | *0.538* |
| *Right Superior Frontal* | *-0.080* | *0.014* | *-2.745* | *[-0.141 - -0.017]* | *5.541* | *1341* | *0.016* | *0.538* |
| Right Superior Parietal | -0.008 | 0.017 | -0.218 | [-0.082 - 0.067] | 0.000 | 1314 | 0.831 | 0.944 |
| Right Superior Temporal | 0.003 | 0.016 | 0.107 | [-0.064 - 0.070] | 0.000 | 1234 | 0.916 | 0.944 |
| Right Supramarginal | -0.019 | 0.020 | -0.495 | [-0.103 - 0.064] | 0.000 | 1223 | 0.629 | 0.933 |
| Right Frontal Pole | 0.031 | 0.015 | 1.009 | [-0.034 - 0.095] | 0.000 | 1365 | 0.330 | 0.845 |
| Right Temporal Pole | 0.032 | 0.012 | 1.320 | [-0.020 - 0.084] | 0.000 | 1225 | 0.208 | 0.813 |
| Right Transverse Temporal | -0.002 | 0.009 | -0.117 | [-0.041 - 0.037] | 0.000 | 1363 | 0.908 | 0.944 |
| *Right Insula* | *-0.045* | *0.009* | *-2.511* | *[-0.084 - -0.007]* | *0.000* | *1363* | *0.025* | *0.565* |
| Left Thickness | -0.029 | 0.012 | -1.164 | [-0.083 - 0.025] | 0.000 | 1370 | 0.264 | 1.000 |
| Right Thickness | -0.033 | 0.013 | -1.231 | [-0.089 - 0.024] | 0.000 | 1370 | 0.239 | 1.000 |

| ROI | Partial *r* | SE | *t*-Score | 95% CI | *I2* | *N* | *p-value* | FDR |
|---|---|---|---|---|---|---|---|---|
| Left Surface Area | -0.046 | 0.023 | -0.994 | [-0.144 - 0.053] | 55.705 | 1370 | 0.337 | 1.000 |
| Right Surface Area | -0.043 | 0.022 | -0.963 | [-0.138 - 0.053] | 53.299 | 1370 | 0.352 | 1.000 |
| Mean Bankssts | -0.004 | 0.013 | -0.168 | [-0.058 - 0.050] | 0.000 | 1230 | 0.869 | 0.934 |
| Mean Caudal Anterior Cingulate | 0.031 | 0.012 | 1.288 | [-0.021 - 0.083] | 0.000 | 1354 | 0.219 | 0.620 |
| Mean Caudal Middle Frontal | -0.061 | 0.016 | -1.855 | [-0.130 - 0.010] | 10.602 | 1323 | 0.085 | 0.620 |
| Mean Cuneus | -0.004 | 0.017 | -0.108 | [-0.076 - 0.069] | 0.000 | 1243 | 0.915 | 0.943 |
| Mean Entorhinal | 0.063 | 0.020 | 1.592 | [-0.022 - 0.147] | 37.808 | 1137 | 0.134 | 0.620 |
| Mean Fusiform | -0.034 | 0.015 | -1.108 | [-0.099 - 0.032] | 2.753 | 1240 | 0.286 | 0.699 |
| Mean Inferior Parietal | -0.022 | 0.015 | -0.714 | [-0.088 - 0.044] | 0.000 | 1160 | 0.487 | 0.770 |
| Mean Inferior Temporal | -0.021 | 0.014 | -0.741 | [-0.081 - 0.040] | 0.000 | 1227 | 0.471 | 0.770 |
| Mean Isthmus Cingulate | 0.030 | 0.015 | 0.992 | [-0.034 - 0.094] | 0.000 | 1345 | 0.338 | 0.722 |
| Mean Lateral Occipital | -0.032 | 0.018 | -0.879 | [-0.110 - 0.046] | 0.000 | 1237 | 0.394 | 0.722 |
| Mean Lateral Orbitofrontal | -0.038 | 0.014 | -1.377 | [-0.098 - 0.021] | 0.000 | 1327 | 0.190 | 0.620 |
| Mean Lingual | -0.017 | 0.015 | -0.581 | [-0.081 - 0.047] | 0.000 | 1328 | 0.570 | 0.808 |
| Mean Medial Orbitofrontal | -0.015 | 0.015 | -0.480 | [-0.079 - 0.050] | 0.000 | 1298 | 0.639 | 0.835 |
| Mean Middle Temporal | -0.012 | 0.018 | -0.337 | [-0.090 - 0.065] | 12.345 | 1162 | 0.741 | 0.900 |
| Mean Parahippocampal | 0.040 | 0.016 | 1.294 | [-0.027 - 0.107] | 0.000 | 1305 | 0.217 | 0.620 |
| Mean Paracentral | -0.024 | 0.011 | -1.104 | [-0.071 - 0.023] | 0.000 | 1352 | 0.288 | 0.699 |
| Mean Pars Opercularis | -0.061 | 0.018 | -1.692 | [-0.137 - 0.016] | 0.000 | 1273 | 0.113 | 0.620 |
| *Mean Pars Orbitalis* | *-0.064* | *0.011* | *-2.888* | *[-0.111 - -0.017]* | *0.000* | *1301* | *0.012* | *0.204* |
| Mean Pars Triangularis | -0.047 | 0.015 | -1.597 | [-0.110 - 0.016] | 0.000 | 1256 | 0.133 | 0.620 |
| Mean Pericalcarine | 0.021 | 0.019 | 0.544 | [-0.061 - 0.102] | 0.000 | 1340 | 0.595 | 0.809 |
| Mean Postcentral | -0.013 | 0.017 | -0.377 | [-0.084 - 0.059] | 0.000 | 1316 | 0.712 | 0.897 |
| Mean Posterior Cingulate | 0.011 | 0.008 | 0.696 | [-0.024 - 0.046] | 0.000 | 1358 | 0.498 | 0.770 |
| Mean Precentral | -0.039 | 0.015 | -1.309 | [-0.104 - 0.025] | 0.000 | 1318 | 0.212 | 0.620 |
| Mean Precuneus | -0.004 | 0.011 | -0.197 | [-0.051 - 0.042] | 0.000 | 1311 | 0.846 | 0.934 |
| Mean Rostral Anterior Cingulate | -0.005 | 0.016 | -0.160 | [-0.072 - 0.062] | 0.000 | 1309 | 0.875 | 0.934 |
| *Mean Rostral Middle Frontal* | *-0.062* | *0.011* | *-2.885* | *[-0.109 - -0.016]* | *0.000* | *1263* | *0.012* | *0.204* |
| *Mean Superior Frontal* | *-0.068* | *0.014* | *-2.483* | *[-0.126 - -0.009]* | *0.000* | *1325* | *0.026* | *0.298* |
| Mean Superior Parietal | -0.005 | 0.016 | -0.155 | [-0.074 - 0.064] | 0.000 | 1248 | 0.879 | 0.934 |
| Mean Superior Temporal | 0.001 | 0.016 | 0.026 | [-0.067 - 0.069] | 0.000 | 1132 | 0.980 | 0.980 |
| Mean Supramarginal | -0.029 | 0.017 | -0.867 | [-0.102 - 0.043] | 0.000 | 1147 | 0.401 | 0.722 |
| Mean Frontal Pole | 0.027 | 0.016 | 0.862 | [-0.040 - 0.094] | 0.000 | 1363 | 0.403 | 0.722 |
| Mean Temporal Pole | 0.023 | 0.012 | 0.942 | [-0.030 - 0.077] | 0.000 | 1206 | 0.362 | 0.722 |
| Mean Transverse Temporal | 0.014 | 0.012 | 0.611 | [-0.036 - 0.065] | 0.000 | 1356 | 0.551 | 0.808 |
| Mean Insula | -0.026 | 0.010 | -1.316 | [-0.068 - 0.016] | 0.000 | 1358 | 0.209 | 0.620 |

Note: nominally significant results are italicized and FDR significant results are italicized and bolded

Supplementary Table S9 Associations between Cortical Thickness and SANS Avolition (Factor 3)

| ROI | Partial *r* | SE | *t*-Score | 95% CI | *I2* | *N* | *p-value* | FDR |
|---|---|---|---|---|---|---|---|---|
| Left Bankssts | -0.041 | 0.016 | -1.306 | [-0.107 - 0.026] | 0.000 | 1266 | 0.213 | 0.601 |
| Left Caudal Anterior Cingulate | 0.030 | 0.015 | 0.965 | [-0.036 - 0.095] | 0.000 | 1362 | 0.351 | 0.681 |
| Left Caudal Middle Frontal | -0.044 | 0.014 | -1.560 | [-0.104 - 0.016] | 0.000 | 1351 | 0.141 | 0.506 |
| Left Cuneus | 0.008 | 0.015 | 0.263 | [-0.057 - 0.074] | 0.000 | 1275 | 0.796 | 0.967 |
| Left Entorhinal | 0.001 | 0.015 | 0.029 | [-0.062 - 0.064] | 0.000 | 1278 | 0.977 | 0.991 |
| Left Fusiform | -0.048 | 0.015 | -1.596 | [-0.112 - 0.017] | 0.000 | 1308 | 0.133 | 0.506 |
| Left Inferior Parietal | 0.046 | 0.016 | 1.457 | [-0.022 - 0.113] | 0.000 | 1234 | 0.167 | 0.532 |
| Left Inferior Temporal | -0.028 | 0.019 | -0.725 | [-0.110 - 0.055] | 0.000 | 1275 | 0.480 | 0.726 |
| Left Isthmus Cingulate | 0.003 | 0.012 | 0.123 | [-0.050 - 0.056] | 0.000 | 1361 | 0.904 | 0.991 |

| ROI | Partial *r* | SE | *t*-Score | 95% CI | *I2* | *N* | *p-value* | FDR |
|---|---|---|---|---|---|---|---|---|
| Left Lateral Occipital | -0.002 | 0.020 | -0.052 | [-0.087 - 0.083] | 34.792 | 1296 | 0.960 | 0.991 |
| *Left Lateral Orbitofrontal* | *-0.044* | *0.010* | *-2.165* | *[-0.087 - 0.000]* | *0.000* | *1364* | *0.048* | *0.374* |
| Left Lingual | -0.026 | 0.016 | -0.795 | [-0.096 - 0.044] | 0.000 | 1339 | 0.440 | 0.712 |
| Left Medial Orbitofrontal | -0.013 | 0.014 | -0.489 | [-0.071 - 0.045] | 0.000 | 1347 | 0.633 | 0.896 |
| Left Middle Temporal | -0.018 | 0.014 | -0.643 | [-0.077 - 0.042] | 0.000 | 1218 | 0.530 | 0.784 |
| Left Parahippocampal | 0.005 | 0.018 | 0.129 | [-0.074 - 0.083] | 31.099 | 1342 | 0.899 | 0.991 |
| Left Paracentral | -0.054 | 0.014 | -1.943 | [-0.113 - 0.006] | 0.000 | 1360 | 0.072 | 0.409 |
| *Left Pars Opercularis* | *-0.077* | *0.017* | *-2.213* | *[-0.150 - -0.002]* | *0.000* | *1323* | *0.044* | *0.374* |
| *Left Pars Orbitalis* | *-0.045* | *0.010* | *-2.285* | *[-0.088 - -0.003]* | *0.000* | *1327* | *0.038* | *0.374* |
| ***Left Pars Triangularis*** | ***-0.086*** | ***0.010*** | ***-4.475*** | ***[-0.127 - -0.045]*** | ***0.000*** | ***1304*** | ***0.001*** | ***0.027*** |
| Left Pericalcarine | 0.002 | 0.020 | 0.053 | [-0.083 - 0.088] | 44.373 | 1350 | 0.958 | 0.991 |
| Left Postcentral | -0.049 | 0.017 | -1.440 | [-0.123 - 0.024] | 0.000 | 1336 | 0.172 | 0.532 |
| Left Posterior Cingulate | -0.014 | 0.019 | -0.359 | [-0.097 - 0.069] | 30.096 | 1363 | 0.725 | 0.954 |
| Left Precentral | -0.063 | 0.016 | -1.977 | [-0.131 - 0.005] | 0.000 | 1344 | 0.068 | 0.409 |
| Left Precuneus | -0.018 | 0.011 | -0.796 | [-0.065 - 0.030] | 0.000 | 1337 | 0.440 | 0.712 |
| Left Rostral Anterior Cingulate | 0.004 | 0.013 | 0.145 | [-0.050 - 0.057] | 0.000 | 1347 | 0.887 | 0.991 |
| Left Rostral Middle Frontal | -0.042 | 0.011 | -1.855 | [-0.091 - 0.007] | 0.000 | 1312 | 0.085 | 0.412 |
| Left Superior Frontal | -0.060 | 0.014 | -2.093 | [-0.120 - 0.001] | 9.684 | 1350 | 0.055 | 0.374 |
| Left Superior Parietal | 0.010 | 0.016 | 0.292 | [-0.061 - 0.080] | 0.000 | 1283 | 0.774 | 0.960 |
| Left Superior Temporal | -0.041 | 0.016 | -1.276 | [-0.110 - 0.028] | 0.000 | 1197 | 0.223 | 0.601 |
| Left Supramarginal | -0.032 | 0.016 | -0.986 | [-0.100 - 0.037] | 0.000 | 1216 | 0.341 | 0.681 |
| Left Frontal Pole | -0.037 | 0.012 | -1.573 | [-0.087 - 0.013] | 0.000 | 1368 | 0.138 | 0.506 |
| *Left Temporal Pole* | *-0.066* | *0.011* | *-3.098* | *[-0.111 - -0.020]* | *0.000* | *1329* | *0.008* | *0.178* |
| Left Transverse Temporal | 0.007 | 0.016 | 0.209 | [-0.063 - 0.077] | 0.000 | 1362 | 0.838 | 0.982 |
| *Left Insula* | *-0.087* | *0.017* | *-2.518* | *[-0.160 - -0.013]* | *24.260* | *1365* | *0.025* | *0.334* |
| Right Bankssts | 0.000 | 0.017 | 0.012 | [-0.070 - 0.071] | 0.000 | 1317 | 0.991 | 0.991 |
| Right Caudal Anterior Cingulate | 0.028 | 0.013 | 1.047 | [-0.029 - 0.085] | 0.000 | 1361 | 0.313 | 0.681 |
| ***Right Caudal Middle Frontal*** | ***-0.078*** | ***0.009*** | ***-4.258*** | ***[-0.116 - -0.039]*** | ***0.000*** | ***1337*** | ***0.001*** | ***0.027*** |
| Right Cuneus | 0.021 | 0.012 | 0.928 | [-0.028 - 0.071] | 0.000 | 1316 | 0.369 | 0.681 |
| Right Entorhinal | -0.007 | 0.012 | -0.290 | [-0.057 - 0.043] | 0.000 | 1164 | 0.776 | 0.960 |
| Right Fusiform | -0.038 | 0.016 | -1.207 | [-0.104 - 0.029] | 0.000 | 1281 | 0.247 | 0.601 |
| Right Inferior Parietal | -0.004 | 0.020 | -0.108 | [-0.090 - 0.081] | 0.000 | 1226 | 0.915 | 0.991 |
| Right Inferior Temporal | -0.032 | 0.017 | -0.932 | [-0.107 - 0.042] | 12.277 | 1283 | 0.367 | 0.681 |
| Right Isthmus Cingulate | 0.010 | 0.012 | 0.393 | [-0.043 - 0.063] | 0.000 | 1351 | 0.700 | 0.954 |
| Right Lateral Occipital | -0.015 | 0.021 | -0.353 | [-0.106 - 0.076] | 41.221 | 1287 | 0.729 | 0.954 |
| Right Lateral Orbitofrontal | -0.038 | 0.012 | -1.559 | [-0.090 - 0.014] | 0.000 | 1333 | 0.141 | 0.506 |
| Right Lingual | -0.023 | 0.015 | -0.773 | [-0.085 - 0.040] | 0.000 | 1350 | 0.452 | 0.715 |
| Right Medial Orbitofrontal | -0.032 | 0.016 | -1.002 | [-0.100 - 0.036] | 0.000 | 1316 | 0.333 | 0.681 |
| Right Middle Temporal | -0.037 | 0.021 | -0.893 | [-0.125 - 0.052] | 39.163 | 1255 | 0.387 | 0.681 |
| Right Parahippocampal | 0.005 | 0.013 | 0.210 | [-0.050 - 0.061] | 0.000 | 1322 | 0.837 | 0.982 |
| Right Paracentral | -0.026 | 0.015 | -0.886 | [-0.090 - 0.037] | 0.000 | 1361 | 0.390 | 0.681 |
| Right Pars Opercularis | -0.048 | 0.018 | -1.352 | [-0.124 - 0.028] | 9.199 | 1305 | 0.198 | 0.584 |
| Right Pars Orbitalis | -0.033 | 0.013 | -1.228 | [-0.090 - 0.024] | 0.000 | 1336 | 0.240 | 0.601 |
| Right Pars Triangularis | -0.020 | 0.013 | -0.731 | [-0.077 - 0.038] | 0.000 | 1305 | 0.477 | 0.726 |
| Right Pericalcarine | -0.031 | 0.015 | -0.995 | [-0.097 - 0.036] | 0.000 | 1350 | 0.337 | 0.681 |
| Right Postcentral | -0.046 | 0.019 | -1.214 | [-0.127 - 0.035] | 43.110 | 1342 | 0.245 | 0.601 |
| Right Posterior Cingulate | -0.011 | 0.015 | -0.368 | [-0.074 - 0.052] | 0.000 | 1364 | 0.718 | 0.954 |
| Right Precentral | -0.034 | 0.010 | -1.636 | [-0.078 - 0.010] | 0.000 | 1333 | 0.124 | 0.506 |
| Right Precuneus | 0.013 | 0.012 | 0.552 | [-0.039 - 0.066] | 0.000 | 1333 | 0.590 | 0.853 |
| Right Rostral Anterior Cingulate | -0.022 | 0.013 | -0.860 | [-0.078 - 0.033] | 0.000 | 1327 | 0.405 | 0.688 |
| Right Rostral Middle Frontal | -0.043 | 0.011 | -1.901 | [-0.092 - 0.006] | 0.000 | 1302 | 0.078 | 0.409 |
| *Right Superior Frontal* | *-0.069* | *0.012* | *-2.953* | *[-0.118 - -0.019]* | *0.000* | *1341* | *0.010* | *0.178* |

| ROI | Partial *r* | SE | *t*-Score | 95% CI | *I2* | *N* | *p-value* | FDR |
|---|---|---|---|---|---|---|---|---|
| Right Superior Parietal | 0.009 | 0.014 | 0.333 | [-0.051 - 0.069] | 0.000 | 1314 | 0.744 | 0.955 |
| Right Superior Temporal | -0.025 | 0.013 | -0.923 | [-0.082 - 0.033] | 0.000 | 1234 | 0.372 | 0.681 |
| Right Supramarginal | -0.064 | 0.022 | -1.476 | [-0.157 - 0.029] | 43.570 | 1223 | 0.162 | 0.532 |
| Right Frontal Pole | 0.001 | 0.012 | 0.047 | [-0.051 - 0.054] | 0.000 | 1365 | 0.963 | 0.991 |
| Right Temporal Pole | 0.000 | 0.013 | -0.016 | [-0.056 - 0.055] | 0.000 | 1225 | 0.987 | 0.991 |
| Right Transverse Temporal | -0.032 | 0.014 | -1.157 | [-0.090 - 0.027] | 0.000 | 1363 | 0.267 | 0.626 |
| Right Insula | -0.074 | 0.017 | -2.142 | [-0.148 - 0.000] | 0.000 | 1363 | 0.050 | 0.374 |
| Left Thickness | -0.042 | 0.015 | -1.434 | [-0.104 - 0.021] | 0.000 | 1370 | 0.174 | 1.000 |
| Right Thickness | -0.040 | 0.015 | -1.316 | [-0.104 - 0.025] | 0.000 | 1370 | 0.209 | 1.000 |
| *Left Surface Area* | *-0.090* | *0.013* | *-3.320* | *[-0.147 - -0.032]* | *0.000* | *1370* | *0.005* | *1.000* |
| *Right Surface Area* | *-0.090* | *0.012* | *-3.725* | *[-0.141 - -0.038]* | *0.000* | *1370* | *0.002* | *1.000* |
| Mean Bankssts | -0.026 | 0.016 | -0.794 | [-0.097 - 0.044] | 0.000 | 1230 | 0.440 | 0.748 |
| Mean Caudal Anterior Cingulate | 0.038 | 0.017 | 1.141 | [-0.033 - 0.109] | 0.000 | 1354 | 0.273 | 0.580 |
| *Mean Caudal Middle Frontal* | *-0.066* | *0.011* | *-2.897* | *[-0.115 - -0.017]* | *0.000* | *1323* | *0.012* | *0.199* |
| Mean Cuneus | 0.020 | 0.014 | 0.712 | [-0.041 - 0.081] | 0.000 | 1243 | 0.488 | 0.759 |
| Mean Entorhinal | 0.001 | 0.014 | 0.029 | [-0.059 - 0.060] | 0.000 | 1137 | 0.977 | 0.977 |
| Mean Fusiform | -0.044 | 0.017 | -1.280 | [-0.116 - 0.029] | 11.202 | 1240 | 0.221 | 0.537 |
| Mean Inferior Parietal | 0.021 | 0.018 | 0.566 | [-0.058 - 0.099] | 0.000 | 1160 | 0.580 | 0.822 |
| Mean Inferior Temporal | -0.042 | 0.023 | -0.892 | [-0.142 - 0.059] | 60.693 | 1227 | 0.387 | 0.693 |
| Mean Isthmus Cingulate | 0.006 | 0.013 | 0.245 | [-0.049 - 0.062] | 0.000 | 1345 | 0.810 | 0.927 |
| Mean Lateral Occipital | -0.005 | 0.022 | -0.110 | [-0.101 - 0.091] | 44.717 | 1237 | 0.914 | 0.971 |
| Mean Lateral Orbitofrontal | -0.043 | 0.011 | -1.970 | [-0.090 - 0.004] | 0.000 | 1327 | 0.069 | 0.327 |
| Mean Lingual | -0.032 | 0.016 | -0.989 | [-0.101 - 0.038] | 0.000 | 1328 | 0.339 | 0.641 |
| Mean Medial Orbitofrontal | -0.022 | 0.016 | -0.707 | [-0.089 - 0.045] | 0.000 | 1298 | 0.491 | 0.759 |
| Mean Middle Temporal | -0.026 | 0.019 | -0.656 | [-0.109 - 0.058] | 24.911 | 1162 | 0.523 | 0.772 |
| Mean Parahippocampal | 0.008 | 0.016 | 0.235 | [-0.063 - 0.078] | 3.696 | 1305 | 0.818 | 0.927 |
| Mean Paracentral | -0.038 | 0.015 | -1.292 | [-0.100 - 0.025] | 0.000 | 1352 | 0.217 | 0.537 |
| Mean Pars Opercularis | -0.073 | 0.020 | -1.844 | [-0.157 - 0.012] | 21.844 | 1273 | 0.086 | 0.327 |
| *Mean Pars Orbitalis* | *-0.056* | *0.009* | *-3.233* | *[-0.093 - -0.019]* | *0.000* | *1301* | *0.006* | *0.199* |
| *Mean Pars Triangularis* | *-0.058* | *0.013* | *-2.315* | *[-0.112 - -0.004]* | *0.000* | *1256* | *0.036* | *0.213* |
| Mean Pericalcarine | -0.012 | 0.017 | -0.346 | [-0.086 - 0.062] | 0.000 | 1340 | 0.735 | 0.892 |
| Mean Postcentral | -0.060 | 0.017 | -1.755 | [-0.132 - 0.013] | 0.000 | 1316 | 0.101 | 0.344 |
| Mean Posterior Cingulate | -0.014 | 0.017 | -0.405 | [-0.088 - 0.060] | 18.157 | 1358 | 0.692 | 0.892 |
| *Mean Precentral* | *-0.060* | *0.013* | *-2.297* | *[-0.116 - -0.004]* | *0.000* | *1318* | *0.038* | *0.213* |
| Mean Precuneus | -0.001 | 0.012 | -0.059 | [-0.051 - 0.049] | 0.000 | 1311 | 0.954 | 0.977 |
| Mean Rostral Anterior Cingulate | -0.005 | 0.013 | -0.175 | [-0.061 - 0.052] | 0.000 | 1309 | 0.864 | 0.948 |
| Mean Rostral Middle Frontal | -0.044 | 0.012 | -1.886 | [-0.095 - 0.006] | 0.000 | 1263 | 0.080 | 0.327 |
| *Mean Superior Frontal* | *-0.066* | *0.012* | *-2.690* | *[-0.119 - -0.014]* | *0.000* | *1325* | *0.018* | *0.199* |
| Mean Superior Parietal | 0.012 | 0.016 | 0.375 | [-0.056 - 0.080] | 0.000 | 1248 | 0.713 | 0.892 |
| Mean Superior Temporal | -0.036 | 0.015 | -1.233 | [-0.099 - 0.027] | 0.000 | 1132 | 0.238 | 0.539 |
| Mean Supramarginal | -0.053 | 0.018 | -1.468 | [-0.131 - 0.025] | 0.000 | 1147 | 0.164 | 0.507 |
| Mean Frontal Pole | -0.020 | 0.010 | -1.001 | [-0.062 - 0.023] | 0.000 | 1363 | 0.334 | 0.641 |
| Mean Temporal Pole | -0.030 | 0.012 | -1.288 | [-0.081 - 0.020] | 0.000 | 1206 | 0.219 | 0.537 |
| Mean Transverse Temporal | -0.015 | 0.015 | -0.487 | [-0.079 - 0.050] | 0.000 | 1356 | 0.634 | 0.862 |
| *Mean Insula* | *-0.081* | *0.017* | *-2.322* | *[-0.154 - -0.006]* | *0.000* | *1358* | *0.036* | *0.213* |

Note: nominally significant results are italicized and FDR significant results are italicized and bolded

Supplementary Table S10 Associations between Cortical Thickness and SANS EXP

| ROI | Partial *r* | SE | *t*-Score | 95% CI | *I2* | *N* | *p-value* | FDR |
|---|---|---|---|---|---|---|---|---|
| Left Bankssts | -0.046 | 0.014 | -1.594 | [-0.108 - 0.016] | 0.000 | 1266 | 0.133 | 0.414 |
| Left Caudal Anterior Cingulate | 0.050 | 0.013 | 1.966 | [-0.005 - 0.104] | 0.000 | 1362 | 0.069 | 0.337 |
| Left Caudal Middle Frontal | -0.005 | 0.014 | -0.184 | [-0.066 - 0.056] | 0.000 | 1351 | 0.857 | 0.893 |
| Left Cuneus | -0.004 | 0.018 | -0.125 | [-0.080 - 0.071] | 0.000 | 1275 | 0.902 | 0.916 |
| Left Entorhinal | 0.007 | 0.009 | 0.423 | [-0.030 - 0.044] | 0.000 | 1278 | 0.679 | 0.839 |
| Left Fusiform | -0.058 | 0.014 | -2.073 | [-0.118 - 0.002] | 0.000 | 1308 | 0.057 | 0.298 |
| Left Inferior Parietal | -0.011 | 0.016 | -0.353 | [-0.080 - 0.057] | 0.000 | 1234 | 0.729 | 0.867 |
| Left Inferior Temporal | -0.007 | 0.012 | -0.317 | [-0.057 - 0.042] | 0.000 | 1275 | 0.756 | 0.867 |
| Left Isthmus Cingulate | -0.029 | 0.013 | -1.128 | [-0.084 - 0.026] | 0.000 | 1361 | 0.278 | 0.601 |
| Left Lateral Occipital | -0.043 | 0.015 | -1.467 | [-0.106 - 0.020] | 0.000 | 1296 | 0.165 | 0.426 |
| Left Lateral Orbitofrontal | -0.032 | 0.014 | -1.118 | [-0.093 - 0.029] | 0.000 | 1364 | 0.282 | 0.601 |
| Left Lingual | -0.047 | 0.016 | -1.449 | [-0.115 - 0.022] | 0.000 | 1339 | 0.169 | 0.426 |
| Left Medial Orbitofrontal | -0.051 | 0.015 | -1.722 | [-0.114 - 0.013] | 0.000 | 1347 | 0.107 | 0.405 |
| Left Middle Temporal | -0.026 | 0.012 | -1.095 | [-0.076 - 0.025] | 0.000 | 1218 | 0.292 | 0.601 |
| Left Parahippocampal | -0.023 | 0.015 | -0.772 | [-0.088 - 0.042] | 0.000 | 1342 | 0.453 | 0.671 |
| Left Paracentral | -0.009 | 0.016 | -0.301 | [-0.077 - 0.058] | 0.000 | 1360 | 0.768 | 0.867 |
| *Left Pars Opercularis* | *-0.069* | *0.013* | *-2.746* | *[-0.123 - -0.015]* | *0.000* | *1323* | *0.016* | *0.268* |
| *Left Pars Orbitalis* | *-0.074* | *0.009* | *-4.115* | *[-0.113 - -0.036]* | *0.000* | *1327* | *0.001* | *0.071* |
| *Left Pars Triangularis* | *-0.055* | *0.013* | *-2.181* | *[-0.109 - -0.001]* | *0.000* | *1304* | *0.047* | *0.298* |
| Left Pericalcarine | 0.006 | 0.016 | 0.181 | [-0.061 - 0.073] | 0.000 | 1350 | 0.859 | 0.893 |
| Left Postcentral | -0.040 | 0.014 | -1.465 | [-0.098 - 0.018] | 0.000 | 1336 | 0.165 | 0.426 |
| Left Posterior Cingulate | -0.058 | 0.015 | -1.917 | [-0.121 - 0.007] | 0.000 | 1363 | 0.076 | 0.344 |
| Left Precentral | -0.055 | 0.015 | -1.779 | [-0.120 - 0.011] | 0.000 | 1344 | 0.097 | 0.389 |
| Left Precuneus | -0.032 | 0.015 | -1.054 | [-0.097 - 0.033] | 0.000 | 1337 | 0.310 | 0.601 |
| Left Rostral Anterior Cingulate | 0.046 | 0.013 | 1.778 | [-0.009 - 0.101] | 0.000 | 1347 | 0.097 | 0.389 |
| *Left Rostral Middle Frontal* | *-0.058* | *0.013* | *-2.298* | *[-0.111 - -0.004]* | *0.000* | *1312* | *0.038* | *0.298* |
| *Left Superior Frontal* | *-0.081* | *0.011* | *-3.530* | *[-0.130 - -0.032]* | *0.000* | *1350* | *0.003* | *0.075* |
| Left Superior Parietal | -0.014 | 0.021 | -0.327 | [-0.106 - 0.078] | 49.127 | 1283 | 0.748 | 0.867 |
| *Left Superior Temporal* | *-0.060* | *0.012* | *-2.492* | *[-0.111 - -0.008]* | *0.000* | *1197* | *0.026* | *0.298* |
| *Left Supramarginal* | *-0.062* | *0.014* | *-2.249* | *[-0.122 - -0.003]* | *0.000* | *1216* | *0.041* | *0.298* |
| Left Frontal Pole | -0.014 | 0.013 | -0.551 | [-0.068 - 0.040] | 0.000 | 1368 | 0.590 | 0.787 |
| Left Temporal Pole | 0.008 | 0.009 | 0.449 | [-0.031 - 0.047] | 0.000 | 1329 | 0.661 | 0.832 |
| Left Transverse Temporal | -0.008 | 0.012 | -0.339 | [-0.061 - 0.044] | 0.000 | 1362 | 0.740 | 0.867 |
| Left Insula | -0.026 | 0.016 | -0.820 | [-0.092 - 0.041] | 0.000 | 1365 | 0.426 | 0.671 |
| Right Bankssts | -0.059 | 0.026 | -1.118 | [-0.171 - 0.054] | 70.770 | 1317 | 0.282 | 0.601 |
| Right Caudal Anterior Cingulate | -0.003 | 0.020 | -0.067 | [-0.087 - 0.082] | 0.000 | 1361 | 0.948 | 0.948 |
| *Right Caudal Middle Frontal* | *-0.066* | *0.015* | *-2.180* | *[-0.131 - -0.001]* | *0.000* | *1337* | *0.047* | *0.298* |
| Right Cuneus | -0.053 | 0.018 | -1.477 | [-0.130 - 0.024] | 15.034 | 1316 | 0.162 | 0.426 |
| Right Entorhinal | 0.034 | 0.016 | 1.067 | [-0.034 - 0.101] | 0.000 | 1164 | 0.304 | 0.601 |
| Right Fusiform | -0.016 | 0.013 | -0.578 | [-0.073 - 0.042] | 0.000 | 1281 | 0.572 | 0.783 |
| Right Inferior Parietal | -0.033 | 0.020 | -0.827 | [-0.118 - 0.053] | 0.000 | 1226 | 0.422 | 0.671 |
| Right Inferior Temporal | -0.021 | 0.013 | -0.807 | [-0.077 - 0.035] | 0.000 | 1283 | 0.433 | 0.671 |
| Right Isthmus Cingulate | -0.012 | 0.013 | -0.454 | [-0.069 - 0.045] | 0.000 | 1351 | 0.657 | 0.832 |
| Right Lateral Occipital | -0.024 | 0.015 | -0.788 | [-0.091 - 0.042] | 0.000 | 1287 | 0.444 | 0.671 |
| Right Lateral Orbitofrontal | -0.048 | 0.016 | -1.518 | [-0.114 - 0.020] | 0.000 | 1333 | 0.151 | 0.426 |
| Right Lingual | -0.036 | 0.013 | -1.397 | [-0.091 - 0.019] | 0.000 | 1350 | 0.184 | 0.447 |
| Right Medial Orbitofrontal | -0.048 | 0.014 | -1.642 | [-0.109 - 0.015] | 0.000 | 1316 | 0.123 | 0.414 |
| Right Middle Temporal | -0.006 | 0.018 | -0.185 | [-0.082 - 0.069] | 0.000 | 1255 | 0.856 | 0.893 |
| Right Parahippocampal | 0.011 | 0.010 | 0.575 | [-0.031 - 0.054] | 0.000 | 1322 | 0.574 | 0.783 |
| Right Paracentral | -0.030 | 0.016 | -0.969 | [-0.098 - 0.037] | 0.000 | 1361 | 0.349 | 0.642 |
| Right Pars Opercularis | -0.055 | 0.013 | -2.113 | [-0.110 - 0.001] | 0.000 | 1305 | 0.053 | 0.298 |
| Right Pars Orbitalis | -0.015 | 0.013 | -0.573 | [-0.072 - 0.042] | 0.000 | 1336 | 0.576 | 0.783 |

| ROI | Partial *r* | SE | *t*-Score | 95% CI | *I2* | *N* | *p-value* | FDR |
|---|---|---|---|---|---|---|---|---|
| *Right Pars Triangularis* | *-0.058* | *0.012* | *-2.457* | *[-0.108 - -0.007]* | *0.000* | *1305* | *0.028* | *0.298* |
| Right Pericalcarine | -0.024 | 0.015 | -0.795 | [-0.087 - 0.040] | 0.000 | 1350 | 0.440 | 0.671 |
| Right Postcentral | -0.020 | 0.012 | -0.855 | [-0.070 - 0.030] | 0.000 | 1342 | 0.407 | 0.671 |
| Right Posterior Cingulate | -0.015 | 0.017 | -0.456 | [-0.086 - 0.056] | 0.000 | 1364 | 0.655 | 0.832 |
| Right Precentral | -0.038 | 0.012 | -1.591 | [-0.088 - 0.013] | 0.000 | 1333 | 0.134 | 0.414 |
| Right Precuneus | -0.007 | 0.018 | -0.195 | [-0.082 - 0.068] | 0.000 | 1333 | 0.848 | 0.893 |
| Right Rostral Anterior Cingulate | 0.006 | 0.017 | 0.171 | [-0.066 - 0.078] | 0.000 | 1327 | 0.867 | 0.893 |
| Right Rostral Middle Frontal | -0.057 | 0.014 | -2.077 | [-0.115 - 0.002] | 0.000 | 1302 | 0.057 | 0.298 |
| *Right Superior Frontal* | *-0.081* | *0.011* | *-3.593* | *[-0.129 - -0.033]* | *0.000* | *1341* | *0.003* | *0.075* |
| Right Superior Parietal | -0.010 | 0.018 | -0.287 | [-0.086 - 0.065] | 22.714 | 1314 | 0.778 | 0.867 |
| Right Superior Temporal | -0.040 | 0.020 | -1.014 | [-0.124 - 0.045] | 0.000 | 1234 | 0.328 | 0.619 |
| Right Supramarginal | -0.051 | 0.016 | -1.645 | [-0.117 - 0.016] | 0.000 | 1223 | 0.122 | 0.414 |
| Right Frontal Pole | -0.019 | 0.010 | -0.916 | [-0.063 - 0.025] | 0.000 | 1365 | 0.375 | 0.671 |
| Right Temporal Pole | -0.009 | 0.006 | -0.732 | [-0.037 - 0.018] | 0.000 | 1225 | 0.476 | 0.689 |
| Right Transverse Temporal | -0.018 | 0.011 | -0.770 | [-0.067 - 0.032] | 0.000 | 1363 | 0.454 | 0.671 |
| Right Insula | -0.034 | 0.016 | -1.074 | [-0.100 - 0.033] | 0.000 | 1363 | 0.301 | 0.601 |
| Left Thickness | -0.049 | 0.013 | -1.847 | [-0.105 - 0.008] | 0.000 | 1370 | 0.086 | 1.000 |
| Right Thickness | -0.052 | 0.013 | -2.044 | [-0.106 - 0.003] | 0.000 | 1370 | 0.060 | 1.000 |
| Left Surface Area | -0.022 | 0.014 | -0.754 | [-0.084 - 0.040] | 0.000 | 1370 | 0.463 | 1.000 |
| Right Surface Area | -0.014 | 0.015 | -0.454 | [-0.078 - 0.051] | 0.000 | 1370 | 0.657 | 1.000 |
| Mean Bankssts | -0.071 | 0.024 | -1.475 | [-0.173 - 0.032] | 63.345 | 1230 | 0.162 | 0.460 |
| Mean Caudal Anterior Cingulate | 0.035 | 0.016 | 1.075 | [-0.035 - 0.104] | 0.000 | 1354 | 0.300 | 0.565 |
| Mean Caudal Middle Frontal | -0.036 | 0.015 | -1.199 | [-0.101 - 0.029] | 0.000 | 1323 | 0.250 | 0.516 |
| Mean Cuneus | -0.042 | 0.021 | -1.004 | [-0.132 - 0.048] | 47.918 | 1243 | 0.332 | 0.565 |
| Mean Entorhinal | 0.015 | 0.011 | 0.719 | [-0.030 - 0.061] | 0.000 | 1137 | 0.484 | 0.629 |
| Mean Fusiform | -0.046 | 0.013 | -1.802 | [-0.101 - 0.009] | 0.000 | 1240 | 0.093 | 0.316 |
| Mean Inferior Parietal | -0.033 | 0.018 | -0.888 | [-0.111 - 0.046] | 0.000 | 1160 | 0.390 | 0.576 |
| Mean Inferior Temporal | -0.009 | 0.012 | -0.409 | [-0.059 - 0.040] | 0.000 | 1227 | 0.689 | 0.744 |
| Mean Isthmus Cingulate | -0.023 | 0.013 | -0.895 | [-0.078 - 0.032] | 0.000 | 1345 | 0.386 | 0.576 |
| Mean Lateral Occipital | -0.039 | 0.015 | -1.272 | [-0.104 - 0.027] | 0.000 | 1237 | 0.224 | 0.508 |
| Mean Lateral Orbitofrontal | -0.039 | 0.015 | -1.310 | [-0.102 - 0.025] | 0.000 | 1327 | 0.211 | 0.508 |
| Mean Lingual | -0.051 | 0.015 | -1.748 | [-0.113 - 0.012] | 0.000 | 1328 | 0.102 | 0.316 |
| Mean Medial Orbitofrontal | -0.056 | 0.015 | -1.885 | [-0.120 - 0.008] | 0.000 | 1298 | 0.080 | 0.304 |
| Mean Middle Temporal | -0.023 | 0.016 | -0.742 | [-0.090 - 0.044] | 0.000 | 1162 | 0.470 | 0.629 |
| Mean Parahippocampal | -0.007 | 0.013 | -0.278 | [-0.064 - 0.049] | 0.000 | 1305 | 0.785 | 0.785 |
| Mean Paracentral | -0.021 | 0.017 | -0.641 | [-0.092 - 0.050] | 0.000 | 1352 | 0.532 | 0.646 |
| *Mean Pars Opercularis* | *-0.072* | *0.013* | *-2.756* | *[-0.128 - -0.016]* | *0.000* | *1273* | *0.015* | *0.131* |
| *Mean Pars Orbitalis* | *-0.062* | *0.011* | *-2.911* | *[-0.108 - -0.016]* | *0.000* | *1301* | *0.011* | *0.129* |
| *Mean Pars Triangularis* | *-0.075* | *0.012* | *-2.974* | *[-0.128 - -0.021]* | *0.000* | *1256* | *0.010* | *0.129* |
| Mean Pericalcarine | -0.011 | 0.015 | -0.373 | [-0.073 - 0.052] | 0.000 | 1340 | 0.715 | 0.744 |
| Mean Postcentral | -0.034 | 0.012 | -1.397 | [-0.087 - 0.018] | 0.000 | 1316 | 0.184 | 0.482 |
| Mean Posterior Cingulate | -0.038 | 0.016 | -1.180 | [-0.108 - 0.031] | 0.000 | 1358 | 0.258 | 0.516 |
| *Mean Precentral* | *-0.059* | *0.013* | *-2.275* | *[-0.115 - -0.003]* | *0.000* | *1318* | *0.039* | *0.190* |
| Mean Precuneus | -0.020 | 0.018 | -0.556 | [-0.096 - 0.056] | 0.000 | 1311 | 0.587 | 0.688 |
| Mean Rostral Anterior Cingulate | 0.030 | 0.016 | 0.946 | [-0.038 - 0.097] | 0.000 | 1309 | 0.360 | 0.576 |
| *Mean Rostral Middle Frontal* | *-0.067* | *0.013* | *-2.498* | *[-0.123 - -0.009]* | *0.000* | *1263* | *0.026* | *0.166* |
| *Mean Superior Frontal* | *-0.083* | *0.012* | *-3.590* | *[-0.132 - -0.033]* | *0.000* | *1325* | *0.003* | *0.100* |
| Mean Superior Parietal | -0.019 | 0.021 | -0.446 | [-0.110 - 0.073] | 46.494 | 1248 | 0.663 | 0.744 |
| Mean Superior Temporal | -0.059 | 0.015 | -1.973 | [-0.122 - 0.005] | 0.000 | 1132 | 0.069 | 0.291 |
| *Mean Supramarginal* | *-0.070* | *0.014* | *-2.427* | *[-0.131 - -0.008]* | *0.000* | *1147* | *0.029* | *0.166* |
| Mean Frontal Pole | -0.015 | 0.010 | -0.694 | [-0.059 - 0.030] | 0.000 | 1363 | 0.499 | 0.629 |

| ROI | Partial *r* | SE | *t*-Score | 95% CI | *I*2 | *N* | *p-value* | FDR |
|---|---|---|---|---|---|---|---|---|
| Mean Temporal Pole | 0.004 | 0.006 | 0.363 | [-0.022 - 0.031] | 0.000 | 1206 | 0.722 | 0.744 |
| Mean Transverse Temporal | -0.016 | 0.011 | -0.740 | [-0.062 - 0.030] | 0.000 | 1356 | 0.471 | 0.629 |
| Mean Insula | -0.032 | 0.016 | -1.024 | [-0.099 - 0.035] | 0.000 | 1358 | 0.323 | 0.565 |

Note: nominally significant results are italicized and FDR significant results are italicized and bolded

Supplementary Table S11 Associations between Cortical Thickness and SANS Blunted Affect (Factor 4)

| ROI | Partial *r* | SE | *t*-Score | 95% CI | *I*2 | *N* | *p-value* | FDR |
|---|---|---|---|---|---|---|---|---|
| Left Bankssts | -0.041 | 0.014 | -1.447 | [-0.102 - 0.020] | 0.000 | 1266 | 0.170 | 0.578 |
| *Left Caudal Anterior Cingulate* | *0.051* | *0.011* | *2.296* | *[ 0.003 - 0.098]* | *0.000* | *1362* | *0.038* | *0.458* |
| Left Caudal Middle Frontal | 0.011 | 0.014 | 0.414 | [-0.047 - 0.069] | 0.000 | 1351 | 0.685 | 0.867 |
| Left Cuneus | -0.009 | 0.020 | -0.231 | [-0.096 - 0.077] | 0.000 | 1275 | 0.820 | 0.950 |
| Left Entorhinal | 0.009 | 0.008 | 0.584 | [-0.024 - 0.041] | 0.000 | 1278 | 0.568 | 0.867 |
| Left Fusiform | -0.045 | 0.014 | -1.668 | [-0.104 - 0.013] | 0.000 | 1308 | 0.118 | 0.458 |
| Left Inferior Parietal | -0.001 | 0.017 | -0.023 | [-0.072 - 0.070] | 0.000 | 1234 | 0.982 | 0.982 |
| Left Inferior Temporal | 0.001 | 0.012 | 0.025 | [-0.052 - 0.053] | 0.000 | 1275 | 0.980 | 0.982 |
| Left Isthmus Cingulate | -0.028 | 0.014 | -1.037 | [-0.087 - 0.030] | 0.000 | 1361 | 0.317 | 0.775 |
| Left Lateral Occipital | -0.037 | 0.016 | -1.131 | [-0.107 - 0.033] | 0.000 | 1296 | 0.277 | 0.754 |
| Left Lateral Orbitofrontal | -0.028 | 0.014 | -0.971 | [-0.090 - 0.034] | 0.000 | 1364 | 0.348 | 0.778 |
| Left Lingual | -0.038 | 0.018 | -1.051 | [-0.115 - 0.040] | 0.000 | 1339 | 0.311 | 0.775 |
| Left Medial Orbitofrontal | -0.054 | 0.015 | -1.832 | [-0.117 - 0.009] | 0.000 | 1347 | 0.088 | 0.458 |
| Left Middle Temporal | -0.013 | 0.011 | -0.551 | [-0.062 - 0.036] | 0.000 | 1218 | 0.591 | 0.867 |
| Left Parahippocampal | -0.015 | 0.016 | -0.450 | [-0.085 - 0.056] | 13.620 | 1342 | 0.660 | 0.867 |
| Left Paracentral | 0.006 | 0.015 | 0.202 | [-0.058 - 0.070] | 0.000 | 1360 | 0.843 | 0.955 |
| *Left Pars Opercularis* | *-0.060* | *0.013* | *-2.265* | *[-0.117 - -0.003]* | *0.000* | *1323* | *0.040* | *0.458* |
| *Left Pars Orbitalis* | *-0.081* | *0.010* | *-3.987* | *[-0.125 - -0.038]* | *0.000* | *1327* | *0.001* | *0.092* |
| Left Pars Triangularis | -0.044 | 0.012 | -1.789 | [-0.097 - 0.009] | 0.000 | 1304 | 0.095 | 0.458 |
| Left Pericalcarine | 0.004 | 0.017 | 0.119 | [-0.067 - 0.075] | 0.000 | 1350 | 0.907 | 0.976 |
| Left Postcentral | -0.031 | 0.014 | -1.142 | [-0.090 - 0.028] | 0.000 | 1336 | 0.273 | 0.754 |
| Left Posterior Cingulate | -0.047 | 0.014 | -1.650 | [-0.108 - 0.014] | 0.000 | 1363 | 0.121 | 0.458 |
| Left Precentral | -0.040 | 0.015 | -1.365 | [-0.102 - 0.023] | 0.000 | 1344 | 0.194 | 0.601 |
| Left Precuneus | -0.023 | 0.015 | -0.752 | [-0.088 - 0.043] | 0.000 | 1337 | 0.464 | 0.867 |
| *Left Rostral Anterior Cingulate* | *0.052* | *0.011* | *2.240* | *[ 0.002 - 0.101]* | *0.000* | *1347* | *0.042* | *0.458* |
| Left Rostral Middle Frontal | -0.047 | 0.014 | -1.731 | [-0.106 - 0.011] | 0.000 | 1312 | 0.105 | 0.458 |
| *Left Superior Frontal* | *-0.067* | *0.011* | *-3.088* | *[-0.114 - -0.021]* | *0.000* | *1350* | *0.008* | *0.182* |
| Left Superior Parietal | -0.004 | 0.021 | -0.086 | [-0.093 - 0.086] | 45.814 | 1283 | 0.933 | 0.976 |
| Left Superior Temporal | -0.041 | 0.011 | -1.822 | [-0.089 - 0.007] | 0.000 | 1197 | 0.090 | 0.458 |
| Left Supramarginal | -0.052 | 0.014 | -1.877 | [-0.112 - 0.007] | 0.000 | 1216 | 0.082 | 0.458 |
| Left Frontal Pole | -0.008 | 0.013 | -0.288 | [-0.064 - 0.049] | 0.000 | 1368 | 0.777 | 0.927 |
| Left Temporal Pole | 0.011 | 0.011 | 0.509 | [-0.036 - 0.058] | 0.000 | 1329 | 0.619 | 0.867 |
| Left Transverse Temporal | 0.009 | 0.011 | 0.420 | [-0.037 - 0.054] | 0.000 | 1362 | 0.681 | 0.867 |
| Left Insula | -0.020 | 0.015 | -0.652 | [-0.086 - 0.046] | 4.239 | 1365 | 0.525 | 0.867 |
| Right Bankssts | -0.049 | 0.026 | -0.958 | [-0.159 - 0.061] | 70.254 | 1317 | 0.355 | 0.778 |
| Right Caudal Anterior Cingulate | -0.004 | 0.018 | -0.120 | [-0.082 - 0.073] | 0.000 | 1361 | 0.906 | 0.976 |
| Right Caudal Middle Frontal | -0.050 | 0.014 | -1.730 | [-0.111 - 0.012] | 0.000 | 1337 | 0.106 | 0.458 |
| Right Cuneus | -0.060 | 0.022 | -1.363 | [-0.154 - 0.035] | 56.887 | 1316 | 0.194 | 0.601 |
| Right Entorhinal | 0.033 | 0.016 | 1.033 | [-0.036 - 0.102] | 0.000 | 1164 | 0.319 | 0.775 |
| Right Fusiform | -0.008 | 0.013 | -0.311 | [-0.066 - 0.049] | 0.000 | 1281 | 0.760 | 0.923 |
| Right Inferior Parietal | -0.026 | 0.020 | -0.650 | [-0.113 - 0.061] | 0.000 | 1226 | 0.526 | 0.867 |
| Right Inferior Temporal | -0.013 | 0.013 | -0.508 | [-0.068 - 0.042] | 0.000 | 1283 | 0.619 | 0.867 |
| Right Isthmus Cingulate | -0.015 | 0.015 | -0.482 | [-0.080 - 0.051] | 0.000 | 1351 | 0.637 | 0.867 |
| Right Lateral Occipital | -0.015 | 0.016 | -0.461 | [-0.086 - 0.055] | 0.000 | 1287 | 0.652 | 0.867 |

| ROI | Partial *r* | SE | *t*-Score | 95% CI | *I2* | *N* | *p-value* | FDR |
|---|---|---|---|---|---|---|---|---|
| Right Lateral Orbitofrontal | -0.047 | 0.016 | -1.525 | [-0.114 - 0.019] | 0.000 | 1333 | 0.150 | 0.535 |
| Right Lingual | -0.022 | 0.014 | -0.785 | [-0.082 - 0.038] | 0.000 | 1350 | 0.445 | 0.867 |
| Right Medial Orbitofrontal | -0.049 | 0.014 | -1.690 | [-0.110 - 0.013] | 0.000 | 1316 | 0.113 | 0.458 |
| Right Middle Temporal | 0.001 | 0.019 | 0.037 | [-0.078 - 0.081] | 0.000 | 1255 | 0.971 | 0.982 |
| Right Parahippocampal | 0.019 | 0.011 | 0.840 | [-0.030 - 0.068] | 0.000 | 1322 | 0.415 | 0.867 |
| Right Paracentral | -0.018 | 0.016 | -0.547 | [-0.087 - 0.052] | 0.000 | 1361 | 0.593 | 0.867 |
| Right Pars Opercularis | -0.044 | 0.012 | -1.768 | [-0.096 - 0.009] | 0.000 | 1305 | 0.099 | 0.458 |
| Right Pars Orbitalis | -0.010 | 0.013 | -0.382 | [-0.067 - 0.047] | 0.000 | 1336 | 0.708 | 0.876 |
| Right Pars Triangularis | -0.054 | 0.013 | -2.128 | [-0.108 - 0.000] | 0.000 | 1305 | 0.052 | 0.458 |
| Right Pericalcarine | -0.013 | 0.016 | -0.410 | [-0.079 - 0.054] | 0.000 | 1350 | 0.688 | 0.867 |
| Right Postcentral | -0.013 | 0.012 | -0.521 | [-0.064 - 0.039] | 0.000 | 1342 | 0.610 | 0.867 |
| Right Posterior Cingulate | -0.007 | 0.016 | -0.227 | [-0.078 - 0.063] | 0.000 | 1364 | 0.824 | 0.950 |
| Right Precentral | -0.026 | 0.013 | -0.997 | [-0.080 - 0.029] | 0.000 | 1333 | 0.336 | 0.778 |
| Right Precuneus | 0.005 | 0.018 | 0.137 | [-0.073 - 0.083] | 0.000 | 1333 | 0.893 | 0.976 |
| Right Rostral Anterior Cingulate | 0.016 | 0.015 | 0.543 | [-0.047 - 0.079] | 0.000 | 1327 | 0.596 | 0.867 |
| Right Rostral Middle Frontal | -0.057 | 0.014 | -1.965 | [-0.118 - 0.005] | 0.000 | 1302 | 0.070 | 0.458 |
| *Right Superior Frontal* | *-0.069* | *0.011* | *-3.197* | *[-0.115 - -0.023]* | *0.000* | *1341* | *0.006* | *0.182* |
| Right Superior Parietal | -0.004 | 0.018 | -0.097 | [-0.083 - 0.076] | 30.079 | 1314 | 0.924 | 0.976 |
| Right Superior Temporal | -0.024 | 0.019 | -0.628 | [-0.106 - 0.058] | 0.000 | 1234 | 0.540 | 0.867 |
| Right Supramarginal | -0.038 | 0.015 | -1.268 | [-0.103 - 0.027] | 0.000 | 1223 | 0.225 | 0.666 |
| Right Frontal Pole | -0.015 | 0.010 | -0.710 | [-0.059 - 0.029] | 0.000 | 1365 | 0.489 | 0.867 |
| Right Temporal Pole | -0.007 | 0.007 | -0.439 | [-0.038 - 0.025] | 0.000 | 1225 | 0.667 | 0.867 |
| Right Transverse Temporal | -0.010 | 0.010 | -0.461 | [-0.054 - 0.035] | 0.000 | 1363 | 0.652 | 0.867 |
| Right Insula | -0.021 | 0.016 | -0.639 | [-0.089 - 0.048] | 0.000 | 1363 | 0.533 | 0.867 |
| Left Thickness | -0.035 | 0.013 | -1.330 | [-0.092 - 0.022] | 0.000 | 1370 | 0.205 | 1.000 |
| Right Thickness | -0.038 | 0.014 | -1.396 | [-0.096 - 0.020] | 0.000 | 1370 | 0.184 | 1.000 |
| Left Surface Area | -0.009 | 0.013 | -0.336 | [-0.066 - 0.048] | 0.000 | 1370 | 0.742 | 1.000 |
| Right Surface Area | -0.003 | 0.014 | -0.106 | [-0.063 - 0.057] | 0.000 | 1370 | 0.917 | 1.000 |
| Mean Bankssts | -0.042 | 0.020 | -1.058 | [-0.126 - 0.043] | 0.000 | 1230 | 0.308 | 0.633 |
| Mean Caudal Anterior Cingulate | 0.036 | 0.014 | 1.287 | [-0.024 - 0.095] | 0.000 | 1354 | 0.219 | 0.620 |
| Mean Caudal Middle Frontal | -0.019 | 0.014 | -0.665 | [-0.079 - 0.042] | 0.000 | 1323 | 0.517 | 0.703 |
| Mean Cuneus | -0.053 | 0.025 | -1.039 | [-0.160 - 0.056] | 66.200 | 1243 | 0.316 | 0.633 |
| Mean Entorhinal | 0.017 | 0.010 | 0.876 | [-0.025 - 0.059] | 0.000 | 1137 | 0.396 | 0.651 |
| Mean Fusiform | -0.036 | 0.012 | -1.465 | [-0.089 - 0.017] | 0.000 | 1240 | 0.165 | 0.564 |
| Mean Inferior Parietal | -0.025 | 0.018 | -0.673 | [-0.102 - 0.054] | 0.000 | 1160 | 0.512 | 0.703 |
| Mean Inferior Temporal | -0.002 | 0.012 | -0.101 | [-0.055 - 0.050] | 0.000 | 1227 | 0.921 | 0.921 |
| Mean Isthmus Cingulate | -0.023 | 0.014 | -0.813 | [-0.085 - 0.038] | 0.000 | 1345 | 0.430 | 0.664 |
| Mean Lateral Occipital | -0.031 | 0.017 | -0.938 | [-0.102 - 0.040] | 0.000 | 1237 | 0.364 | 0.651 |
| Mean Lateral Orbitofrontal | -0.036 | 0.015 | -1.220 | [-0.099 - 0.027] | 0.000 | 1327 | 0.243 | 0.633 |
| Mean Lingual | -0.038 | 0.016 | -1.172 | [-0.107 - 0.032] | 0.000 | 1328 | 0.261 | 0.633 |
| Mean Medial Orbitofrontal | -0.060 | 0.015 | -2.050 | [-0.123 - 0.003] | 0.000 | 1298 | 0.060 | 0.289 |
| Mean Middle Temporal | -0.014 | 0.016 | -0.425 | [-0.082 - 0.055] | 0.000 | 1162 | 0.677 | 0.874 |
| Mean Parahippocampal | 0.004 | 0.014 | 0.148 | [-0.057 - 0.065] | 0.000 | 1305 | 0.885 | 0.921 |
| Mean Paracentral | -0.006 | 0.016 | -0.187 | [-0.075 - 0.063] | 0.000 | 1352 | 0.854 | 0.921 |
| *Mean Pars Opercularis* | *-0.060* | *0.013* | *-2.309* | *[-0.116 - -0.004]* | *0.000* | *1273* | *0.037* | *0.289* |
| *Mean Pars Orbitalis* | *-0.063* | *0.012* | *-2.637* | *[-0.115 - -0.012]* | *0.000* | *1301* | *0.020* | *0.289* |
| *Mean Pars Triangularis* | *-0.065* | *0.013* | *-2.493* | *[-0.121 - -0.009]* | *0.000* | *1256* | *0.026* | *0.289* |
| Mean Pericalcarine | -0.005 | 0.015 | -0.175 | [-0.072 - 0.061] | 0.000 | 1340 | 0.864 | 0.921 |
| Mean Postcentral | -0.027 | 0.013 | -1.062 | [-0.081 - 0.027] | 0.000 | 1316 | 0.306 | 0.633 |
| Mean Posterior Cingulate | -0.028 | 0.016 | -0.883 | [-0.097 - 0.041] | 0.000 | 1358 | 0.392 | 0.651 |
| Mean Precentral | -0.044 | 0.013 | -1.651 | [-0.101 - 0.013] | 0.000 | 1318 | 0.121 | 0.514 |

| ROI | Partial *r* | SE | *t*-Score | 95% CI | I2 | N | *p-value* | FDR |
|---|---|---|---|---|---|---|---|---|
| Mean Precuneus | -0.036 | 0.021 | -0.864 | [-0.126 - 0.054] | 49.730 | 1311 | 0.402 | 0.651 |
| Mean Rostral Anterior Cingulate | 0.039 | 0.013 | 1.462 | [-0.018 - 0.095] | 0.000 | 1309 | 0.166 | 0.564 |
| *Mean Rostral Middle Frontal* | *-0.062* | *0.014* | *-2.176* | *[-0.123 - -0.001]* | *0.000* | *1263* | *0.047* | *0.289* |
| *Mean Superior Frontal* | *-0.070* | *0.011* | *-3.198* | *[-0.117 - -0.023]* | *0.000* | *1325* | *0.006* | *0.219* |
| Mean Superior Parietal | -0.009 | 0.021 | -0.219 | [-0.099 - 0.081] | 43.663 | 1248 | 0.830 | 0.921 |
| Mean Superior Temporal | -0.038 | 0.014 | -1.331 | [-0.098 - 0.023] | 0.000 | 1132 | 0.205 | 0.620 |
| Mean Supramarginal | -0.058 | 0.014 | -2.100 | [-0.117 - 0.001] | 0.000 | 1147 | 0.054 | 0.289 |
| Mean Frontal Pole | -0.008 | 0.011 | -0.366 | [-0.054 - 0.038] | 0.000 | 1363 | 0.720 | 0.874 |
| Mean Temporal Pole | 0.006 | 0.009 | 0.371 | [-0.031 - 0.043] | 0.000 | 1206 | 0.716 | 0.874 |
| Mean Transverse Temporal | -0.002 | 0.009 | -0.130 | [-0.042 - 0.038] | 0.000 | 1356 | 0.899 | 0.921 |
| Mean Insula | -0.021 | 0.016 | -0.669 | [-0.088 - 0.046] | 0.000 | 1358 | 0.515 | 0.703 |

Note: nominally significant results are italicized and FDR significant results are italicized and bolded

Supplementary Table S12 Associations between Cortical Thickness and SANS Alogia (Factor 5)

| ROI | Partial *r* | SE | *t*-Score | 95% CI | I2 | N | *p-value* | FDR |
|---|---|---|---|---|---|---|---|---|
| Left Bankssts | -0.044 | 0.015 | -1.452 | [-0.109 - 0.021] | 0.000 | 1266 | 0.168 | 0.327 |
| Left Caudal Anterior Cingulate | 0.035 | 0.014 | 1.240 | [-0.025 - 0.094] | 0.000 | 1362 | 0.235 | 0.370 |
| *Left Caudal Middle Frontal* | *-0.058* | *0.011* | *-2.550* | *[-0.107 - -0.009]* | *0.000* | *1351* | *0.023* | *0.117* |
| Left Cuneus | 0.003 | 0.016 | 0.092 | [-0.066 - 0.072] | 0.000 | 1275 | 0.928 | 0.954 |
| Left Entorhinal | -0.006 | 0.017 | -0.176 | [-0.079 - 0.067] | 22.784 | 1278 | 0.863 | 0.902 |
| *Left Fusiform* | *-0.078* | *0.016* | *-2.402* | *[-0.147 - -0.008]* | *0.000* | *1308* | *0.031* | *0.117* |
| Left Inferior Parietal | -0.042 | 0.015 | -1.380 | [-0.107 - 0.023] | 0.000 | 1234 | 0.189 | 0.348 |
| Left Inferior Temporal | -0.033 | 0.013 | -1.238 | [-0.089 - 0.024] | 0.000 | 1275 | 0.236 | 0.370 |
| Left Isthmus Cingulate | -0.024 | 0.011 | -1.126 | [-0.069 - 0.021] | 0.000 | 1361 | 0.279 | 0.383 |
| *Left Lateral Occipital* | *-0.053* | *0.011* | *-2.395* | *[-0.101 - -0.006]* | *0.000* | *1296* | *0.031* | *0.117* |
| Left Lateral Orbitofrontal | -0.036 | 0.015 | -1.177 | [-0.101 - 0.030] | 0.000 | 1364 | 0.259 | 0.382 |
| *Left Lingual* | *-0.068* | *0.011* | *-2.991* | *[-0.117 - -0.019]* | *0.000* | *1339* | *0.010* | *0.110* |
| Left Medial Orbitofrontal | -0.024 | 0.013 | -0.902 | [-0.082 - 0.033] | 0.000 | 1347 | 0.382 | 0.464 |
| Left Middle Temporal | -0.060 | 0.014 | -2.144 | [-0.119 - 0.000] | 0.000 | 1218 | 0.050 | 0.151 |
| Left Parahippocampal | -0.056 | 0.014 | -1.981 | [-0.116 - 0.005] | 0.000 | 1342 | 0.068 | 0.185 |
| Left Paracentral | -0.055 | 0.014 | -1.948 | [-0.116 - 0.006] | 0.000 | 1360 | 0.072 | 0.188 |
| *Left Pars Opercularis* | *-0.082* | *0.016* | *-2.497* | *[-0.151 - -0.012]* | *0.000* | *1323* | *0.026* | *0.117* |
| Left Pars Orbitalis | -0.026 | 0.012 | -1.122 | [-0.076 - 0.024] | 0.000 | 1327 | 0.281 | 0.383 |
| *Left Pars Triangularis* | *-0.079* | *0.017* | *-2.368* | *[-0.149 - -0.007]* | *0.000* | *1304* | *0.033* | *0.117* |
| Left Pericalcarine | 0.002 | 0.017 | 0.048 | [-0.070 - 0.074] | 0.000 | 1350 | 0.962 | 0.962 |
| Left Postcentral | -0.053 | 0.014 | -1.862 | [-0.113 - 0.008] | 0.000 | 1336 | 0.084 | 0.204 |
| *Left Posterior Cingulate* | *-0.073* | *0.015* | *-2.489* | *[-0.136 - -0.010]* | *0.000* | *1363* | *0.026* | *0.117* |
| *Left Precentral* | *-0.083* | *0.015* | *-2.698* | *[-0.147 - -0.017]* | *0.000* | *1344* | *0.017* | *0.117* |
| Left Precuneus | -0.056 | 0.018 | -1.505 | [-0.135 - 0.024] | 29.407 | 1337 | 0.154 | 0.309 |
| Left Rostral Anterior Cingulate | 0.016 | 0.017 | 0.480 | [-0.055 - 0.087] | 0.000 | 1347 | 0.639 | 0.689 |
| ***Left Rostral Middle Frontal*** | ***-0.080*** | ***0.011*** | ***-3.788*** | ***[-0.125 - -0.035]*** | ***0.000*** | ***1312*** | ***0.002*** | ***0.044*** |
| ***Left Superior Frontal*** | ***-0.101*** | ***0.010*** | ***-4.989*** | ***[-0.144 - -0.058]*** | ***0.000*** | ***1350*** | ***<0.001*** | ***0.014*** |
| Left Superior Parietal | -0.023 | 0.017 | -0.682 | [-0.096 - 0.050] | 8.394 | 1283 | 0.506 | 0.583 |
| *Left Superior Temporal* | *-0.100* | *0.015* | *-3.300* | *[-0.164 - -0.035]* | *0.000* | *1197* | *0.005* | *0.072* |
| *Left Supramarginal* | *-0.073* | *0.015* | *-2.376* | *[-0.138 - -0.007]* | *0.000* | *1216* | *0.032* | *0.117* |
| Left Frontal Pole | -0.029 | 0.013 | -1.098 | [-0.086 - 0.028] | 0.000 | 1368 | 0.291 | 0.388 |
| Left Temporal Pole | -0.008 | 0.008 | -0.480 | [-0.043 - 0.027] | 0.000 | 1329 | 0.639 | 0.689 |
| Left Transverse Temporal | -0.060 | 0.014 | -2.133 | [-0.120 - 0.000] | 0.000 | 1362 | 0.051 | 0.151 |
| Left Insula | -0.038 | 0.018 | -1.024 | [-0.116 - 0.041] | 11.502 | 1365 | 0.323 | 0.421 |
| Right Bankssts | -0.043 | 0.021 | -1.013 | [-0.134 - 0.048] | 28.028 | 1317 | 0.328 | 0.421 |

| ROI | Partial *r* | SE | *t*-Score | 95% CI | *I2* | *N* | *p-value* | FDR |
|---|---|---|---|---|---|---|---|---|
| Right Caudal Anterior Cingulate | 0.010 | 0.019 | 0.268 | [-0.072 - 0.092] | 0.000 | 1361 | 0.793 | 0.842 |
| ***Right Caudal Middle Frontal*** | ***-0.099*** | ***0.013*** | ***-3.656*** | ***[-0.156 - -0.041]*** | ***0.000*** | ***1337*** | ***0.003*** | ***0.044*** |
| Right Cuneus | -0.059 | 0.017 | -1.723 | [-0.132 - 0.015] | 0.000 | 1316 | 0.107 | 0.242 |
| Right Entorhinal | 0.045 | 0.023 | 0.968 | [-0.055 - 0.145] | 52.099 | 1164 | 0.349 | 0.432 |
| Right Fusiform | -0.034 | 0.015 | -1.139 | [-0.099 - 0.030] | 0.000 | 1281 | 0.274 | 0.383 |
| Right Inferior Parietal | -0.055 | 0.021 | -1.291 | [-0.146 - 0.037] | 47.376 | 1226 | 0.218 | 0.366 |
| Right Inferior Temporal | -0.042 | 0.015 | -1.363 | [-0.107 - 0.024] | 0.000 | 1283 | 0.194 | 0.348 |
| Right Isthmus Cingulate | -0.002 | 0.010 | -0.077 | [-0.045 - 0.042] | 0.000 | 1351 | 0.940 | 0.954 |
| Right Lateral Occipital | -0.052 | 0.014 | -1.893 | [-0.111 - 0.007] | 0.000 | 1287 | 0.079 | 0.199 |
| Right Lateral Orbitofrontal | -0.038 | 0.015 | -1.310 | [-0.100 - 0.024] | 0.000 | 1333 | 0.211 | 0.366 |
| *Right Lingual* | *-0.081* | *0.016* | *-2.581* | *[-0.147 - -0.014]* | *0.000* | *1350* | *0.022* | *0.117* |
| Right Medial Orbitofrontal | -0.029 | 0.015 | -0.990 | [-0.092 - 0.034] | 0.000 | 1316 | 0.339 | 0.427 |
| Right Middle Temporal | -0.029 | 0.017 | -0.880 | [-0.100 - 0.042] | 0.000 | 1255 | 0.394 | 0.470 |
| Right Parahippocampal | -0.024 | 0.018 | -0.660 | [-0.101 - 0.054] | 0.000 | 1322 | 0.520 | 0.589 |
| *Right Paracentral* | *-0.066* | *0.012* | *-2.794* | *[-0.116 - -0.015]* | *0.000* | *1361* | *0.014* | *0.117* |
| *Right Pars Opercularis* | *-0.079* | *0.015* | *-2.556* | *[-0.144 - -0.013]* | *0.000* | *1305* | *0.023* | *0.117* |
| Right Pars Orbitalis | -0.032 | 0.012 | -1.282 | [-0.085 - 0.021] | 0.000 | 1336 | 0.221 | 0.366 |
| Right Pars Triangularis | -0.056 | 0.014 | -1.978 | [-0.117 - 0.005] | 0.000 | 1305 | 0.068 | 0.185 |
| Right Pericalcarine | -0.062 | 0.017 | -1.773 | [-0.136 - 0.013] | 0.000 | 1350 | 0.098 | 0.230 |
| Right Postcentral | -0.040 | 0.013 | -1.548 | [-0.095 - 0.015] | 0.000 | 1342 | 0.144 | 0.296 |
| Right Posterior Cingulate | -0.034 | 0.012 | -1.400 | [-0.086 - 0.018] | 0.000 | 1364 | 0.183 | 0.346 |
| *Right Precentral* | *-0.065* | *0.012* | *-2.797* | *[-0.115 - -0.015]* | *0.000* | *1333* | *0.014* | *0.117* |
| Right Precuneus | -0.042 | 0.013 | -1.633 | [-0.097 - 0.013] | 0.000 | 1333 | 0.125 | 0.274 |
| Right Rostral Anterior Cingulate | -0.024 | 0.016 | -0.757 | [-0.094 - 0.045] | 0.000 | 1327 | 0.462 | 0.541 |
| Right Rostral Middle Frontal | -0.038 | 0.012 | -1.551 | [-0.091 - 0.015] | 0.000 | 1302 | 0.143 | 0.296 |
| ***Right Superior Frontal*** | ***-0.090*** | ***0.010*** | ***-4.452*** | ***[-0.134 - -0.047]*** | ***0.000*** | ***1341*** | ***0.001*** | ***0.019*** |
| Right Superior Parietal | -0.033 | 0.014 | -1.223 | [-0.091 - 0.025] | 0.000 | 1314 | 0.242 | 0.370 |
| *Right Superior Temporal* | *-0.075* | *0.015* | *-2.500* | *[-0.138 - -0.011]* | *0.000* | *1234* | *0.025* | *0.117* |
| *Right Supramarginal* | *-0.091* | *0.020* | *-2.248* | *[-0.176 - -0.004]* | *41.053* | *1223* | *0.041* | *0.134* |
| Right Frontal Pole | -0.029 | 0.013 | -1.119 | [-0.084 - 0.026] | 0.000 | 1365 | 0.282 | 0.383 |
| Right Temporal Pole | -0.020 | 0.016 | -0.608 | [-0.089 - 0.050] | 0.000 | 1225 | 0.553 | 0.616 |
| Right Transverse Temporal | -0.040 | 0.016 | -1.214 | [-0.109 - 0.030] | 0.000 | 1363 | 0.245 | 0.370 |
| *Right Insula* | *-0.066* | *0.015* | *-2.275* | *[-0.128 - -0.004]* | *0.000* | *1363* | *0.039* | *0.133* |
| *Left Thickness* | *-0.078* | *0.014* | *-2.820* | *[-0.136 - -0.019]* | *0.000* | *1370* | *0.014* | *1.000* |
| *Right Thickness* | *-0.084* | *0.012* | *-3.378* | *[-0.136 - -0.031]* | *0.000* | *1370* | *0.005* | *1.000* |
| Left Surface Area | -0.051 | 0.016 | -1.620 | [-0.118 - 0.017] | 0.000 | 1370 | 0.128 | 1.000 |
| Right Surface Area | -0.038 | 0.016 | -1.201 | [-0.106 - 0.030] | 0.000 | 1370 | 0.249 | 1.000 |
| Mean Bankssts | -0.056 | 0.019 | -1.514 | [-0.135 - 0.023] | 0.000 | 1230 | 0.152 | 0.273 |
| Mean Caudal Anterior Cingulate | 0.029 | 0.015 | 0.967 | [-0.035 - 0.092] | 0.000 | 1354 | 0.350 | 0.426 |
| ***Mean Caudal Middle Frontal*** | ***-0.082*** | ***0.012*** | ***-3.317*** | ***[-0.135 - -0.029]*** | ***0.000*** | ***1323*** | ***0.005*** | ***0.043*** |
| Mean Cuneus | -0.036 | 0.019 | -0.965 | [-0.116 - 0.044] | 0.000 | 1243 | 0.351 | 0.426 |
| Mean Entorhinal | 0.023 | 0.025 | 0.477 | [-0.082 - 0.128] | 57.586 | 1137 | 0.641 | 0.681 |
| Mean Fusiform | -0.063 | 0.016 | -1.985 | [-0.130 - 0.005] | 0.000 | 1240 | 0.067 | 0.175 |
| Mean Inferior Parietal | -0.054 | 0.021 | -1.279 | [-0.143 - 0.037] | 45.036 | 1160 | 0.222 | 0.314 |
| Mean Inferior Temporal | -0.030 | 0.013 | -1.122 | [-0.088 - 0.027] | 0.000 | 1227 | 0.281 | 0.382 |
| Mean Isthmus Cingulate | -0.017 | 0.010 | -0.856 | [-0.060 - 0.026] | 0.000 | 1345 | 0.406 | 0.476 |
| *Mean Lateral Occipital* | *-0.055* | *0.013* | *-2.157* | *[-0.109 - 0.000]* | *0.000* | *1237* | *0.049* | *0.138* |
| Mean Lateral Orbitofrontal | -0.041 | 0.015 | -1.325 | [-0.107 - 0.025] | 0.000 | 1327 | 0.206 | 0.305 |
| *Mean Lingual* | *-0.087* | *0.014* | *-3.064* | *[-0.147 - -0.026]* | *0.000* | *1328* | *0.008* | *0.057* |
| Mean Medial Orbitofrontal | -0.024 | 0.015 | -0.789 | [-0.088 - 0.041] | 0.000 | 1298 | 0.443 | 0.502 |
| Mean Middle Temporal | -0.045 | 0.017 | -1.336 | [-0.117 - 0.027] | 0.000 | 1162 | 0.203 | 0.305 |

| ROI | Partial *r* | SE | *t*-Score | 95% CI | *I2* | *N* | *p-value* | FDR |
|---|---|---|---|---|---|---|---|---|
| Mean Parahippocampal | -0.050 | 0.019 | -1.332 | [-0.129 - 0.030] | 38.397 | 1305 | 0.204 | 0.305 |
| *Mean Paracentral* | *-0.063* | *0.013* | *-2.386* | *[-0.120 - -0.006]* | *0.000* | *1352* | *0.032* | *0.108* |
| *Mean Pars Opercularis* | *-0.095* | *0.017* | *-2.720* | *[-0.169 - -0.020]* | *28.507* | *1273* | *0.017* | *0.070* |
| Mean Pars Orbitalis | -0.041 | 0.011 | -1.898 | [-0.088 - 0.005] | 0.000 | 1301 | 0.079 | 0.191 |
| *Mean Pars Triangularis* | *-0.087* | *0.015* | *-2.911* | *[-0.150 - -0.023]* | *0.000* | *1256* | *0.011* | *0.065* |
| Mean Pericalcarine | -0.035 | 0.018 | -0.987 | [-0.111 - 0.041] | 0.000 | 1340 | 0.341 | 0.426 |
| Mean Postcentral | -0.048 | 0.014 | -1.746 | [-0.106 - 0.011] | 0.000 | 1316 | 0.103 | 0.218 |
| *Mean Posterior Cingulate* | *-0.058* | *0.013* | *-2.304* | *[-0.112 - -0.004]* | *0.000* | *1358* | *0.037* | *0.115* |
| ***Mean Precentral*** | ***-0.088*** | ***0.013*** | ***-3.326*** | ***[-0.144 - -0.031]*** | ***0.000*** | ***1318*** | ***0.005*** | ***0.043*** |
| Mean Precuneus | -0.047 | 0.016 | -1.468 | [-0.115 - 0.022] | 2.046 | 1311 | 0.164 | 0.279 |
| Mean Rostral Anterior Cingulate | -0.004 | 0.018 | -0.111 | [-0.081 - 0.073] | 0.000 | 1309 | 0.913 | 0.913 |
| *Mean Rostral Middle Frontal* | *-0.063* | *0.011* | *-2.833* | *[-0.110 - -0.015]* | *0.000* | *1263* | *0.013* | *0.065* |
| ***Mean Superior Frontal*** | ***-0.098*** | ***0.010*** | ***-4.978*** | ***[-0.139 - -0.056]*** | ***0.000*** | ***1325*** | ***<0.001*** | ***0.007*** |
| Mean Superior Parietal | -0.025 | 0.017 | -0.756 | [-0.097 - 0.046] | 0.000 | 1248 | 0.462 | 0.507 |
| ***Mean Superior Temporal*** | ***-0.105*** | ***0.014*** | ***-3.780*** | ***[-0.164 - -0.046]*** | ***0.000*** | ***1132*** | ***0.002*** | ***0.034*** |
| *Mean Supramarginal* | *-0.083* | *0.017* | *-2.442* | *[-0.155 - -0.010]* | *0.000* | *1147* | *0.028* | *0.107* |
| Mean Frontal Pole | -0.032 | 0.010 | -1.622 | [-0.075 - 0.010] | 0.000 | 1363 | 0.127 | 0.245 |
| Mean Temporal Pole | -0.007 | 0.013 | -0.275 | [-0.065 - 0.050] | 0.000 | 1206 | 0.788 | 0.812 |
| Mean Transverse Temporal | -0.055 | 0.015 | -1.804 | [-0.119 - 0.010] | 0.000 | 1356 | 0.093 | 0.210 |
| Mean Insula | -0.056 | 0.017 | -1.610 | [-0.130 - 0.019] | 0.000 | 1358 | 0.130 | 0.245 |

Note: nominally significant results are italicized and FDR significant results are italicized and bolded

Supplementary Table S13 Associations between Subcortical Volumes and SANS Total

| ROI | Partial *r* | SE | *t*-score | CI | *I2* | *N* | *p*-value | FDR |
|---|---|---|---|---|---|---|---|---|
| *Left Lateral Ventricle* | *0.050* | *0.009* | *2.683* | *[ 0.010 - 0.090]* | *0.000* | *1355* | *0.018* | *0.143* |
| *Right Lateral Ventricle* | *0.061* | *0.011* | *2.768* | *[ 0.014 - 0.109]* | *0.000* | *1355* | *0.015* | *0.143* |
| Left Thalamus | -0.022 | 0.016 | -0.696 | [-0.089 - 0.046] | 0.000 | 1344 | 0.498 | 0.612 |
| Right Thalamus | -0.047 | 0.020 | -1.187 | [-0.132 - 0.038] | 12.938 | 1347 | 0.255 | 0.461 |
| Left Caudate | 0.033 | 0.014 | 1.128 | [-0.029 - 0.094] | 0.000 | 1345 | 0.278 | 0.461 |
| Right Caudate | 0.028 | 0.013 | 1.054 | [-0.029 - 0.086] | 0.000 | 1348 | 0.310 | 0.461 |
| Left Putamen | 0.012 | 0.014 | 0.410 | [-0.049 - 0.072] | 0.000 | 1347 | 0.688 | 0.787 |
| Right Putamen | -0.006 | 0.014 | -0.212 | [-0.066 - 0.054] | 0.000 | 1347 | 0.835 | 0.890 |
| Left Pallidum | 0.032 | 0.023 | 0.698 | [-0.066 - 0.129] | 74.360 | 1341 | 0.497 | 0.612 |
| Right Pallidum | 0.043 | 0.020 | 1.037 | [-0.045 - 0.130] | 54.829 | 1348 | 0.317 | 0.461 |
| Left Hippocampus | -0.043 | 0.015 | -1.450 | [-0.107 - 0.021] | 0.000 | 1339 | 0.169 | 0.451 |
| Right Hippocampus | -0.048 | 0.013 | -1.818 | [-0.104 - 0.009] | 0.000 | 1349 | 0.091 | 0.290 |
| Left Amygdala | -0.003 | 0.015 | -0.109 | [-0.066 - 0.060] | 0.000 | 1345 | 0.915 | 0.915 |
| Right Amygdala | -0.051 | 0.022 | -1.162 | [-0.145 - 0.044] | 41.363 | 1349 | 0.265 | 0.461 |
| Left Accumbens | -0.072 | 0.018 | -2.038 | [-0.147 - 0.004] | 0.000 | 1329 | 0.061 | 0.256 |
| Right Accumbens | -0.059 | 0.015 | -2.011 | [-0.121 - 0.004] | 0.000 | 1335 | 0.064 | 0.256 |
| *Mean Lateral Ventricle* | *0.057* | *0.010* | *2.838* | *[ 0.014 - 0.100]* | *0.000* | *1355* | *0.013* | *0.093* |

| | | | | | | | | |
|---|---|---|---|---|---|---|---|---|
| Mean Thalamus | -0.033 | 0.017 | -0.943 | [-0.107 - 0.042] | 0.000 | 1342 | 0.362 | 0.413 |
| Mean Caudate | 0.031 | 0.013 | 1.159 | [-0.026 - 0.088] | 0.000 | 1343 | 0.266 | 0.413 |
| Mean Putamen | 0.005 | 0.014 | 0.180 | [-0.055 - 0.065] | 0.000 | 1341 | 0.860 | 0.860 |
| Mean Pallidum | 0.040 | 0.020 | 1.012 | [-0.045 - 0.123] | 0.000 | 1335 | 0.329 | 0.413 |
| Mean Hippocampus | -0.043 | 0.014 | -1.561 | [-0.102 - 0.016] | 0.000 | 1335 | 0.141 | 0.376 |
| Mean Amygdala | -0.035 | 0.017 | -1.014 | [-0.110 - 0.039] | 0.000 | 1341 | 0.328 | 0.413 |
| *Mean Accumbens* | *-0.078* | *0.015* | *-2.545* | *[-0.143 - -0.012]* | *0.000* | *1318* | *0.023* | *0.093* |

Note: nominally significant results are italicized and FDR significant results are italicized and bolded

Supplementary Table S14 Associations between Subcortical Volumes and SANS MAP

| ROI | Partial *r* | SE | *t*-score | CI | *I*2 | *N* | *p*-value | FDR |
|---|---|---|---|---|---|---|---|---|
| Left Lateral Ventricle | 0.031 | 0.012 | 1.320 | [-0.020 - 0.082] | 0.000 | 1355 | 0.208 | 0.380 |
| Right Lateral Ventricle | 0.040 | 0.016 | 1.283 | [-0.027 - 0.107] | 0.000 | 1355 | 0.220 | 0.380 |
| Left Thalamus | -0.007 | 0.012 | -0.308 | [-0.058 - 0.044] | 0.000 | 1344 | 0.763 | 0.763 |
| Right Thalamus | -0.038 | 0.016 | -1.233 | [-0.105 - 0.028] | 0.000 | 1347 | 0.238 | 0.380 |
| Left Caudate | 0.029 | 0.014 | 1.052 | [-0.030 - 0.088] | 0.000 | 1345 | 0.311 | 0.452 |
| Right Caudate | 0.025 | 0.013 | 0.983 | [-0.029 - 0.078] | 0.000 | 1348 | 0.342 | 0.456 |
| Left Putamen | 0.022 | 0.014 | 0.787 | [-0.037 - 0.081] | 0.000 | 1347 | 0.445 | 0.547 |
| Right Putamen | 0.017 | 0.012 | 0.715 | [-0.035 - 0.069] | 0.000 | 1347 | 0.487 | 0.556 |
| Left Pallidum | 0.028 | 0.021 | 0.657 | [-0.063 - 0.118] | 68.092 | 1341 | 0.522 | 0.556 |
| Right Pallidum | 0.059 | 0.016 | 1.891 | [-0.008 - 0.126] | 0.000 | 1348 | 0.080 | 0.267 |
| Left Hippocampus | -0.041 | 0.015 | -1.405 | [-0.103 - 0.022] | 0.000 | 1339 | 0.182 | 0.380 |
| Right Hippocampus | -0.039 | 0.015 | -1.280 | [-0.105 - 0.027] | 0.000 | 1349 | 0.221 | 0.380 |
| Left Amygdala | -0.048 | 0.013 | -1.863 | [-0.102 - 0.007] | 0.000 | 1345 | 0.084 | 0.267 |
| Right Amygdala | -0.075 | 0.019 | -2.002 | [-0.155 - 0.005] | 15.320 | 1349 | 0.065 | 0.267 |
| Left Accumbens | -0.065 | 0.016 | -2.018 | [-0.134 - 0.004] | 0.000 | 1329 | 0.063 | 0.267 |
| *Right Accumbens* | *-0.049* | *0.011* | *-2.246* | *[-0.096 - -0.002]* | *0.000* | *1335* | *0.041* | *0.267* |
| Mean Lateral Ventricle | 0.037 | 0.014 | 1.346 | [-0.022 - 0.096] | 0.000 | 1355 | 0.200 | 0.434 |
| Mean Thalamus | -0.021 | 0.014 | -0.753 | [-0.079 - 0.038] | 0.000 | 1342 | 0.464 | 0.464 |
| Mean Caudate | 0.026 | 0.013 | 1.019 | [-0.029 - 0.080] | 0.000 | 1343 | 0.326 | 0.434 |
| Mean Putamen | 0.022 | 0.014 | 0.803 | [-0.037 - 0.081] | 0.000 | 1341 | 0.435 | 0.464 |
| Mean Pallidum | 0.039 | 0.019 | 1.026 | [-0.042 - 0.119] | 0.000 | 1335 | 0.322 | 0.434 |
| Mean Hippocampus | -0.038 | 0.015 | -1.289 | [-0.102 - 0.025] | 0.000 | 1335 | 0.218 | 0.434 |
| Mean Amygdala | -0.069 | 0.016 | -2.118 | [-0.137 - 0.001] | 0.000 | 1341 | 0.053 | 0.210 |
| *Mean Accumbens* | *-0.066* | *0.013* | *-2.469* | *[-0.123 - -0.009]* | *0.000* | *1318* | *0.027* | *0.210* |

Note: nominally significant results are italicized and FDR significant results are italicized and bolded

Supplementary Table S15 Associations between Subcortical Volumes and SANS Anhedonia (Factor 1)

| ROI | Partial *r* | SE | *t*-score | CI | *I*2 | *N* | *p*-value | FDR |
|---|---|---|---|---|---|---|---|---|
| Left Lateral Ventricle | 0.010 | 0.012 | 0.424 | [-0.041 - 0.061] | 0.000 | 1355 | 0.678 | 0.933 |
| Right Lateral Ventricle | 0.026 | 0.015 | 0.860 | [-0.038 - 0.089] | 0.000 | 1355 | 0.404 | 0.905 |
| Left Thalamus | 0.004 | 0.016 | 0.121 | [-0.063 - 0.070] | 0.000 | 1344 | 0.905 | 0.933 |
| Right Thalamus | -0.026 | 0.019 | -0.677 | [-0.108 - 0.056] | 0.000 | 1347 | 0.509 | 0.905 |
| Left Caudate | 0.024 | 0.015 | 0.811 | [-0.039 - 0.087] | 0.000 | 1345 | 0.431 | 0.905 |
| Right Caudate | 0.021 | 0.014 | 0.751 | [-0.040 - 0.083] | 0.000 | 1348 | 0.465 | 0.905 |
| Left Putamen | 0.003 | 0.016 | 0.086 | [-0.068 - 0.073] | 0.000 | 1347 | 0.933 | 0.933 |
| Right Putamen | -0.007 | 0.020 | -0.167 | [-0.094 - 0.081] | 36.450 | 1347 | 0.870 | 0.933 |
| Left Pallidum | 0.005 | 0.018 | 0.152 | [-0.070 - 0.081] | 0.000 | 1341 | 0.881 | 0.933 |
| Right Pallidum | 0.043 | 0.015 | 1.453 | [-0.020 - 0.106] | 0.000 | 1348 | 0.168 | 0.905 |
| Left Hippocampus | -0.013 | 0.017 | -0.380 | [-0.087 - 0.061] | 0.000 | 1339 | 0.710 | 0.933 |
| Right Hippocampus | -0.015 | 0.020 | -0.386 | [-0.100 - 0.069] | 0.000 | 1349 | 0.705 | 0.933 |
| Left Amygdala | -0.029 | 0.014 | -1.020 | [-0.090 - 0.032] | 0.000 | 1345 | 0.325 | 0.905 |
| Right Amygdala | -0.040 | 0.018 | -1.122 | [-0.117 - 0.037] | 0.000 | 1349 | 0.281 | 0.905 |
| Left Accumbens | -0.029 | 0.020 | -0.727 | [-0.115 - 0.057] | 23.432 | 1329 | 0.479 | 0.905 |
| Right Accumbens | -0.045 | 0.016 | -1.427 | [-0.113 - 0.023] | 0.000 | 1335 | 0.176 | 0.905 |
| Mean Lateral Ventricle | 0.019 | 0.013 | 0.717 | [-0.038 - 0.076] | 0.000 | 1355 | 0.485 | 0.815 |
| Mean Thalamus | -0.008 | 0.018 | -0.233 | [-0.083 - 0.067] | 0.000 | 1342 | 0.819 | 0.936 |
| Mean Caudate | 0.024 | 0.014 | 0.841 | [-0.037 - 0.085] | 0.000 | 1343 | 0.415 | 0.815 |
| Mean Putamen | 0.000 | 0.018 | 0.010 | [-0.076 - 0.077] | 0.000 | 1341 | 0.992 | 0.992 |
| Mean Pallidum | 0.023 | 0.017 | 0.677 | [-0.049 - 0.095] | 0.000 | 1335 | 0.510 | 0.815 |
| Mean Hippocampus | -0.014 | 0.018 | -0.369 | [-0.092 - 0.065] | 0.000 | 1335 | 0.717 | 0.936 |
| Mean Amygdala | -0.040 | 0.017 | -1.163 | [-0.113 - 0.034] | 0.000 | 1341 | 0.264 | 0.815 |
| Mean Accumbens | -0.046 | 0.018 | -1.290 | [-0.121 - 0.030] | 1.414 | 1318 | 0.218 | 0.815 |

Note: nominally significant results are italicized and FDR significant results are italicized and bolded

Supplementary Table S16 Associations between Subcortical Volumes and SANS Asociality (Factor 2)

| ROI | Partial *r* | SE | *t*-score | CI | *I*2 | *N* | *p*-value | FDR |
|---|---|---|---|---|---|---|---|---|
| Left Lateral Ventricle | 0.025 | 0.011 | 1.081 | [-0.024 - 0.073] | 0.000 | 1355 | 0.298 | 0.476 |
| Right Lateral Ventricle | 0.034 | 0.015 | 1.148 | [-0.029 - 0.097] | 0.000 | 1355 | 0.270 | 0.476 |
| Left Thalamus | 0.013 | 0.013 | 0.504 | [-0.042 - 0.069] | 0.000 | 1344 | 0.622 | 0.765 |
| Right Thalamus | -0.011 | 0.015 | -0.358 | [-0.077 - 0.055] | 0.000 | 1347 | 0.726 | 0.830 |
| Left Caudate | 0.036 | 0.016 | 1.145 | [-0.032 - 0.103] | 0.000 | 1345 | 0.271 | 0.476 |
| Right Caudate | 0.029 | 0.014 | 1.015 | [-0.032 - 0.089] | 0.000 | 1348 | 0.327 | 0.476 |
| Left Putamen | 0.054 | 0.016 | 1.715 | [-0.014 - 0.121] | 0.000 | 1347 | 0.108 | 0.355 |
| Right Putamen | 0.042 | 0.013 | 1.552 | [-0.016 - 0.099] | 0.000 | 1347 | 0.143 | 0.381 |
| Left Pallidum | 0.035 | 0.023 | 0.742 | [-0.065 - 0.134] | 0.000 | 1341 | 0.470 | 0.627 |
| Right Pallidum | 0.089 | 0.026 | 1.702 | [-0.023 - 0.198] | 78.521 | 1348 | 0.111 | 0.355 |
| Left Hippocampus | -0.005 | 0.015 | -0.156 | [-0.068 - 0.058] | 0.000 | 1339 | 0.878 | 0.937 |

| | | | | | | | | |
|---|---|---|---|---|---|---|---|---|
| Right Hippocampus | 0.002 | 0.017 | 0.047 | [-0.069 - 0.072] | 0.000 | 1349 | 0.963 | 0.963 |
| Left Amygdala | -0.029 | 0.013 | -1.105 | [-0.085 - 0.027] | 0.000 | 1345 | 0.288 | 0.476 |
| Right Amygdala | -0.064 | 0.016 | -1.987 | [-0.133 - 0.005] | 0.000 | 1349 | 0.067 | 0.355 |
| Left Accumbens | -0.071 | 0.017 | -2.046 | [-0.144 - 0.003] | 0.000 | 1329 | 0.060 | 0.355 |
| *Right Accumbens* | *-0.047* | *0.010* | *-2.375* | *[-0.090 - -0.005]* | *0.000* | *1335* | *0.032* | *0.355* |
| Mean Lateral Ventricle | 0.030 | 0.013 | 1.164 | [-0.025 - 0.086] | 0.000 | 1355 | 0.264 | 0.423 |
| Mean Thalamus | 0.006 | 0.014 | 0.193 | [-0.056 - 0.067] | 0.000 | 1342 | 0.850 | 0.926 |
| Mean Caudate | 0.031 | 0.015 | 1.038 | [-0.033 - 0.096] | 0.000 | 1343 | 0.317 | 0.423 |
| Mean Putamen | 0.051 | 0.014 | 1.755 | [-0.011 - 0.112] | 0.000 | 1341 | 0.101 | 0.286 |
| Mean Pallidum | 0.056 | 0.026 | 1.055 | [-0.058 - 0.168] | 0.000 | 1335 | 0.309 | 0.423 |
| Mean Hippocampus | 0.003 | 0.016 | 0.095 | [-0.067 - 0.073] | 41.150 | 1335 | 0.926 | 0.926 |
| Mean Amygdala | -0.051 | 0.015 | -1.721 | [-0.115 - 0.013] | 0.000 | 1341 | 0.107 | 0.286 |
| *Mean Accumbens* | *-0.065* | *0.014* | *-2.308* | *[-0.126 - -0.005]* | *0.000* | *1318* | *0.037* | *0.286* |

Supplementary Table S17 Associations between Subcortical Volumes and SANS Avolition (Factor 3)

| ROI | Partial *r* | SE | *t*-score | CI | *I*2 | *N* | *p*-value | FDR |
|---|---|---|---|---|---|---|---|---|
| Left Lateral Ventricle | 0.044 | 0.014 | 1.571 | [-0.016 - 0.103] | 0.000 | 1355 | 0.138 | 0.277 |
| Right Lateral Ventricle | 0.041 | 0.015 | 1.406 | [-0.022 - 0.103] | 0.000 | 1355 | 0.181 | 0.323 |
| Left Thalamus | -0.039 | 0.011 | -1.758 | [-0.085 - 0.008] | 0.000 | 1344 | 0.101 | 0.230 |
| *Right Thalamus* | *-0.069* | *0.012* | *-2.776* | *[-0.122 - -0.016]* | *0.000* | *1347* | *0.015* | *0.059* |
| Left Caudate | 0.008 | 0.012 | 0.331 | [-0.044 - 0.060] | 0.000 | 1345 | 0.746 | 0.796 |
| Right Caudate | 0.006 | 0.010 | 0.330 | [-0.035 - 0.047] | 0.000 | 1348 | 0.746 | 0.796 |
| Left Putamen | -0.019 | 0.013 | -0.764 | [-0.073 - 0.035] | 0.000 | 1347 | 0.457 | 0.563 |
| Right Putamen | -0.015 | 0.008 | -1.005 | [-0.048 - 0.017] | 0.000 | 1347 | 0.332 | 0.471 |
| Left Pallidum | -0.006 | 0.011 | -0.257 | [-0.054 - 0.043] | 0.000 | 1341 | 0.801 | 0.801 |
| Right Pallidum | 0.015 | 0.008 | 0.960 | [-0.019 - 0.049] | 0.000 | 1348 | 0.353 | 0.471 |
| ***Left Hippocampus*** | ***-0.092*** | ***0.011*** | ***-4.232*** | ***[-0.138 - -0.045]*** | ***0.000*** | ***1339*** | ***0.001*** | ***0.007*** |
| ***Right Hippocampus*** | ***-0.096*** | ***0.011*** | ***-4.317*** | ***[-0.142 - -0.048]*** | ***0.000*** | ***1349*** | ***0.001*** | ***0.007*** |
| *Left Amygdala* | *-0.064* | *0.012* | *-2.784* | *[-0.114 - -0.015]* | *0.000* | *1345* | *0.015* | *0.059* |
| *Right Amygdala* | *-0.080* | *0.016* | *-2.437* | *[-0.150 - -0.010]* | *0.000* | *1349* | *0.029* | *0.092* |
| Left Accumbens | -0.054 | 0.014 | -1.922 | [-0.113 - 0.006] | 0.000 | 1329 | 0.075 | 0.200 |
| Right Accumbens | -0.032 | 0.015 | -1.099 | [-0.095 - 0.031] | 0.000 | 1335 | 0.290 | 0.465 |
| Mean Lateral Ventricle | 0.044 | 0.014 | 1.541 | [-0.017 - 0.104] | 0.000 | 1355 | 0.146 | 0.233 |
| *Mean Thalamus* | *-0.056* | *0.012* | *-2.401* | *[-0.105 - -0.006]* | *0.000* | *1342* | *0.031* | *0.082* |
| Mean Caudate | 0.004 | 0.011 | 0.205 | [-0.042 - 0.050] | 0.000 | 1343 | 0.840 | 0.854 |
| Mean Putamen | -0.015 | 0.011 | -0.646 | [-0.064 - 0.034] | 0.000 | 1341 | 0.529 | 0.705 |
| Mean Pallidum | 0.004 | 0.011 | 0.187 | [-0.041 - 0.049] | 0.000 | 1335 | 0.854 | 0.854 |
| ***Mean Hippocampus*** | ***-0.096*** | ***0.011*** | ***-4.522*** | ***[-0.141 - -0.051]*** | ***0.000*** | ***1335*** | ***<0.001*** | ***0.004*** |
| *Mean Amygdala* | *-0.079* | *0.014* | *-2.774* | *[-0.139 - -0.018]* | *0.000* | *1341* | *0.015* | *0.060* |
| Mean Accumbens | -0.054 | 0.014 | -1.971 | [-0.112 - 0.005] | 0.000 | 1318 | 0.069 | 0.138 |

Note: nominally significant results are italicized and FDR significant results are italicized and bolded

Supplementary Table S18 Associations between Subcortical Volumes and SANS EXP

| ROI | Partial *r* | SE | *t*-score | CI | *I*2 | *N* | *p*-value | FDR |
|---|---|---|---|---|---|---|---|---|
| Left Lateral Ventricle | 0.043 | 0.013 | 1.646 | [-0.013 - 0.098] | 0.000 | 1355 | 0.122 | 0.578 |
| Right Lateral Ventricle | 0.055 | 0.014 | 2.034 | [-0.003 - 0.113] | 0.000 | 1355 | 0.061 | 0.578 |
| Left Thalamus | -0.023 | 0.017 | -0.683 | [-0.095 - 0.049] | 0.000 | 1344 | 0.506 | 0.578 |
| Right Thalamus | -0.047 | 0.023 | -1.011 | [-0.144 - 0.052] | 53.318 | 1347 | 0.329 | 0.578 |
| Left Caudate | 0.027 | 0.014 | 0.946 | [-0.034 - 0.088] | 0.000 | 1345 | 0.360 | 0.578 |
| Right Caudate | 0.025 | 0.014 | 0.901 | [-0.034 - 0.083] | 0.000 | 1348 | 0.383 | 0.578 |
| Left Putamen | -0.002 | 0.016 | -0.072 | [-0.071 - 0.067] | 0.000 | 1347 | 0.943 | 0.943 |
| Right Putamen | -0.030 | 0.016 | -0.941 | [-0.099 - 0.039] | 0.000 | 1347 | 0.362 | 0.578 |
| Left Pallidum | 0.028 | 0.018 | 0.782 | [-0.049 - 0.105] | 0.000 | 1341 | 0.447 | 0.578 |
| Right Pallidum | 0.025 | 0.018 | 0.720 | [-0.050 - 0.101] | 0.000 | 1348 | 0.483 | 0.578 |
| Left Hippocampus | -0.027 | 0.015 | -0.864 | [-0.092 - 0.039] | 0.000 | 1339 | 0.402 | 0.578 |
| Right Hippocampus | -0.033 | 0.013 | -1.266 | [-0.090 - 0.023] | 0.000 | 1349 | 0.226 | 0.578 |
| Left Amygdala | 0.041 | 0.015 | 1.375 | [-0.023 - 0.105] | 0.000 | 1345 | 0.191 | 0.578 |
| Right Amygdala | -0.016 | 0.022 | -0.367 | [-0.108 - 0.077] | 46.148 | 1349 | 0.719 | 0.767 |
| Left Accumbens | -0.062 | 0.016 | -1.884 | [-0.131 - 0.009] | 0.000 | 1329 | 0.081 | 0.578 |
| Right Accumbens | -0.045 | 0.016 | -1.431 | [-0.112 - 0.022] | 16.963 | 1335 | 0.174 | 0.578 |
| Mean Lateral Ventricle | 0.050 | 0.013 | 1.859 | [-0.008 - 0.107] | 0.000 | 1355 | 0.084 | 0.336 |
| Mean Thalamus | -0.030 | 0.019 | -0.792 | [-0.110 - 0.051] | 0.000 | 1342 | 0.442 | 0.589 |
| Mean Caudate | 0.029 | 0.014 | 1.052 | [-0.030 - 0.087] | 0.000 | 1343 | 0.311 | 0.589 |
| Mean Putamen | -0.015 | 0.016 | -0.454 | [-0.083 - 0.054] | 0.000 | 1341 | 0.657 | 0.750 |
| Mean Pallidum | 0.031 | 0.018 | 0.861 | [-0.047 - 0.109] | 0.000 | 1335 | 0.403 | 0.589 |
| Mean Hippocampus | -0.027 | 0.014 | -0.979 | [-0.087 - 0.032] | 0.000 | 1335 | 0.344 | 0.589 |
| Mean Amygdala | 0.007 | 0.017 | 0.217 | [-0.065 - 0.079] | 0.000 | 1341 | 0.831 | 0.831 |
| *Mean Accumbens* | *-0.068* | *0.015* | *-2.253* | *[-0.133 - -0.003]* | *0.000* | *1318* | *0.041* | *0.327* |

Note: nominally significant results are italicized and FDR significant results are italicized and bolded

Supplementary Table S19 Associations between Subcortical Volumes and SANS Blunted Affect (Factor 4)

| ROI | Partial *r* | SE | *t*-score | CI | *I*2 | *N* | *p*-value | FDR |
|---|---|---|---|---|---|---|---|---|
| Left Lateral Ventricle | 0.041 | 0.014 | 1.527 | [-0.017 - 0.099] | 0.000 | 1355 | 0.149 | 0.597 |
| Right Lateral Ventricle | 0.053 | 0.014 | 1.920 | [-0.006 - 0.112] | 0.000 | 1355 | 0.076 | 0.597 |
| Left Thalamus | -0.010 | 0.018 | -0.296 | [-0.086 - 0.065] | 0.000 | 1344 | 0.772 | 0.882 |
| Right Thalamus | -0.040 | 0.025 | -0.803 | [-0.145 - 0.066] | 66.436 | 1347 | 0.435 | 0.674 |
| Left Caudate | 0.031 | 0.014 | 1.125 | [-0.028 - 0.090] | 0.000 | 1345 | 0.279 | 0.674 |
| Right Caudate | 0.029 | 0.014 | 1.025 | [-0.031 - 0.088] | 0.000 | 1348 | 0.323 | 0.674 |
| Left Putamen | 0.004 | 0.017 | 0.110 | [-0.070 - 0.077] | 0.000 | 1347 | 0.914 | 0.914 |
| Right Putamen | -0.026 | 0.017 | -0.773 | [-0.099 - 0.046] | 0.000 | 1347 | 0.452 | 0.674 |

| | | | | | | | | |
|---|---|---|---|---|---|---|---|---|
| Left Pallidum | 0.029 | 0.020 | 0.722 | [-0.057 - 0.115] | 0.000 | 1341 | 0.482 | 0.674 |
| Right Pallidum | 0.027 | 0.019 | 0.716 | [-0.053 - 0.106] | 0.000 | 1348 | 0.486 | 0.674 |
| Left Hippocampus | -0.013 | 0.015 | -0.433 | [-0.080 - 0.053] | 0.000 | 1339 | 0.672 | 0.827 |
| Right Hippocampus | -0.019 | 0.014 | -0.684 | [-0.080 - 0.042] | 0.000 | 1349 | 0.505 | 0.674 |
| Left Amygdala | 0.053 | 0.016 | 1.695 | [-0.014 - 0.119] | 0.000 | 1345 | 0.112 | 0.597 |
| Right Amygdala | -0.008 | 0.023 | -0.172 | [-0.107 - 0.091] | 59.701 | 1349 | 0.866 | 0.914 |
| Left Accumbens | -0.065 | 0.017 | -1.935 | [-0.137 - 0.007] | 0.000 | 1329 | 0.073 | 0.597 |
| Right Accumbens | -0.037 | 0.016 | -1.146 | [-0.107 - 0.033] | 28.732 | 1335 | 0.271 | 0.674 |
| Mean Lateral Ventricle | 0.048 | 0.014 | 1.741 | [-0.011 - 0.107] | 0.000 | 1355 | 0.104 | 0.414 |
| Mean Thalamus | -0.018 | 0.020 | -0.443 | [-0.103 - 0.068] | 8.355 | 1342 | 0.664 | 0.712 |
| Mean Caudate | 0.033 | 0.014 | 1.219 | [-0.025 - 0.091] | 0.000 | 1343 | 0.243 | 0.648 |
| Mean Putamen | -0.014 | 0.018 | -0.377 | [-0.092 - 0.064] | 18.043 | 1341 | 0.712 | 0.712 |
| Mean Pallidum | 0.033 | 0.020 | 0.824 | [-0.052 - 0.117] | 0.000 | 1335 | 0.424 | 0.712 |
| Mean Hippocampus | -0.013 | 0.014 | -0.464 | [-0.075 - 0.048] | 0.000 | 1335 | 0.650 | 0.712 |
| Mean Amygdala | 0.023 | 0.019 | 0.611 | [-0.058 - 0.104] | 28.867 | 1341 | 0.551 | 0.712 |
| Mean Accumbens | -0.064 | 0.016 | -1.989 | [-0.132 - 0.005] | 11.193 | 1318 | 0.067 | 0.414 |

Note: nominally significant results are italicized and FDR significant results are italicized and bolded

Supplementary Table S20 Associations between Subcortical Volumes and SANS Alogia

| ROI | Partial *r* | SE | *t*-score | CI | *I*2 | *N* | *p*-value | FDR |
|---|---|---|---|---|---|---|---|---|
| Left Lateral Ventricle | 0.034 | 0.012 | 1.474 | [-0.016 - 0.084] | 0.000 | 1355 | 0.163 | 0.325 |
| Right Lateral Ventricle | 0.045 | 0.013 | 1.740 | [-0.010 - 0.100] | 0.000 | 1355 | 0.104 | 0.237 |
| Left Thalamus | -0.054 | 0.015 | -1.813 | [-0.118 - 0.010] | 0.000 | 1344 | 0.091 | 0.237 |
| Right Thalamus | -0.064 | 0.016 | -2.059 | [-0.131 - 0.003] | 0.000 | 1347 | 0.059 | 0.237 |
| Left Caudate | 0.003 | 0.015 | 0.112 | [-0.060 - 0.067] | 0.000 | 1345 | 0.913 | 0.962 |
| Right Caudate | 0.001 | 0.013 | 0.048 | [-0.053 - 0.055] | 0.000 | 1348 | 0.962 | 0.962 |
| Left Putamen | -0.022 | 0.015 | -0.736 | [-0.087 - 0.043] | 0.000 | 1347 | 0.474 | 0.689 |
| Right Putamen | -0.033 | 0.015 | -1.127 | [-0.097 - 0.030] | 0.000 | 1347 | 0.279 | 0.446 |
| Left Pallidum | 0.013 | 0.015 | 0.424 | [-0.051 - 0.077] | 0.000 | 1341 | 0.678 | 0.784 |
| Right Pallidum | 0.016 | 0.015 | 0.541 | [-0.047 - 0.078] | 0.000 | 1348 | 0.597 | 0.784 |
| Left Hippocampus | -0.060 | 0.016 | -1.876 | [-0.128 - 0.009] | 0.000 | 1339 | 0.082 | 0.237 |
| *Right Hippocampus* | *-0.066* | *0.012* | *-2.778* | *[-0.116 - -0.015]* | *0.000* | *1349* | *0.015* | *0.237* |
| Left Amygdala | -0.009 | 0.011 | -0.413 | [-0.059 - 0.040] | 0.000 | 1345 | 0.686 | 0.784 |
| Right Amygdala | -0.054 | 0.015 | -1.836 | [-0.116 - 0.009] | 0.000 | 1349 | 0.088 | 0.237 |
| Left Accumbens | -0.028 | 0.012 | -1.154 | [-0.081 - 0.024] | 0.000 | 1329 | 0.268 | 0.446 |
| Right Accumbens | -0.052 | 0.014 | -1.874 | [-0.111 - 0.008] | 0.000 | 1335 | 0.082 | 0.237 |
| Mean Lateral Ventricle | 0.040 | 0.012 | 1.612 | [-0.013 - 0.092] | 0.000 | 1355 | 0.129 | 0.258 |
| Mean Thalamus | -0.059 | 0.014 | -2.047 | [-0.120 - 0.003] | 0.000 | 1342 | 0.060 | 0.167 |
| Mean Caudate | 0.003 | 0.013 | 0.123 | [-0.054 - 0.061] | 0.000 | 1343 | 0.904 | 0.904 |
| Mean Putamen | -0.029 | 0.015 | -0.974 | [-0.092 - 0.034] | 0.000 | 1341 | 0.347 | 0.462 |

| | | | | | | | | |
|---|---|---|---|---|---|---|---|---|
| Mean Pallidum | 0.016 | 0.015 | 0.529 | [-0.048 - 0.079] | 0.000 | 1335 | 0.605 | 0.692 |
| Mean Hippocampus | -0.062 | 0.014 | -2.126 | [-0.124 - 0.001] | 0.000 | 1335 | 0.052 | 0.167 |
| Mean Amygdala | -0.034 | 0.013 | -1.342 | [-0.088 - 0.020] | 0.000 | 1341 | 0.201 | 0.322 |
| Mean Accumbens | -0.050 | 0.012 | -2.022 | [-0.103 - 0.003] | 0.000 | 1318 | 0.063 | 0.167 |

Note: nominally significant results are italicized and FDR significant results are italicized and bolded

Supplementary Table S21. Associations Between Symptom Correlation Effect Size Maps and Receptor Distribution Maps

| | **SANS Total** | | **MAP** | | **Anhedonia** | | **Asociality** | | **Avolition** | | **EXP** | | **Blunted Affect** | | **Alogia** | |
|---|---|---|---|---|---|---|---|---|---|---|---|---|---|---|---|---|
| | Corr | *p* | Corr | *p* | Corr | *p* | Corr | *p* | Corr | *p* | Corr | *p* | Corr | *p* | Corr | *p* |
| 5HT1a | 0.047 | 0.319 | 0.002 | 0.469 | -0.149 | 0.120 | 0.182 | 0.085 | -0.144 | 0.149 | 0.127 | 0.139 | 0.184 | 0.053 | 0.056 | 0.332 |
| 5HT1b | -0.183 | 0.099 | -0.219 | 0.070 | -0.097 | 0.242 | _-0.273*_ | 0.021 | -0.159 | 0.139 | -0.116 | 0.221 | -0.131 | 0.211 | -0.152 | 0.122 |
| 5HT2a | -0.114 | 0.149 | _-0.222*_ | 0.028 | -0.176 | 0.066 | _-0.300*_ | 0.005 | -0.103 | 0.162 | -0.081 | 0.204 | -0.057 | 0.244 | -0.158 | 0.104 |
| 5HT4 | -0.062 | 0.225 | -0.109 | 0.174 | -0.121 | 0.143 | -0.041 | 0.340 | -0.138 | 0.105 | -0.001 | 0.413 | 0.065 | 0.384 | -0.053 | 0.314 |
| 5HT6 | -0.086 | 0.200 | -0.156 | 0.108 | -0.101 | 0.201 | -0.155 | 0.116 | -0.145 | 0.114 | -0.006 | 0.429 | 0.009 | 0.560 | -0.140 | 0.143 |
| 5HTT | _0.271*_ | 0.018 | _0.311*_ | 0.002 | 0.215 | 0.050 | _0.432*_ | 0.000 | 0.120 | 0.174 | _0.228*_ | 0.044 | _0.227_ | 0.040 | 0.185 | 0.065 |
| A4B2 | _-0.329*_ | 0.008 | _-0.339*_ | 0.009 | _-0.363*_ | 0.006 | _-0.285*_ | 0.019 | _-0.313*_ | 0.012 | _-0.249*_ | 0.042 | _-0.252*_ | 0.037 | -0.169 | 0.101 |
| CB1 | _-0.302*_ | 0.008 | _-0.357*_ | 0.004 | _-0.363*_ | 0.001 | -0.212 | 0.050 | _-0.377*_ | 0.001 | -0.138 | 0.106 | -0.090 | 0.169 | -0.117 | 0.171 |
| D1 | 0.124 | 0.098 | 0.081 | 0.226 | 0.064 | 0.261 | 0.189 | 0.053 | -0.045 | 0.475 | 0.153 | 0.062 | _0.177*_ | 0.038 | 0.017 | 0.450 |
| D2 | -0.063 | 0.318 | -0.110 | 0.192 | -0.163 | 0.107 | 0.041 | 0.371 | -0.210 | 0.053 | 0.074 | 0.254 | 0.134 | 0.129 | 0.016 | 0.461 |
| DAT | _0.233*_ | 0.034 | _0.206*_ | 0.044 | 0.175 | 0.063 | _0.299*_ | 0.007 | 0.042 | 0.311 | _0.275*_ | 0.013 | _0.302*_ | 0.007 | 0.166 | 0.103 |
| GABAa | 0.124 | 0.187 | 0.115 | 0.205 | _0.228*_ | 0.041 | 0.053 | 0.358 | 0.106 | 0.204 | 0.083 | 0.260 | 0.082 | 0.291 | -0.066 | 0.300 |
| H3 | _-0.259*_ | 0.033 | -0.261 | 0.056 | -0.240 | 0.050 | -0.162 | 0.134 | _-0.263*_ | 0.041 | -0.184 | 0.093 | -0.162 | 0.129 | -0.218 | 0.057 |
| M1 | -0.212 | 0.085 | _-0.255*_ | 0.031 | -0.208 | 0.066 | -0.212 | 0.055 | -0.208 | 0.078 | -0.187 | 0.106 | -0.197 | 0.101 | -0.197 | 0.064 |
| mGluR5 | -0.075 | 0.331 | -0.051 | 0.417 | -0.023 | 0.451 | 0.008 | 0.477 | -0.109 | 0.249 | -0.064 | 0.411 | -0.039 | 0.480 | -0.124 | 0.184 |
| MOR | _-0.231*_ | 0.031 | _-0.291*_ | 0.018 | _-0.331*_ | 0.005 | -0.141 | 0.140 | _-0.350*_ | 0.004 | -0.088 | 0.254 | -0.048 | 0.371 | -0.041 | 0.376 |
| NET | -0.162 | 0.141 | -0.093 | 0.267 | -0.048 | 0.393 | -0.065 | 0.335 | -0.119 | 0.222 | -0.163 | 0.135 | -0.184 | 0.097 | -0.122 | 0.169 |
| NMDA | 0.159 | 0.081 | 0.086 | 0.238 | 0.094 | 0.207 | 0.141 | 0.127 | -0.004 | 0.550 | _0.202*_ | 0.036 | _0.208*_ | 0.029 | 0.085 | 0.253 |
| VAChT | 0.007 | 0.442 | 0.073 | 0.306 | 0.065 | 0.305 | 0.170 | 0.116 | -0.056 | 0.368 | 0.011 | 0.427 | 0.023 | 0.368 | -0.014 | 0.467 |

Note: Significant correlations denoted with * and italicized

**Factor Score Formulas**

SANS factor calculation. Factor computation was based on confirmatory factor analyses (Strauss & Ahmed, personal communication, January 6, 2020).

**MAP** = [Recreational Interests and Activities (score)*0.810] + [Sexual Activity (score)*0.641] + [Ability to Feel Intimacy and Closeness (score)*0.688] + [Relationships with Friends and Peers (score)*0.804] + [Grooming and Hygiene (score)*0.314] + [Inpersistence at Work or School (score)*0.553] + [Physical Anergia (score)*0.648]

**Anhedonia Factor** = Recreational Interests and Activities (score)*0.707

**Asociality Factor** = [Sexual Activity (score)*0.685] + [Ability to Feel Intimacy and Closeness (score)*0.736] + [Relationships with Friends and Peers (score)*0.868]

**Avolition Factor** = [Grooming and Hygiene (score)*0.395] + [Inpersistence at Work or School (score)*0.666] + [Physical Anergia (score)*0.807]

**EXP** = [Unchanging Facial Expression (score)*0.831] + [Decreased Spontaneous Movements (score)*0.835] + [Paucity of Expressive Gestures (score)*0.901] + [Poor Eye Contact (score)*0.535] + [Affective Nonresponsivity (score)*0.654] + [Lack of Vocal Inflections (score)*0.765] + [Poverty of Speech (score)*0.775] + [Blocking (score)*0.291] + [Increased Latency of Response (score)*0.415]

**Blunted Affect Factor** = [Unchanging Facial Expression*0.835] + [Decreased Spontaneous Movements (score)*0.840] + [Paucity of Expressive Gestures (score)*0.904] + [Poor Eye Contact (score)*0.541] + [Affective Nonresponsivity (score)*0.658] + [Lack of Vocal Inflections (score)*0.770]

**Alogia Factor** = [Poverty of Speech (score)*0.867] + [Blocking (score)*0.313] + [Increased Latency of Response (score)*0.440]

*Note*. * denotes multiplication.
